# Prefrontal α5-GABA-A receptors: putative role in perceptual learning

**DOI:** 10.64898/2026.03.25.714213

**Authors:** Matthew C. D. Bailey, Emilie Preisler, Clara Velazquez-Sanchez, Lucia Marti Prats, Olivia Stupart, Livia J.F. Wilod Versprille, Johann F. du Hoffmann, Zoe Kourtzi, Jeffrey W. Dalley

## Abstract

Perceptual learning is a dynamic process involving the acquisition and integration of sensory information necessary for adaptive decision making. Resolving the neural basis of perceptual learning may have relevance for our understanding of schizophrenia and other neurodevelopmental disorders that implicate impaired perceptual acuity. We developed a novel touchscreen task that utilizes orientation discrimination and variable attentional load to elucidate the perceptual and attentional components of visual perceptual learning (VPL) in male and female rats. Based on previous evidence we hypothesised that individual variation in VPL may depend in part on inhibitory neurotransmission mediated by γ-amino butyric acid (GABA). Segregating subjects based on ‘poor’ and ‘good’ learning after task pretraining revealed dose-dependent improvements in VPL in poor learners following the administration of an α5 subunit specific GABA-A (GABRA5) positive allosteric modulator (Alogabat) and a GABA-B agonist (R-baclofen) early in learning. Poor VPL performance was associated with significantly reduced GABRA5 mRNA expression in dorsal regions of the prefrontal cortex (PFC), notably the prelimbic cortex. Decreased GABRA5 expression in this region was co-localized to somatostatin-and parvalbumin-expressing interneurons. These findings reveal a potential link between GABRA5 expression in selective PFC populations of inhibitory interneurons and the speed and acuity of VPL.

---

Visual perceptual learning (VPL) refers to long-term improvements in visual discrimination that emerge after repeated training. Previous studies have revealed finely tuned stimulus- and context-specific cortical responses during VPL (Ball and Sekuler 1987; Karni and Sagi 1991; Shiu and Pashler 1992; Ahissar and Hochstein 1997). Such responses are linked to synaptic plasticity in primary visual cortex (V1) with intrinsic neurons exhibiting small receptive fields and narrow feature selectivity (Hubel and Wiesel 1962; Hubel and Wiesel 1968; Schoups et al. 2001). Higher visual processing areas including parietal and frontal cortices involved in decision-making and attention are also progressively recruited alongside V1 and the superior colliculus to refine behavioural output and improve visual discrimination (Mukai et al. 2007; Kahnt et al. 2011; Jing et al. 2021). However, the relative contributions and balance between bottom-up sensory *versus* top-down decisional and attentional control processes in VPL remain poorly understood.

Inhibitory control is fundamental to reshaping cortical circuits throughout development and adulthood, coordinating plasticity during sensitive periods (Ganguly et al. 2001; Kirmse et al. 2015) and VPL (Froemke 2015; Tremblay et al. 2016; van Versendaal and Levelt 2016; Desrosiers and Basha 2023). γ-Aminobutyric acid (GABA), the primary inhibitory neurotransmitter in the brain acts in concert with glutamate, the primary excitatory neurotransmitter, to regulate local circuit and global network excitability *via* the excitatory-inhibitory balance (E/I). Dysregulated E/I balance is associated with perceptual deficits in neurodevelopmental disorders such as autism (Cooper et al. 2017; Hollestein et al. 2023) and schizophrenia (Lesh et al. 2011; McCutcheon et al. 2023; Howes et al. 2024), with aberrant inhibition driving network abnormalities (Hoftman et al. 2015; Ferguson and Gao 2018; Mueller-Buehl et al. 2023; van Hugte et al. 2023; Kaar et al. 2024; Marín 2024). Resolving the mechanism by which inhibition contributes to VPL may thus inform the development of new interventions to improve perceptual learning in a variety of disorders.

At the cortical circuit level, distinct populations of inhibitory interneurons contribute to experience-dependent remodelling of sensory, associative and executive networks recruited during learning and attentional tasks (Yazaki-Sugiyama et al. 2009; Kato et al. 2015; Kuchibhotla et al. 2017; Poort et al. 2022). These include parvalbumin (Pvalb)-, somatostatin (Sst)-, and vasoactive intestinal peptide (VIP)-expressing neurons in V1, which dynamically remap their activity as VPL proceeds (Kato et al. 2015; Khan et al. 2018; Aime et al. 2022; Myers-Joseph et al. 2024; Cheng et al. 2025). During late stages of learning, increased attentional engagement recruits higher-order inhibitory microcircuits to sharpen stimulus selectivity and enhance network fidelity (Snyder et al. 2016; Speed et al. 2020; Raven and Aton 2021; Snyder et al. 2021). However, within these microcircuits, GABAergic interneurons exhibit significant heterogeneity in expressed receptors and receptor composition and are recruited in a context dependent manner to modify network excitability (Olsen and Sieghart 2009; Kohl and Paulsen 2010; Pinard et al. 2010; Weir 2016).

Given the established role of the α5 subunit containing GABA-A receptor (GABRA5) in hippocampal-dependent learning and memory (Caraiscos et al. 2004; Zarnowska et al. 2009; Prut et al. 2010; Engin et al. 2015), the present study investigated the potential of GABRA5 to enhance VPL. Although GABRA5 represent less than 5% of cortically expressed GABA-A receptors, they are expressed both intra- and extra-synaptically and thus contribute respectively to phasic and tonic inhibition (Caraiscos et al. 2004; Serwanski et al. 2006; Panzanelli et al. 2011). GABRA5 receptors also show dynamic activity-dependent synaptic re-localisation, a process hypothesised to facilitate fine tuning of tonic and phasic inhibitory tone to preserve learned associations in a context dependent manner (Davenport et al. 2021). Thus, modulation of GABRA5 receptors may offer a promising strategy to rescue or enhance VPL in autism, schizophrenia and other neurodevelopmental disorders.

To investigate the role of GABRA5 in VPL we established a novel touchscreen task for rats involving the discrimination of visual gratings with distinct orientations, paired with an additional “blank” distractor to separate perceptual and attentional components of VPL. We compared sex-differences in task acquisition and baseline performance before testing the sensitivity of the task to a range of GABAergic compounds that affect distinct aspects of GABAergic synaptic transmission and stratified these effects according to poor and good learning performance on the task. Given that the rate of learning in visual touchscreen discrimination tasks is linked to thresholds of visual acuity (Crijns and Op de Beeck 2019) we also investigated the dimensional relationship of VPL in individual animals and expression levels of the GABRA5 mRNA transcript and protein product in V1 and frontal cortex. We hypothesised that enhancing GABAergic inhibition with selective GABA reuptake inhibition (tiagabine) and a positive allosteric modulator of GABRA5 (alogabat) (Cecere et al. 2025) would enhance VPL. We further hypothesised that inter-individual variation in VPL would relate to GABRA5 expression with faster VPL in animals exhibiting increased GABRA5 expression.

## 2.0 Materials and Methods

### 2.1 Subjects

Thirty-two Lister-hooded rats (16 male, 16 female) were raised from time mated dams (Envigo, Bicester, UK), weaned at post-natal-day (PND) 21, and housed in groups of four with environmental enrichment under a 12/12hr reverse light/dark cycle (lights on 6pm). From weaning, animals had *ad libitum* access to food (Safe A40, Germany) and water. The temperature of the holding room was maintained between 20-22°C with a relative humidity of 55 ± 3%. After PND26, animals were handled three times each week to habituate them to human contact. On PND56 animals were food restricted to no less than 85% free feeding weight for the remainder of the experiment and weighed at least once per week. Water access was not restricted. For the first experiment, larger male rats were shifted to pairwise housing to maintain adequate space in the holding cages. All procedures were carried out in accordance with the UK (1986) Animal (Scientific Procedures) Act (PPL PP9536688) and were approved by the University of Cambridge Animal Welfare and Ethical Review Body (AWERB).

### 2.2 Visual perceptual learning task

Two rooms, each containing 8 identical operant chambers in a light/sound attenuating and fan-ventilated box were used for behavioural testing. One was used for male subjects, the other for female subjects. Chambers running K-Limbic operant control software (Med Associates, UK) were integrated with a 1024x768 resolution portrait touchscreen (Nexio, UK) on one wall, and a food magazine with light, pellet dispenser, house light, and tone generator on the opposite wall (see **Figure 1A**). A camera was installed overhead to record behaviour.

**Figure 1:**
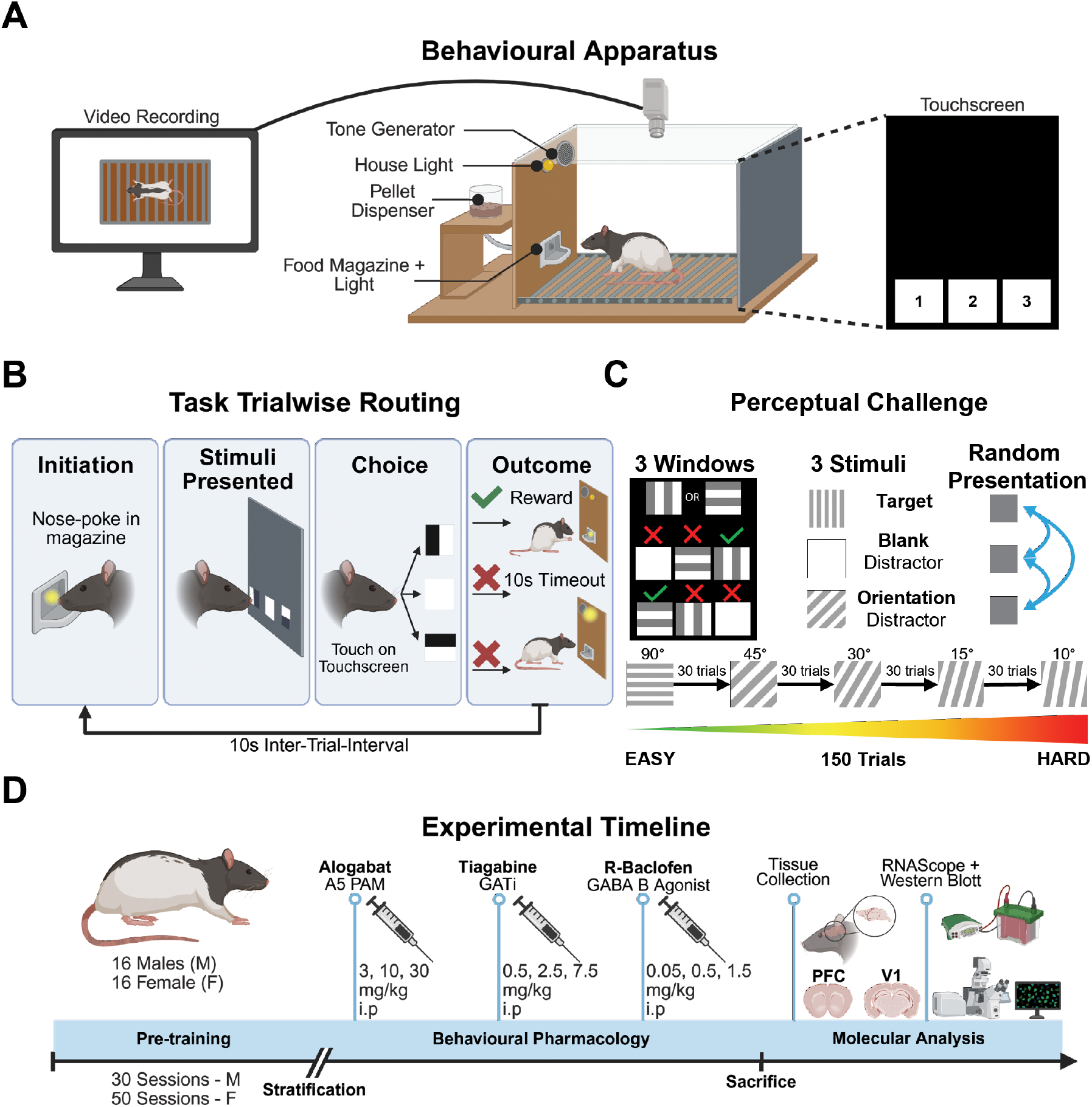
Visual discrimination task and experimental timeline. **A** Operant chamber apparatus. Med Associates chambers showing the position of the house-light, tone generator and food magazine with a light connected to a pellet dispenser on one side of the chamber and a 1024 x 768 resolution touchscreen on the other. A CCTV camera was mounted overhead to record behaviour. **B** Trial wise decomposition of final task contingencies. A nose poke in the illuminated food magazine initiated a trial. Stimuli were then presented on the touchscreen (1 target grating, 2 distractors (1 blank, 1 grating) with animals making a choice between the 3 presented stimuli. A correct response was reinforced with a sugar pellet paired with tone (magazine light on during collection). Incorrect responses at either the distractor or background stimuli received a 10 second timeout with no reward and house light illumination. **C** Baseline task perceptual challenge and distractor presentation sequence. In total, the task consisted of 150 trials in blocks of 30 at each distractor frequency, starting at 90° from target. Stimuli were presented randomly on a trial-by-trial basis. **D** Experimental timeline showing behavioural stratification, pharmacological challenges and molecular analyses. Figure created using https://app.biorender.com/

To investigate VPL, a novel 3-stimuli orientation-based visual discrimination paradigm adapted from Horner and colleagues (Horner et al. 2013) was developed using sinusoidal stimuli presented in three equally spaced response windows (200x200 pixels), located 150 pixels from the bottom of the touchscreen. Target stimuli were assigned at the beginning of training, counterbalanced within the cohort (16 animals 0°, 16 animals 90°) and spatially randomised on a trial-by-trial basis to reduce procedural learning and location bias.

#### 2.2.1 Pretraining

Pretraining (PT) proceeded in 4 stages (PT1-4): The purpose of PT1 was to acclimatise animals to the operant chamber and reward pellets (TestDiet AIN-76, 5TUL, Cat No.1811155). PT1 involved placing the animals in the chamber for 30 minutes with 10-20 reward pellets in the food magazine. Following 1 session of PT1 all animals were moved to PT2.

PT2 involved appetitive autoshaping to the touchscreen with reward, and variable locations of the target stimulus. PT2 sessions included 90 trials/session of auto-initiated presentation of the single counterbalanced target stimuli in any of the 3 response windows on the touch screen. Nose poking the target was rewarded with a single pellet in the magazine paired with magazine light and tone as visual and auditory reinforcers followed by 2 second “eating time” and 10 second inter-trial-interval (ITI). There was no stimulus presentation timeout or timeout for touching elsewhere on the touchscreen. Target stimulus window presentation varied on a trial-by-trial basis. Animals were deemed to have acquired this association after 2 consecutive sessions with 80% touches (72/90) completed, after which they progressed to PT3.

PT3 focused the reward association to the specific target sinusoid stimulus and task-related attention between the magazine and the touchscreen on a trial-by-trial basis, introducing a timeout for background touches and self-initiation of trials via food magazine nose poke. PT3 consisted of 90 self-initiated trials/session where animals were required to nose poke into the illuminated food magazine (no pellet present).

The target sinusoid was then presented individually on the touchscreen as in PT2 but now touches on the black touchscreen background were punished with a 10 second timeout signalled by 10 second sustained flash of the house light before the ITI. Trials were then reinitiated via poking in the lit magazine. There was no timeout on stimuli response presentation. As in PT2, animals were deemed to have made this association after 2 consecutive sessions with 80% touches (72/90) completed, after which they were progressed to PT4.

PT4 involved a discrimination between 3 simultaneously presented stimuli. Here, in addition to the target sinusoid, a pair of blank stimuli were simultaneously presented on each trial with the target as white squares of identical dimensions. Touches at the target stimuli were rewarded whilst touches at either one of the blank stimuli or the touchscreen background were punished with a timeout period as before. The spatial arrangement of targets and distractors was varied on a trial-by-trial basis within the 3 touchscreen response windows to reduce window/side bias. Sessions now included 120 trials. The criterion for progression was >80% correct on 2 consecutive sessions (see **Figure S1**).

#### 2.2.2 VD training

The efficacy of the chosen distractor was tested during a series of sessions involving the discrimination of single distractor orientations. For animals that reached criterion on PT4, 5 distractor orientations (measured in degrees from target orientation) were tested in VD1-5 sessions: VD1 = 90°, VD2 = 45°, VD3 = 30°, VD4 = 15°, VD5 = 10°.

VD training sessions consisted of 150 trials of target sinusoid, orientation distractor (fixed per session), and blank distractor stimuli presentation. Reward and punishment remained unchanged from previous training stages, touches at orientation distractor had the same punishment as other errors. Session-wise progression from easy to hard distractors was accuracy based with 2 consecutive sessions at criteria: >80% VD1 → VD2, >70% VD2 → VD3, >60% VD3→VD4, >50% VD4 →VD5, >50% VD5 → baseline task. If animals could progress through these stages and showed a decreasing accuracy trend, the angular separations were considered appropriate.

Animals that reached VD5 but failed to achieve >50% accuracy in 5 sessions were deemed to have reached their perceptual limit and were moved to the baseline task to prevent unlearning of their target sinusoid reward association.

It should be noted that 5 male animals achieved this criteria, and the remainder of the male cohort was moved to the final task after 33 training sessions to limit unequal exposure to the baseline task. At the same point in the study, no female animals had reached this threshold, with most still in PT3/4. To account for this variation, females were kept on pretraining for longer and moved to baseline after 52 sessions. This point was selected due to plateau in performance, a demonstration that a subset of female animals could reach VD training (n = 4), and to match the timing of the Latin square pharmacology study in males.

#### 2.2.3 Baseline (final task)

The final task involved 150 trials split into five, 30-trial blocks of distractor orientation which converged on the target stimulus orientation (Easy -90°, 45°, 30°, 15° and 10° -Hard). On any given trial, three stimuli were presented, one target grating, one distractor grating, and one white box stimulus randomised on a trial-by-trial basis. As in pretraining, animals were required to initiate trials via poking in the lit food magazine, before responding to presented stimuli on the touchscreen. Responses at the target stimulus were rewarded with a sucrose pellet in the lit food magazine and auditory tone whereas incorrect responses at either the distractor stimuli (distractor grating or white) or background were punished with a 10s house light pulse timeout and no food reward. Sessions lasted up to 1 hour with an inter-trial-interval (ITI) of 10s and were completed once per day.

Additionally, target distractor distance (TDD) was classified as ‘close’ when target grating and distractor grating were presented in adjacent windows, and ‘far’ when target distractor gratings were separated by the third, white stimuli. TDD was employed to independently investigate the role of attentional load and simultaneous suppression of multiple stimuli on perceptual decision making. (see **Figure 1B, C**). For further details of the training stages and task contingencies refer to **Figure S1**.

### 2.3 Subject stratification

To investigate individual differences in visual perceptual learning, animals were stratified into ‘learning type’ groups based on their task acquisition rate. For males (n=16), animals that completed pretraining and VD training during the thirty-three-session training period were classified as “good learners” (n=5), whereas animals that failed to reach visual discrimination training were classified as “poor learners” (n=6). Animals that completed pretraining but did not finish VD training were classed as “intermediate learners” (n=5) and represented the mean task acquisition rate.

Although females were included in the study, it was not possible to stratify them in the same way as males because considerably more sessions were required for females to acquire the task (mean 52 sessions). Moreover, only some of the females completed the more difficult VD training stages of the VPL task, as shown in **Figure S2**, meaning that based on the criteria applied to the males, no females were classed as ‘good learners’ despite the extended training period. Instead, females were classed as ‘poor’ (n=8), or, for animals which either reached criterion on the PT4 at least once or began VD training, ‘intermediate’ (n=8). These groups represent an equal proficiency-based stratification, applied after greater training, and were used in comparing within-sex performance, pharmacology and molecular variations. Details of all female analysis can be found in the supplementary material section.

### 2.4 Pharmacological interventions

Selective GABAergic compounds were administered according to a Latin square design. The compounds (and doses) were tested in with a 3-day dosing schedule (day 1 (dose level 1), day 2 (no drug), day 3 (baseline) and at least 3 day washout period between each compound: (1) alogabat (3, 10, 30 mg/kg) (2) tiagabine hydrochloride (0.5, 2.5, 7.5 mg/kg); (3) R-baclofen (0.05, 0.5, 1.5 mg/kg). Alogabat (provided by Boehringer Ingelheim Pharma, Germany) was administered in 5% HCl (1M) and 95% 2-(hydroxypropyl)-β-cyclodextrin (20%) at 2 ml/kg (pH 7-7.5). Tiagabine-HCl (Insight Biotechnology, UK-CAS 145821-59-6) was administered in deionised water pH 7-7.5 at 1 ml/kg. R-Baclofen (Insight Biotechnology, UK-CAS 69308-37-8) was administered in 0.9% saline (pH 7-7.5) at 1 ml/kg. Tiagabine and R-baclofen were administered simultaneously to male and female animals, whilst the Latin square for alogabat was split, and administered first to males, and last for females, accounting for the time difference in pretraining. This resulted in differences between sexes in animal baseline task exposure: specifically, alogabat (sessions 1-13 male, 16-23 female), and alogabat administration in males (sessions: 1-8 poor, 5-13 good). All drugs were administered *via* intraperitoneal (i.p.) injection with 30-minute pretreatment time (See **Figure 1D** and **Figure S10** for further details).

### 2.5 Tissue collection

Rats were anesthetised with isoflurane (Sigma-Aldrich, UK) *via* inhalation before decapitation 5 days after the final VPL session. Brains were collected and snap-frozen with dry-ice pre-cooled isopentane (Sigma-Aldrich, UK) and stored at -80°C. Brains were sectioned using a cryostat (Leica, UK) to obtain coronal slices of the medial prefrontal cortex (mPFC) (5.16 to 2.28 mm from Bregma) and V1 (-5.6 to -9.1 mm from Bregma). Each hemisphere was prepared for Western blotting (WB) with sections cut at 200 μm or RNAscope *in situ* hybridisation (RNAScope) with sections cut at 12 μm. Hemispheres were counterbalanced between the two assays, and slices were mounted on Menzel gläzer superfrost® slides (Thermo Fisher Scientific, UK) for WB or Superfrost® plus slides (Thermo Fisher Scientific, UK) for RNAscope. Sections were stored at -80°C prior to processing.

### 2.6 Western immunoblot analysis

WB were conducted using a WB kit according to manufacturer guidelines (Invitrogen-Life Technologies, UK). Four regions-of-interest (ROI)s were extracted from slices of each rat: V1, Cg1, PrL, and IL. After immunoblotting, GABRA5 expression was developed using a primary antibody for GABRA5 (Biorbyt, UK – 1:1000 Cat No. orb374299) incubated overnight at 4°C on a shaker. The primary antibody for the housekeeping protein actin (Abcam, UK – 1:10,000 Cat No. ab8226) was incubated for 1h at room temperature. Secondary antibodies, IRDye680RD and IRDye800CW (LI-COR, UK – 1:15,000 Cat No: 680, 926-68071; 800, 926-32210) were incubated at room temperature for 1h alongside a chameleon duo ladder (LI-COR, UK – Cat No. 928-60000) before imaging (Odyssey Imaging System, LI-COR, UK).

### 2.7 RNAscope fluorescence in-situ hybridisation

The spatial transcriptomic profile of GABRA5 in the mPFC and V1 was assayed in replicate samples to investigate cell-type specific GABRA5 mRNA expression. The assay and target probes were performed (Wang et al. 2012) according to the fresh frozen tissue protocol provided by the manufacturer (RNAscope fluorescent multiplex reagent kit, ACD, UK) with specific details given in supplementary **Table 1**. Positive and negative control probes were used to validate the assay in each of the 4 runs.

Tissue was imaged using a Zeiss Axio Imager M2 microscope with an AxioCam MRm microscope camera (Oberkochen, Germany) running Visiopharm software (Medicon Valley, Denmark). Approximately 15 images were each taken of V1 and mPFC subregions (cingulate cortex -Cg1; prelimbic cortex – PrL; infralimbic cortex -IL) with 20X magnification. Imaging settings such as exposure time and light source power were controlled between assays of each region (V1, PFC) to facilitate within region comparison of expression profiles.

Signals were quantified using an RNAscope signal analysis pipeline developed by Velazquez-Sanchez et al. (Velazquez-Sanchez et al. 2023), which utilised a custom MATLAB script (The MathWorks Inc) combined with a FIJI/ImageJ (Schindelin et al. 2012) stardist machine learning algorithm to count mRNA molecules within DAPI labelled cells. Briefly, images were pre-processed to remove background, before running stardist segmentation on DAPI channel images. Segmented DAPI channel images were then merged with mRNA fluorescent channels and run through the MATLAB script which applies a threshold to each channel based on percentage max fluorescence to identify and quantify signal intensity via puncta counts. Signals were associated with segmented DAPI signals if within 10 pixels radius of the edge of the nucleus. Thresholds were held constant between assays with region to allow comparison between cell markers equally. Output figures were inspected to identify and remove outlier images. A Python script was then used to concatenate outputs where cells co-expressed GABRA5 with inhibitory cell markers of interest, and presented by cell number, and number of mRNA copies.

### 2.8 Statistical analysis

Behavioural data were collected on a trial-by-trial basis and pre-processed to remove outliers. Baseline outliers were removed on the basis on response latencies that were too fast (<100ms) to be instrumental in nature (e.g. arising from unintentional tail contacts with the touchscreen). For the pharmacological experiments, outliers were removed for <96/150 trials completed in each session – only reaching the third block of distractor, failed/incomplete dosing, or a failure to reach the trial completion criteria on <2/4 doses of each compound (n=1 removed per compound), final n: n=5 poor learners, n=5 good learners. Reaction time responses were inverse transformed (-100/RT) to normalise the distribution of the data and facilitate more robust modelling.

Statistical comparisons were made in R (Version 4.4.2)(R: The R Project for Statistical Computing 2026) using within-subjects mixed-effects modelling, a generalised logit link function to quantify trial wise accuracy, and a linear model to analyse inverse transformed mean response latencies (inverseRT). Fixed-effects relationships between angular separation from target (ang_sep), sex, learning type (LT) and target distractor distance (TDD) were investigated. Animal ID was added as a random effect with a random intercept to account for within group individual variation (see **Figure S3**). Pharmacological models were similar, although due to within-sex comparisons only, model selection showed TDD did not have significant explanatory power and was therefore not included in the final model. Ang_sep was also only included as a fixed effect rather than interaction due to model convergence violations. Where appropriate, estimated-marginal-means pairwise comparisons with Tukey HSD correction were performed and reported as odds ratios for generalised models.

A within-subjects linear mixed-effects model was applied to molecular data to investigate the relationships between learning-type and GABRA5 protein and mRNA expression within the various brain regions and specified cell types of interest, respectively. Each animal was added as a random effect with a random intercept to account for within group individual variation. An ANOVA was performed to identify main effects, and where appropriate, estimated-marginal-means pairwise comparisons with Tukey HSD correction were performed between groups within region, within cell type, and adjusted p (adj.p) values ≤ 0.05 reported (see **Figure S4**). All variables and models were performed and evaluated for analysis suitability using Shapiro-wilks normality and model evaluation metrics available in the Performance package in R which involved likelihood ratio testing, and model goodness of fit plots including q-q, homogeneity of variance, and collinearity (see **Figure S3**). Of note, for more robust modelling GABRA5:actin was square root transformed for male derived samples, while for female derived samples, RNAScope outputs were log_10_ transformed. Confidence intervals are reported because power analyses are estimates for mixed-effects model structures. Full summaries and outputs of statistical models can be found in supplementary **Table 2**.

## 3.0 Results

### 3.1 Modelling VPL using a novel orientation discrimination task

Assessment of performance on the baseline version of the VPL task revealed that narrowing the angular separation from the target stimulus decreased discriminative accuracy (**Figure 2A**) (relative to ang_sep90: ang_sep45: odds ratio = 1, 95% CI [0.91, 1.09], p = 0.9703, ang_sep30: odds ratio = 0.83, 95% CI [0.76, 0.9], p < 0.001, ang_sep15: odds ratio = 0.55, 95% CI [0.5, 0.6], p < 2e^-16^, ang_sep10, odds ratio = 0.51, 95% CI [0.46, 0.56], p < 2e^-16^). Here there was also a within-block effect of sex on trial-wise mean accuracy with females displaying lower accuracy from block 2 onwards (ang_sep90, F-M: odds ratio = 0.87, 95% CI [0.61, 1.22], adj.p = 0.4181, ang_sep45, F-M: odds ratio = 0.7, 95% CI [0.5, 0.99], adj.p = 0.0471, ang_sep30, F-M odds ratio = 0.71, 95% CI [0.5, 1], adj.p = 0.0531, ang_sep15, F-M: odds ratio = 0.68, 95% CI [0.48, 0.96], adj.p = 0.0275, ang_sep10, F-M: odds ratio = 0.67, 95% CI [0.47, 0.95], adj.p = 0.0247). However, the accuracy of males and females across all blocks was not significantly different (comparison F: M: odds ratio = 1.06, 95% CI [0.75, 1.5], p = 0.732) (**Figure 2B**).

**Figure 2:**
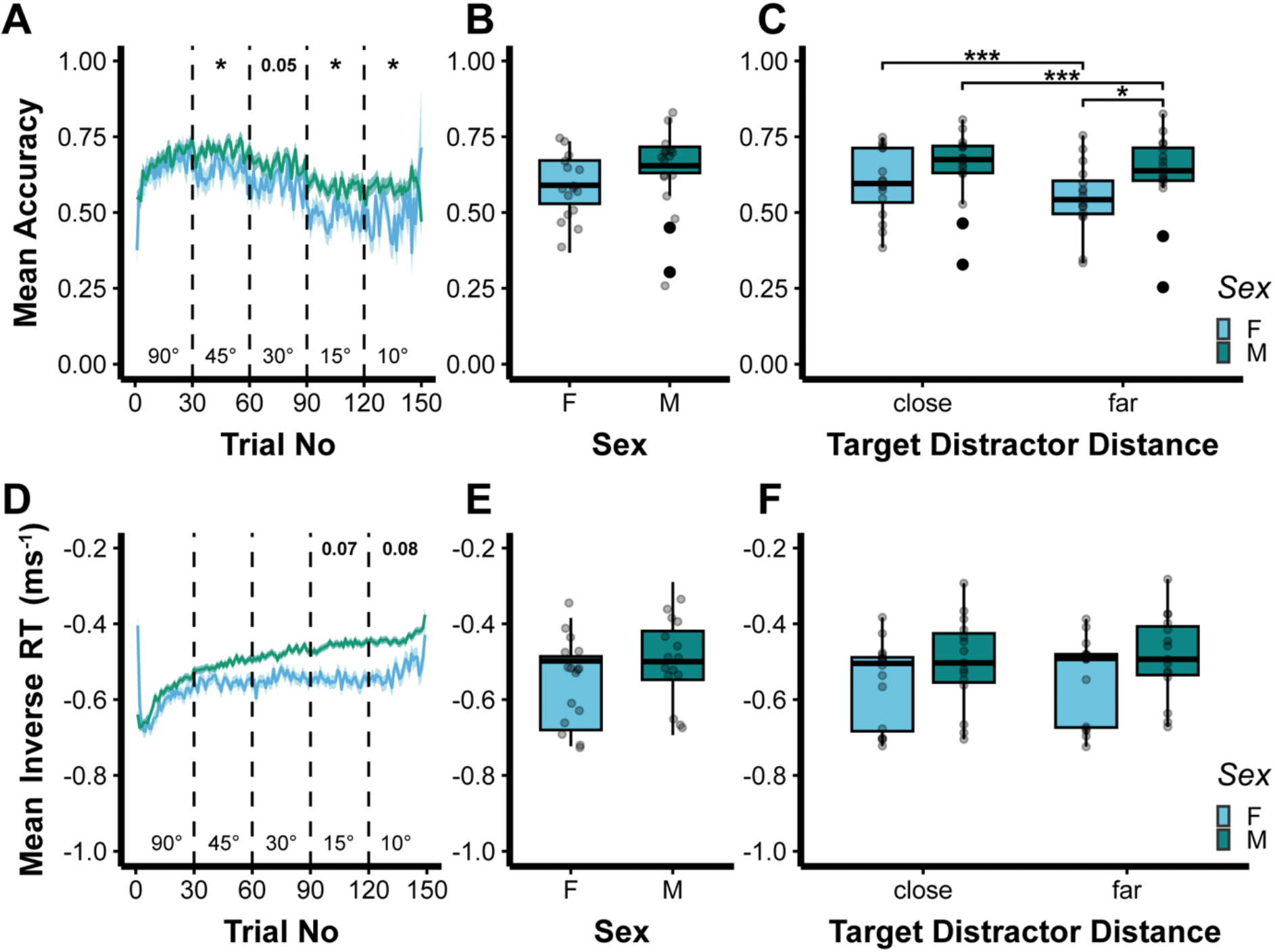
Baseline performance measures on the VPL task. **A** Trial-wise mean accuracy by sex presented as mean ± SEM. Dashed lines denote distractor frequency blocks. Annotations denote between-sex comparisons within each distractor frequency block. **B** Distribution of baseline task accuracy segregated by sex. Each data point denotes an individual animal. **C** Distribution of mean task accuracy for each target distractor distance (TDD). Comparisons made within TDD-between sex and within sex-between TDD. **D** Trial-wise mean inverseRT by sex presented as mean ± SEM. Dashed lines mark distractor frequency blocks.

When the relative location of the target and distractor gratings was considered (Target Distractor Distance – TDD), both sexes were impaired by a greater separation distance between stimuli (F, TDDclose – TDDfar: odds ratio = 1.27, 95% CI [1.2, 1.35], adj.p < 0.001. M, TDDclose – TDDfar: odds ratio = 1.08, 95% CI [1.04, 1.12], adj.p < 0.001), with females showing a more pronounced reduction in accuracy than males (TDDfar, F-M: odds ratio = 0.67, 95% CI [0.47, 0.94], adj.p = 0.0199) (**Figure 2C**). There was no such effect with close TDD (TDDclose, F-M: odds ratio = 0.79, 95% CI [0.56, 1.1], adj.p = 0.1629). Analysis of concordant inverse transformed response latencies (inverseRT) revealed a slowing of responses with a narrowing of angular separation (**Figure 2D**) (F(_4,269_) = 123, Pr(>|z|) < 2.2e^-16^) and a blockwise effect of sex (ang_sep90, F-M: difference = -0.01, 95% CI [-0.09, 0.07], adj.p = 0.8477, ang_sep45, F-M: difference = -0.04, 95% CI [-0.12, 0.04], adj.p = 0.3632, ang_sep30, F-M: difference = -0.05, 95% CI [-0.14, 0.03], adj.p = 0.2043, ang_sep15, F-M: difference = -0.08, 95% CI [-0.16, 0], adj.p = 0.0696, ang_sep10, F-M: difference = -0.08, 95% CI [-0.16, 0.01], adj.p = 0.0781 (**Figure 2E**). However, there was no significant difference between males and females on inverse RT across the full task (F(_1, 29_) = 1.54, Pr(>F) 0.225) or when analysed by TDD (Sex:TDD interaction, F(_1, 269_) = 0.224, Pr(>F) = 0.637 (**Figure 2F**).

Annotations denote between-sex comparisons, within distractor frequency block. **E** Distribution of mean inverseRT segregated by sex. Each data point denotes an individual animal **F** Distribution of mean inverseRT by TDD. Comparisons made within TDD-between sex and within sex-between TDD. Baseline sessions displayed for each sex with statistical comparisons made by generalized linear mixed-effects model with binomial link for accuracy (**A-C).** Linear mixed-effects model for inverseRT (**D-F)**. Boxplots denote median, interquartile range with whiskers extending to 1.5x IQR. Outliers denoted by solid points. Pairwise comparisons made via estimated-marginal-means to “Female” condition in each case, within block **(A,D)** and “close” **(C,F)**. Effect sizes: “p value” = p<0.1, “*” = p<0.05, “***” = p<0.001 **(D-F)** as log-odds ratios **(A-C)**.

### 3.2 Segregation of good *versus* poor learners on the VPL task

Given that difficult learning paradigms require extensive training and that subjects are often over trained by the time they are evaluated at baseline, we segregated animals during the task acquisition stage. To achieve this, we grouped subjects by their final training stage before the drug challenges. Those animals that completed the pretraining stages, reaching the baseline task were classed as “Good Learners” (n = 5); those which did not complete pretraining were classed as “Poor Learners” (n = 6) (**Figure 3A**). Due to differing pre-training exposures, we could not directly compare sexes with this stratification approach. Instead, we completed a concordant stratification analysis for female animals available in the supplementary materials (**Figures S7, S8, S9**).

**Figure 3:**
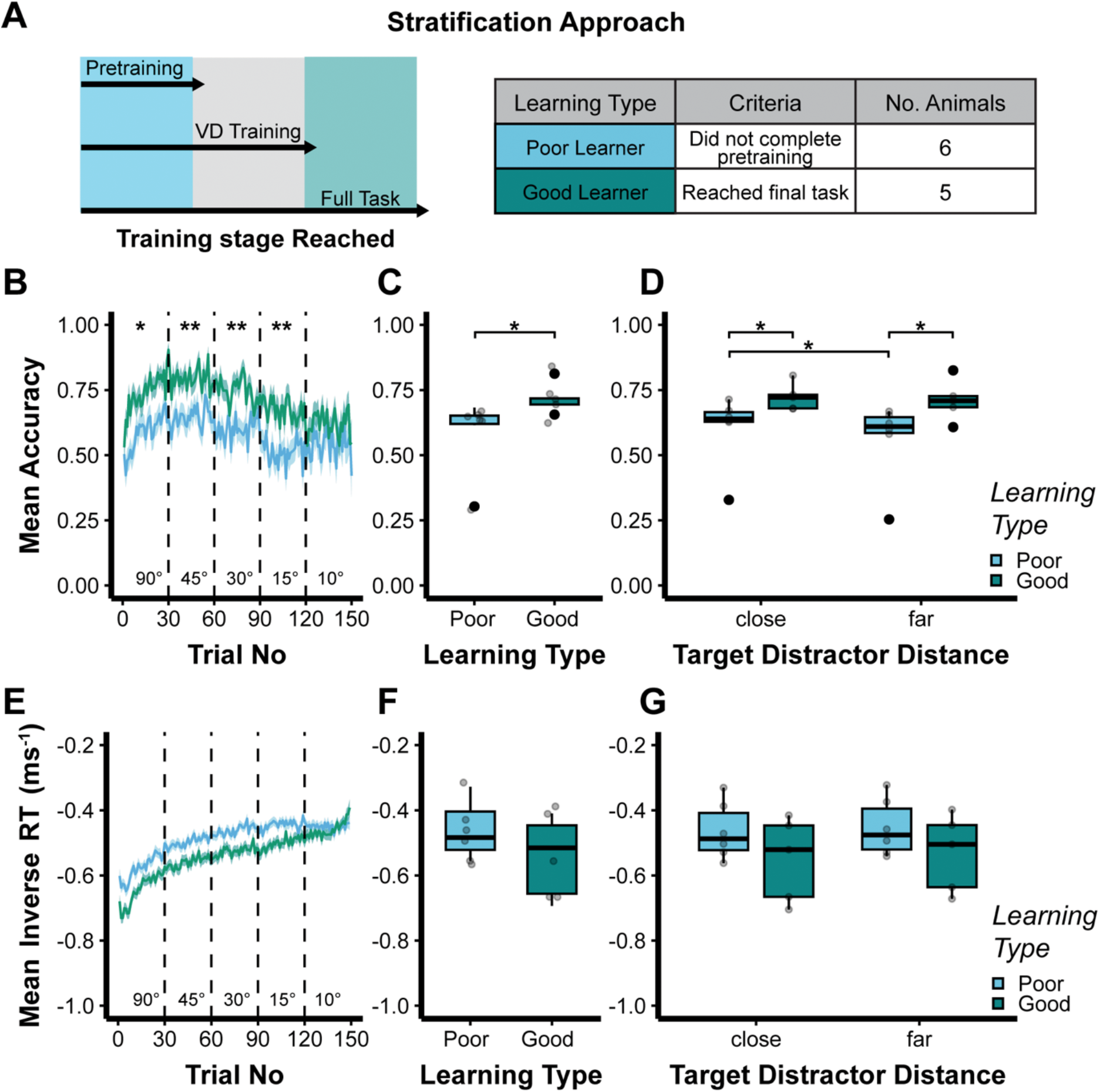
Summary measures of perceptual performance on the VPL task. **A** Summary of stratification procedure based on training stage reached at the end of 33 sessions of pretraining (males only). **B** Trial-wise mean accuracy by learning type presented as mean ± SEM. Dashed lines mark distractor frequency blocks. Annotations denote between-learning type comparisons, within distractor frequency block. **C** Distribution of baseline accuracy by learning type. **D** Distribution of mean accuracy segregated by target distractor distance. Points denote individual animals per TDD. Comparisons made within TDD-between learning type and within learning type-between TDD. **E** Trial-wise mean inverseRT by learning type presented as mean ± SEM. Dashed lines mark distractor frequency blocks. Annotations denote between-learning type comparisons, within distractor frequency block. **F** Distribution of mean inverseRT by learning type. **G** Distribution of mean inverseRT by target distractor distance. Points denote individual animals per TDD. Comparisons made within TDD-between learning type and within learning type-between TDD. Baseline sessions are displayed for each learning type with statistical comparisons made by generalized linear mixed-effects model with binomial link for accuracy (**B-D).** Linear mixed-effects model for inverseRT (**E-G)**. Boxplots denote median, interquartile range with whiskers extending to 1.5x IQR. Outliers denoted by solid points. Pairwise comparisons made via estimated-marginal-means to “Poor learner” condition in each case, within block **(B,E)** and “close” **(D,G)**. Effect sizes: “*” = p<0.05, “**” = p<0.01**(E-G)** as log-odds ratios **(B-D)**.

In males, groupwise comparisons revealed within-block differences in learning accuracy (**Figure 3B**) where good learners were more accurate than poor learners in all but the final block (ang_sep90, poor-good: odds ratio = 0.52, 95% CI [0.31, 0.89], adj.p = 0.0166, ang_sep45, poor-good: odds ratio = 0.47, 95% CI [0.28, 0.8], adj.p = 0.0057, ang_sep30, poor-good: odds ratio = 0.49, 95% CI [0.29, 0.83], adj.p = 0.0083, ang_sep15, poor-good: odds ratio = 0.5, 95% CI [0.29, 0.84], adj.p = 0.0095, ang_sep10, poor-good: odds ratio = 0.69, 95% CI [0.4, 1.17], adj.p = 0.1688). This was associated with good learners displaying a greater full task accuracy (**Figure 3C**) (poor – good: odds ratio = 1.84, 95% CI [1.08, 3.13], p = 0.0253) and both TDD close and far accuracies than poor learners (**Figure 3D**) (TDDclose, Poor-Good: odds ratio = 0.55, 95% CI [0.33, 0.93], adj.p = 0.0258, TDDfar, Poor-Good: odds ratio = 0.51, 95% CI [0.3, 0.86], adj.p = 0.0114). Additionally, whilst poor learner accuracy was negatively impacted by increased TDD, good learners displayed only a marginal reduction in performance between TDDclose and TDDfar trials (Poor, TDDclose – TDDfar: odds ratio = 1.17, 95% CI [1.09, 1.25], adj.p < 0.001, Good, TDDclose – TDDfar: odds ratio = 1.07, 95% CI [1, 1.15], adj.p = 0.0599).

Analysis of inverse RT highlighted no block-wise differences between learning types (**Figure 3E**) (ang_sep90, Poor-Good: difference = 0.08, 95% CI [-0.05, 0.22], adj.p = 0.2625, ang_sep45, Poor-Good: difference = 0.09, 95% CI [-0.04, 0.23], adj.p = 0.2027, ang_sep30, Poor-Good: difference = 0.1, 95% CI [-0.03, 0.24], adj.p = 0.1699, ang_sep15, Poor-Good: difference = 0.09, 95% CI [-0.04, 0.23], adj.p = 0.2044, ang_sep10, Poor-Good: difference = 0.08, 95% CI [-0.05, 0.22], adj.p = 0.2537) or task-wise differences between learning types (**Figure 3F**) (poor – good: F(_1, 9_) = 1.81, Pr(>F) = 0.212. There were also no within or between group effects of TDD on RT (**Figure 3G**) (F(_1, 89_) = 0.558, Pr(>F) = 0.457).

### 3.3 Pharmacological interventions

To investigate the contribution of GABRA5 to VPL we administered the GABRA5 positive allosteric modulator (PAM) alogabat during task performance. We also compared the effects of alogabat with tiagabine, a GABA reuptake inhibitor and baclofen, a GABA B agonist.

As shown in **Figure 4A**, tiagabine had no significant effect on accuracy (compared to Veh: 0.5 mg/kg, odds ratio = 1.03, 95% CI [0.79, 1.33], p = 0.8307, 2.5 mg/kg, odds ratio = 0.91, 95% CI [0.71, 1.17], p = 0.4605, 7.5 mg/kg, odds ratio = 0.99, 95% CI [0.77, 1.28], p = 0.961). However, in good learners, tiagabine exhibited a dose dependent effect on response latencies (main effect of dose: F(_3,161_) = 17.280, Pr(>F) = 8.88e^-10^). Here, low (Veh-0.5 mg/kg: difference = 0.05, 95% CI [0.01, 0.09], adj.p = 0.0365) and middle doses (Veh-2.5 mg/kg: difference = 0.08, 95% CI [0.04, 0.11], adj.p < 0.001) increased response speed whereas the highest dose slowed responding compared to all other doses (Veh-7.5 mg/kg: difference = -0.05, 95% CI [-0.08, -0.01], adj.p = 0.031, 0.5 mg/kg-7.5 mg/kg: difference = -0.1, 95% CI [-0.13, -0.07], adj.p < 0.001, 2.5 mg/kg-7.5 mg/kg: difference = -0.13, 95% CI [-0.16, -0.09], adj.p < 0.001) (**Figure 4B**).

**Figure 4:**
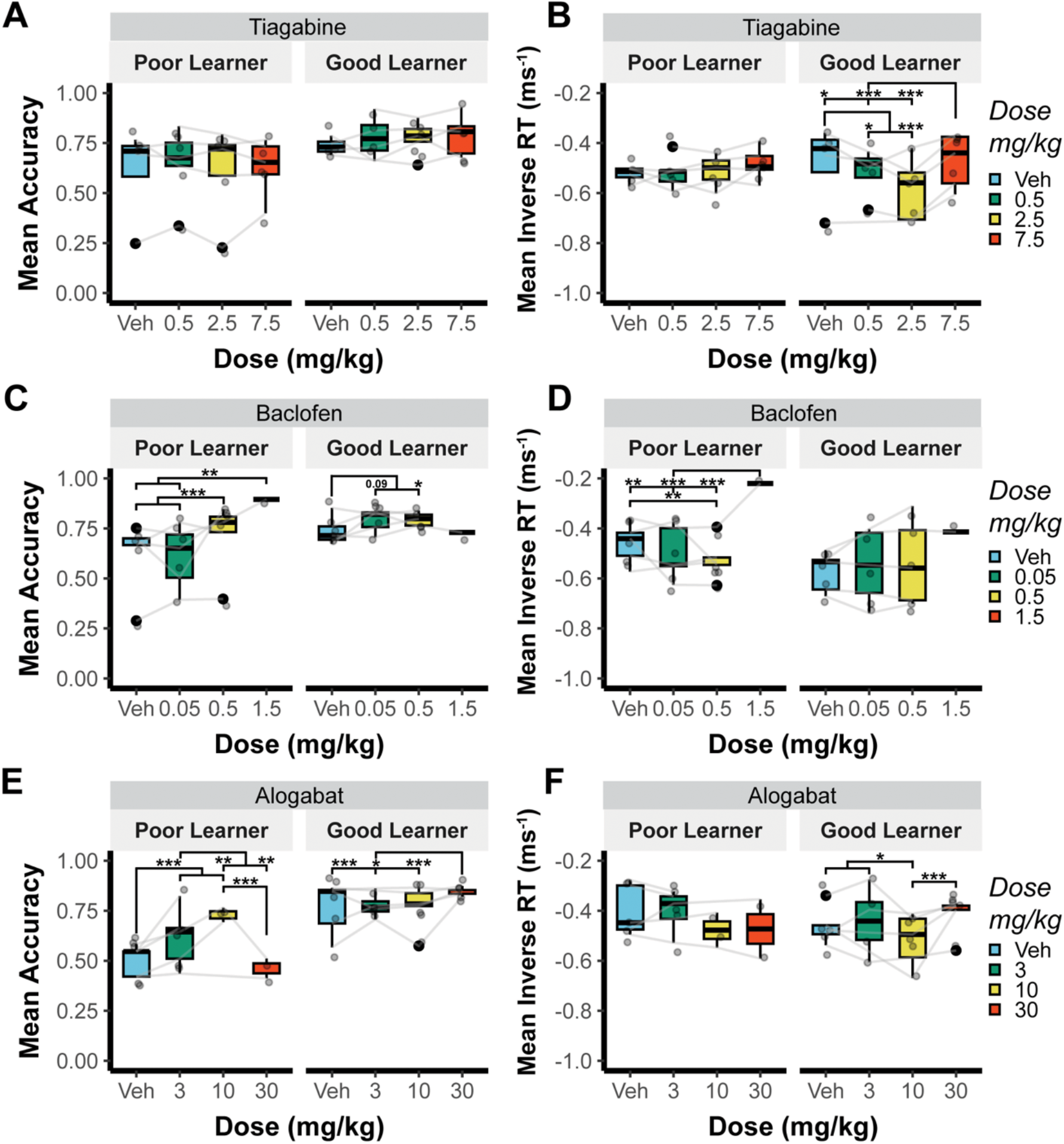
Baseline-dependent effects of GABAergic interventions on the VPL task. **A** Distribution of full task mean accuracy following the administration of the GAT-1 inhibitor tiagabine separated by learning group. **B** Distribution of full task mean inverseRT following the administration of the GAT-1 inhibitor tiagabine separated by learning group. **C** Distribution of full task mean accuracy following the administration of the GABA B agonist R-baclofen separated by learning group. **D** Distribution of full task mean inverseRT following the administration of GABA B agonist R-baclofen separated by learning group. **E** Distribution of full task mean accuracy following the administration of the GABRA5 positive allosteric modulator alogabat separated by learning group. **F** Distribution of full task mean inverseRT following the administration of the GABRA5 positive allosteric modulator alogabat separated by learning group. Generalized linear mixed-effects model with binomial link **(A,C,E).** Linear mixed-effects model **(B,D,F)**. Boxplots denote median, interquartile range with whiskers extending to 1.5x IQR. Outliers denoted by solid points. Points are individual animals, connected across doses with grey trajectories. Pairwise comparisons made via estimated marginal means to Vehicle condition in each case, within “Learning type”. Effect sizes: “p value” = p<0.1, “*” = p<0.05, “**” = p<0.01, “***” = p<0.001**(B,D,F)** and as log-odds ratios **(A,C,E)**.

Administration of R-baclofen affected accuracy in both learning groups (**Figure 4C**) (compared to Veh: 0.05 mg/kg, odds ratio = 0.98, 95% CI [0.78, 1.23], p = 0.8621, 0.5 mg/kg, odds ratio = 1.64, 95% CI [1.3, 2.06], p = 2.89e^-05^, 1.5 mg/kg, odds ratio = 2.93, 95% CI [1.49, 5.76], p = 0.0018. Here, middle-dose R-baclofen improved “Good” learner accuracy (Good: Veh-0.5 mg/kg: odds ratio = 0.72, 95% CI [0.56, 0.92], adj.p = 0.048), and had a positive effect on accuracy in “Poor” learners (Poor: Veh -0.5 mg/kg: odds ratio = 0.61, 95% CI [0.49, 0.77], adj.p < 0.001, 0.05 mg/kg – 0.5 mg/kg: odds ratio = 0.6, 95% CI [0.48, 0.76], adj.p < 0.001)). High-dose baclofen improved the accuracy of “Poor” leaners (Poor: Veh-1.5 mg/kg: odds ratio = 0.34, 95% CI [0.17, 0.67], adj.p = 0.0099, 0.05-1.5 mg/kg: odds ratio = 0.33, 95% CI [0.17, 0.66], adj.p = 0.008) but this comparison was relatively low powered due to subject exclusion (**Figure S6**). Low dose R-baclofen had no impact on “Poor” learner accuracy (Poor: Veh-0.05 mg/kg: odds ratio = 1.02, 95% CI [0.82, 1.28], adj.p = 0.9981), but there was a positive trend associated with “Good” learner performance (Good: Veh-0.05 mg/kg: Odds Ratio = 0.75, 95% CI [0.58, 0.95], adj.p = 0.0868). Whilst there was a main effect of R-baclofen dose on inverseRT (F(_3,139_) = 6.05, Pr(>F) = 6.68e^-04^), this was limited to “Poor” learners only. Here, dose wise comparisons revealed faster responses at the intermediate dose (Poor: Veh-0.5 mg/kg: difference = 0.06, 95% CI [0.03, 0.1], adj.p = 0.0026), and slowing at high doses (Poor: Veh-1.5 mg/kg: difference = -0.14, 95% CI [-0.21, -0.07], adj.p = 0.0014, 0.05-1.5 mg/kg: difference = -0.18, 95% CI [-0.25, -0.11], adj.p < 0.001, 0.5-1.5 mg/kg: difference = -0.21, 95% CI [-0.28, -0.13], adj.p < 0.001) compared to vehicle and low dose conditions (**Figure 4D**).

Like baclofen, alogabat administration affected accuracy in both learning groups (Veh-3 mg/kg, odds ratio = 1.64, 95% CI [1.32, 2.06], p = 1.30e^-05^, Veh-10 mg/kg, odds ratio = 3.11, 95% CI [2.24, 4.31], p = 9.87e^-12^, Veh-30 mg/kg, odds ratio = 0.89, 95% CI [0.65, 1.23], p = 0.4792). A within group analysis of accuracy in “Poor” learners revealed stepwise improvements in accuracy from vehicle at 3 mg/kg (Poor:Veh-3 mg/kg: odds ratio = 0.61, 95% CI [0.49, 0.76], adj.p < 0.001) and 10 mg/kg doses (Poor:Veh-10 mg/kg: odds ratio = 0.32, 95% CI [0.23, 0.45], adj.p < 0.001) whereas high dose exhibited no difference compared with the vehicle control group (Poor:Veh-30 mg/kg: odds ratio = 1.12, 95% CI [0.81, 1.55], adj.p = 0.8941. In contrast, a dose of 30 mg/kg alogabat improved “Good” learner accuracy (odds ratio = 0.54, 95% CI [0.41, 0.71], adj.p < 0.001) with lower doses showing comparable accuracy with the vehicle group (**Figure 4E**) (Veh-3 mg/kg: odds ratio = 0.86, 95% CI [0.65, 1.13], adj.p = 0.7006, Veh-10 mg/kg: odds ratio = 0.96, 95% CI [0.74, 1.23], adj.p = 0.9862). Like tiagabine, alogabat produced significant effects on response latencies in “Good” learners only. Thus, a dose of 10 mg/kg resulted in faster responses compared with vehicle (Veh-10 mg/kg difference = 0.06, 95% CI [0.02, 0.1], adj.p = 0.0275) and 3 mg/kg (3 -10 mg/kg: difference = 0.07, 95% CI [0.02, 0.11], adj.p = 0.0193), whilst 30 mg/kg slowed responding compared to the 10 mg/kg dose level (10-30 mg/kg: difference = -0.1, 95% CI [-0.14, -0.06], adj.p < 0.001) (**Figure 4F**).

### 3.4 Relating perceptual performance to GABRA5 expression

To identify putative substrates underlying individual variability in the effects of alogabat we next investigated GABRA5 expression in primary visual cortex (V1) and subregions of the mPFC (Cg1, PrL, IL – see **Figure 5A**) using Western immunoblot and RNA scope fluorescence in-situ-hybridisation (RNA Scope) (**Figure 5B-D**).

**Figure 5:**
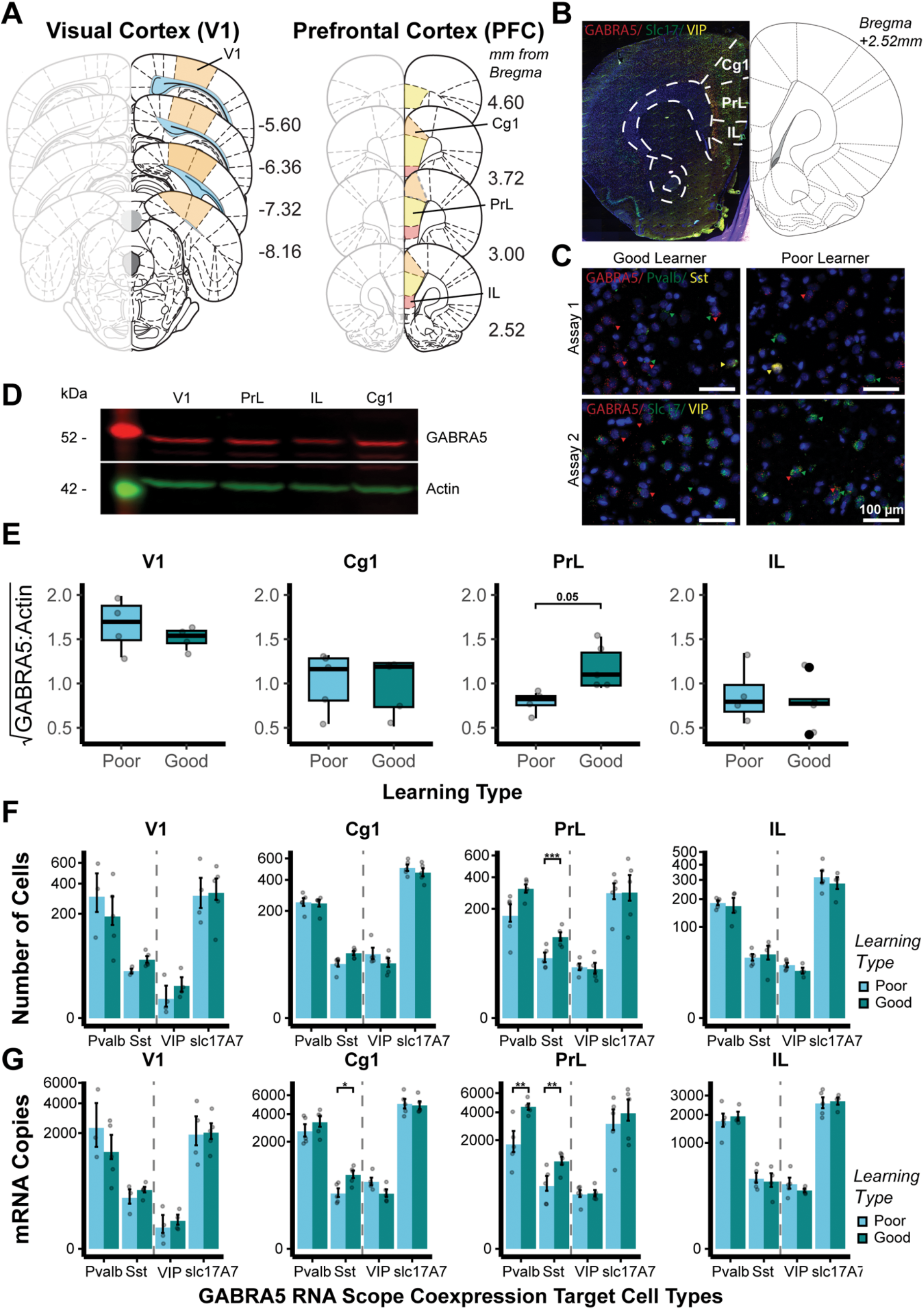
Relationship of GABRA5 expression with perceptual acuity and learning rate. **A** Schematic illustrations showing target regions of interest, left = primary visual cortex (V1) -5.6mm – -8.16 mm from bregma, right = prefrontal cortex (PFC) 4.6 mm – 2.52 mm from bregma. Sub-regions: anterior cingulate cortex (Cg1), prelimbic cortex (PrL), infralimbic cortex (IL). **B** Representative section of PFC showing fluorescent signal aligned with ROIs. **C** RNAscope assays from representative good versus poor learners on the VPL task. Scale bar 100 µm. Assay 1 GABRA5 (red); Pvalb (green); Sst (yellow): Assay 2 GABRA5 (red); slc17A7 (green); VIP (yellow). Arrows highlight cells expressing target probes in each case. **D** Representative Western-immunoblot gel for √GABRA5 protein expression in good versus poor learning groups with actin as the housekeeper control for each ROI. **E** Statistical comparison of Western-immunoblot derived GABRA5 protein expression in cortical ROIs between stratified “learning types”, presented as sqrt GABRA5:Actin. **F** Cell-wise quantification of GABRA5 co-expression from RNAscope assays within inhibitory interneuron subtypes: parvalbumin (Pvalb), somatostatin (Sst), vasoactive intestinal peptide (VIP) and excitatory pyramidal cells (slc17A7) by “Learning type”, per target cortical region. **G** Puncta-wise quantification of GABRA5 co-expression from RNAscope assays within inhibitory interneuron subtypes: parvalbumin (Pvalb), somatostatin (Sst), vasoactive intestinal peptide (VIP) and excitatory pyramidal cells (slc17A7) by “learning type” in target cortical regions. A modulus transformation (p=0.3) was applied to scale the RNAscope y-axes to enable better interpretability between high and low expression values on the same plot. mRNA coexpression values are displayed for each “learning type” as mean ± SEM per subject. RNAscope was performed using 2 assays to assess GABRA5 within (1: Pvalb, Sst, 2: VIP, slc17A7) highlighted with dashed line. Pairwise comparisons made via linear mixed-effects model to “poor learner” condition each case via estimated-marginal-means, within region and cell type: “p value” = p<0.1, “*” = p<0.05, “**” = p<0.01, “***” = p<0.001.

Western blot protein analysis highlighted a significant main effect of region in GABRA5 protein expression F(_3,20.97_) = 15.51, Pr(>F) < 0.001. This effect was driven by V1 expression compared to the mPFC regions (V1-Cg1: difference = 0.61, 95% CI [0.36, 0.85], p = 0.0002, V1-PrL: difference = 0.62, 95% CI [0.36, 0.87], p = 0.0003, V1-IL: 0.77, 95% CI [0.52, 1.02], p < 0.0001). To test our primary hypothesis that protein levels would vary between learning types, *post hoc* comparisons were performed within each region. These revealed marginally greater GABRA5 expression in the PrL of good learners only (Poor -Good: difference = -0.38, 95% CI [-0.75, -0.02], adj.p = 0.0512). All other regions were not statistically different (Poor-Good V1: difference = 0.19, 95% CI [-0.19, 0.57], adj.p = 0.3417, Cg1: difference = 0.04, 95% CI [-0.3, 0.39], adj.p = 0.81, IL: difference = 0.09, 95% CI [-0.27, 0.46], adj.p = 0.6282) (**Figure 5E**).

We then assessed GABRA5 mRNA in selective subpopulations of inhibitory interneurons, specifically: parvalbumin (Pvalb)-, somatostatin (Sst)-, vasoactive intestinal peptide (VIP)-and excitatory pyramidal cells (slc17A7) in each target cortical region of interest. After validating uniform cell counts across learning types within regions (**Figure S5)**, GABRA5-inhibitory interneuron co-expression differences were assessed. Statistical comparison between learning types revealed a significantly increased number of GABRA5 and Sst co-expressing cells in the PrL of good learners (Poor -Good: difference = -49.93, 95% CI [-74.43, -25.43], adj.p < 0.001) (**Figure 5F**), an increased number of GABRA5 and Sst overlapping mRNA copies in the Cg1 and PrL of good learners (Cg1:Poor -Good: difference = -437, 95% CI [-766.28, -107.72], adj.p = 0.0187, PrL: Poor -Good: difference = -563, 95% CI [-892.28, -233.72], adj.p = 0.0038), and a greater number of GABRA5 and Pvalb overlapping mRNA copies in the PrL of good learners (Poor -Good: difference = -2945, 95% CI [-4562, -1328], adj.p = 0.0031) (Figure 5G). There was no difference in within-region total-DAPI derived learning-type cell counts (LT:Region – assay 1: F(_3,16.8_) =0.202, Pr(>F) = 0.8935, assay 2: F_(3,22.4)_ = 0.102, Pr(>F) = 0.958). All other comparisons were non-significant. A summary of the molecular quantification data is given in supplementary **Table 2**.

## 4.0 Discussion

In this study we investigated the role of inhibitory neurotransmission in visual perceptual learning. Using a novel orientation discrimination task, we assessed the association between GABRA5 expression and task acquisition and validated our findings with selective GABAergic pharmacology in male and female animals. We report that superior performance on our task is accompanied by increased GABRA5 expression in Sst and Pvalb interneuronal populations in the prelimbic subregion of the medial prefrontal cortex. Since VPL performance in poor learners was improved following positive allosteric GABRA5 modulation during the early task sessions, our findings indicate that features of the present VPL task such as attentional selection and suppression of distractor interference may depend in part on GABRA5 functional activity, potentially within the prelimbic cortex (PrL).

Orientation discrimination is a commonly employed behavioural paradigm to investigate perceptual acuity in rodents (Speed et al. 2020; Poort et al. 2022) and humans (Edden et al. 2009; Jia et al. 2018). However, cross-species translation is often challenged by differences in how VPL tasks are implemented. We therefore developed an orientation discrimination task that assesses VPL in an analogous way to visual perceptual tasks in humans. By presenting target and distractor gratings close or far from one another, we reduced procedural location-based learning, widened the window for pharmacological intervention, and increased attentional demand (Yoo et al. 2022) in a way that resembles the high attentional load of equivalent human VPL tasks (Edden et al. 2009). Crucially, our approach preserved fine discrimination without restraint, facilitating naturalistic behaviour and negating the limitations of head fixed approaches commonly used in go/no-go-based perceptual tasks. To investigate intrinsic differences in perceptual learning in our task, we stratified behaviour at the end of pretraining. Here, stimulus reward associations (PT2+3), distractor interference tolerance (PT4), and spatial flexibility (PT2-PT4) were assessed. Our approach separated slow *versus* fast learners and asked whether GABRA5 expression patterns in posterior and anterior regions of the visual processing pathway are related to behavioural features of the VPL task.

VPL involves the integration of bottom-up sensory processing and coordinated responses from top-down attentional control mechanisms (Schäfer et al. 2007; Zhang et al. 2014; Bastos et al. 2015; Kok et al. 2016; Huh et al. 2018; Barbera et al. 2022). In the present study, rats showing superior visual perceptual learning exhibited increased GABRA5 gene expression in subpopulations of inhibitory interneurons in the PrL with convergent changes in GABRA5 protein expression. These concordant findings suggest that inhibitory processing in the PrL may be a putative substrate of visual perceptual learning. Supporting this notion, animals showing faster rates of learning were also more tolerant of a wider separation between the target and distractor stimuli, consistent with a recognized role of the PrL in maintaining attentional focus in the face of distraction (Sharpe and Killcross 2014; Auger et al. 2017). Indeed, higher cortical regions including the PFC and parietal association cortex (PC) are implicated in the control of interference suppression (Wöstmann et al. 2022) in response to multiple simultaneously presented visual stimuli on such tasks as multiple-object-tracking (MOT) (Meyerhoff et al. 2017) and visual discrimination (Frangou et al. 2019).

Within each test session we found block-wise differences in performance between fast and slower learners, which diminished toward the narrowest orientation angle (10° from target). This observation suggests that the TDD and perceptual challenges in our paradigm are distinct. In addition, while faster contingency learning during pretraining appears beneficial to overall task performance, when fine levels of discrimination were required, strategies to suppress interference may have been ineffective. Under these conditions, processing may depend instead on bottom-up sensory integration and value-based associative processing by regions such as the orbitofrontal cortex (Gourley et al. 2016; Marquardt et al. 2017). Our findings suggest that increased GABRA5 expression, putatively within the PrL, may enable faster acquisition of top-down control mechanisms to enable effective suppression of distractor interference. However, during late VPL, a limit is reached for fine discriminations that may be independent of GABRA5 functional expression.

To validate the role of GABRA5 in our VPL task we administered the GABRA5 PAM, alogabat and compared its effects to compounds that potentiate metabolic (tiagabine) and post-synaptic tonic inhibitory activity (R-baclofen). Notably, these compounds are reported to exhibit pro-cognitive activity in other contexts (Sałat et al. 2015; Stoppel et al. 2018). Our findings revealed a lack of effect of tiagabine and a dose-dependent improvement in discriminative accuracy following the administration of baclofen in both poor and good learning male rats. Thus, the post-synaptic potentiation of inhibition via the GABA B receptor improves perceptual acuity regardless of task acquisition rate. By contrast, in poor learning subjects only, alogabat both enhanced and impaired VPL performance in an apparently U-shaped function. Thus, potentiating GABRA5 within a relatively narrow dose range accelerates learning in a baseline dependent manner. The partial overlap in the effects of baclofen and alogabat on discriminative accuracy suggests both compounds may act directly or indirectly on a shared inhibitory mechanism. Based on prior evidence this shared mechanism may involve effects on tonic inhibition.

Tonic GABA currents are typically viewed as constraining learning and memory while sharpening sensory processing (reviewed by (Koh et al. 2023)). Although GABA B receptors contribute to ambient inhibitory regulation (Scanziani 2000; Bettler et al. 2004), several extrasynaptic GABA A subtypes including GABRA5 also mediate shunting inhibition (Caraiscos et al. 2004; Farrant and Nusser 2005; Perez-Sanchez et al. 2017). This GABA A-based mechanism has been proposed to narrow the temporal integration window of incoming signals, thereby increasing the signal-to-noise ratio and improving the fidelity of sensory representations (Pavlov et al. 2009; Brickley and Mody 2012). In this way, pharmacological potentiation of GABRA5 would be expected to bias processing toward stability and noise-suppression, potentially enhancing distractor resistance. Indeed, consistent with the clinical indications of alogabat (Cecere et al. 2025) and other pro-cognitive GABRA5 PAMs (Koh et al. 2013; Bernardo et al. 2025) modulating GABRA5 may have time-dependent benefits for early learning-related plasticity. However, once stimulus representations are well trained, ceiling effects of tonic GABA inhibition may act to constrain further improvements in discriminative accuracy.

Our gene expression studies indicated that higher GABRA5 expression in Pvalb and Sst neuronal populations of the mPFC is associated with early task learning and long-term baseline performance. Based on established functional interactions between GABRA5 and Sst interneurons involving gating of dendritic inputs to pyramidal cells (Pfeffer et al. 2013; Schulz et al. 2018; Cao et al. 2020; Fee et al. 2021; Huang 2025), we hypothesise that sensory signal-to-noise ratio and input integration mediated by these cells may be an important determinant of distractor suppression mediated by the PrL during VPL acquisition. A putative involvement of Pvalb interneurons also suggests GABRA5+ cells contribute to PrL recurrent inhibitory activity to fine tune the local E/I balance, thereby enhancing task acquisition via trial-by-trial prediction error updates (Znamenskiy et al. 2024; Jia et al. 2024). Despite previous evidence highlighting pyramidal cell enrichment of GABRA5 (Hu et al. 2019), our findings show a similar GABRA5 expression ratio between excitatory (slc17a7) and inhibitory (Pvalb and Sst) cell types. Further research is needed to resolve how innate and learning-related plasticity mechanisms affect the relative expression of GABRA5 in excitatory and inhibitory neuronal populations in the PFC. Nevertheless, our findings are consistent with the notion that clustering of GABRA5 in PrL Pvalb and Sst cells could bias inhibitory microcircuits towards greater tonic inhibitory tuning of sensory input fidelity. This proposed mechanism could account for faster learning in animals exhibiting a higher intrinsic expression of GABRA5.

There are however several limitations of our research. Firstly, the different rates of learning between male and female rats preclude meaningful sex-dependent comparisons to be made in relation to drug effects and GABRA5 dependent processes. Although the baseline VPL performance of male and female animals was ultimately similar, follow-up work would be required to establish and characterise sex-dependent differences in task acquisition and how these are shaped by cortical inhibitory mechanisms. Secondly, we acknowledge that the timing of the drug administration during learning makes the separation between task acquisition and performance difficult to disentangle. Nonetheless, the directional convergence of our gene expression studies with the behavioural effects of alogabat and baclofen provide an additional level of validation implicating GABRA5 expressing GABA A receptors in VPL. Thirdly, as static readouts in post-mortem samples, our gene expression and Western blot studies cannot directly be related the dynamic behavioural acquisition of the present VPL task. Thus, caution is needed in resolving whether the reported findings reflect true differences in innate receptor expression patterns or instead learning-and/or drug-related plasticity.

In conclusion, we report converging lines of evidence supporting an involvement of GABRA5 in visual perceptual learning. Based on the present findings and previously published reports, we argue that increased expression and functional activity of this GABA A subunit facilitates the acquisition of VPL by suppressing visual interference during task acquisition. Elucidating whether this process causally depends on tonic inhibitory signalling in Sst and Pvalb expressing interneurons of the prelimbic cortex may inform novel therapeutic approaches to restore deficits in visual perceptual learning in schizophrenia and other neuropsychiatric disorders.

## Supporting information

Full Supplementary Material until table S1

## Acknowledgements & Author Contributions

The experimental work was carried out under a Home Office Project Licence held by Prof. Dr. A. L. Milton. Conceptualization: MCDB. Experimental design: MCDB, JFdh, JWD. Data analysis: MCDB, OSRPS, LVW. Investigation: MCDB, EP, CVS, LMP. Visualization: MCDB, EP, CVS, LMP. Supervision: JFdH, JWD. Writing–original draft: MCDB JWD. Writing–review & editing: all authors

