## Supplementary material for "Prefrontal α5-GABA-A receptors: putative role in perceptual learning": Full Supplementary Material until table S1

### Supplementary Information

#### Figure S1: Visual Discrimination Training

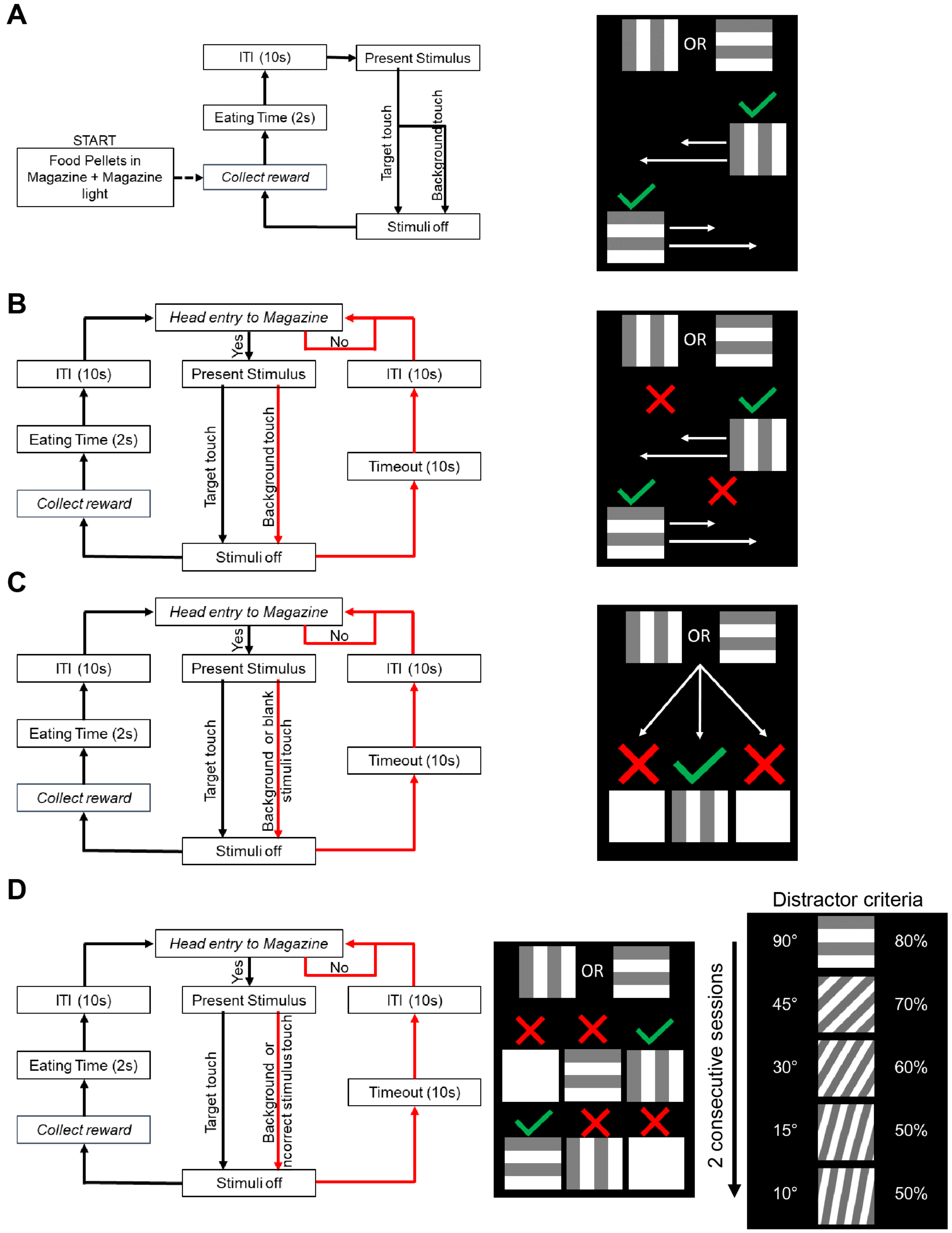


**Figure S1: Visual discrimination task training stages and criteria for progression.** (A) Pre-training 2 (PT2) single target (S+) presented in 3 windows, no background punishment. (B) Pre-training 3 (PT3) single target (S+) presented in 3 windows, background punishment, trial self-initiation. (C) Pre-training 4 (PT4) single target (S+) presented in 3 windows, 2 blank dummy stimuli, background punishment. (D) Visual discrimination (VD) stages 1-5 single target (S+) presented in 3 windows, 1 blank and 1 distractor stimuli of increasing difficulty, background punishment. Progression with 80% correct on 2 consecutive sessions up to VD1. VD1 progression in right panel. Animals that failed to reach criteria >5 sessions on VD5 were moved to final task.

### Supplementary Information

#### Figure S2: Sex Differences in Task Acquisition

**A**

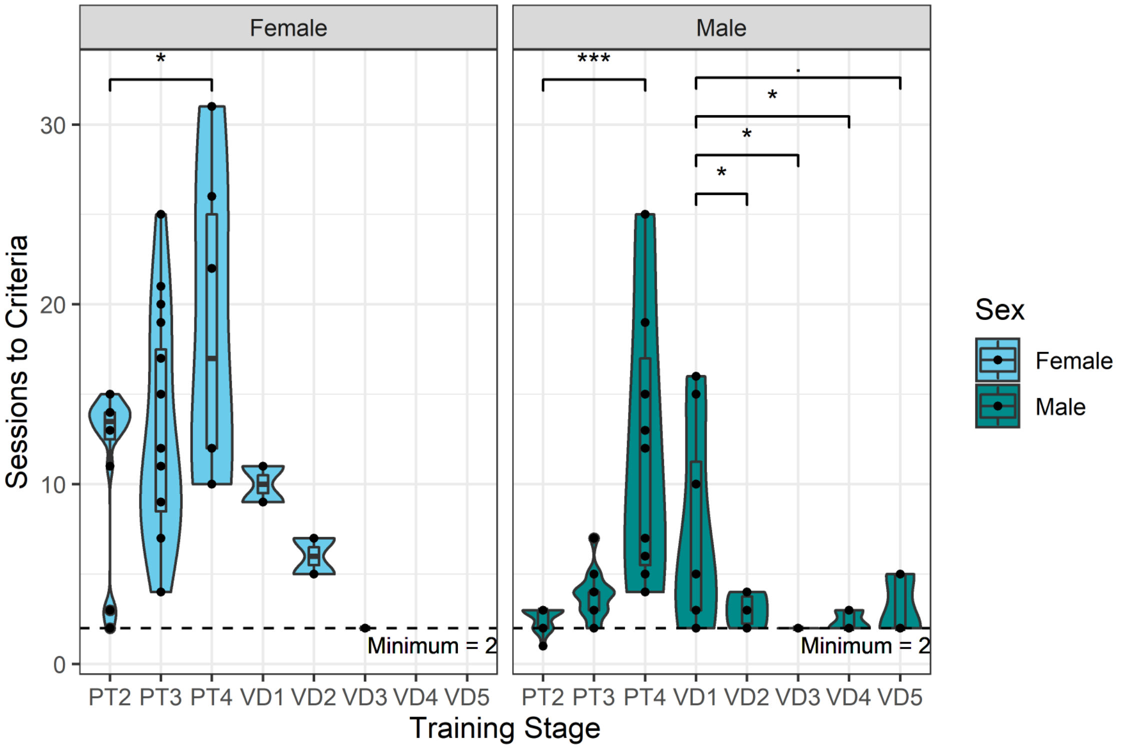


**B**

| Class | Males | Females | Total Animals |
|---|---|---|---|
| Poor Learner | 6 | 8 | 14 |
| Inter Learner | 5 | 8 | 13 |
| Good Learner | 5 | 0 | 5 |

**Figure S2: Visual discrimination task learning rate varies at different training stages:** (A) Violin and boxplots of distributions of sessions completed to criteria for each training stage split females (left) and males (right). PT2-PT4 = pretraining, VD1-VD5 = visual discrimination training. Dashed line is minimum sessions at criteria for progression for all sessions apart from PT2 which is 1. All comparisons made via mixed effects linear model evaluation: "." = P<0.1, "*" = p<0.05, "**" = p<0.01, "***" = p<0.001. pretraining comparisons made to PT2 reference and visual discrimination training comparisons made to VD1 reference. (B) Summary table of learning type classes based on stage reached at the end of pretraining.

### Supplementary Information

#### Figure S3: Statistical Model Technique

**Map Variables**

**Independent:**
- Angular separation from target
- Target distractor distance

**Dependent:**
- Choice correct trialwise
- Choice latency trialwise

**Manipulations**
- Drug/Dose

**Grouping Variables**
- Sex
- Learning Type

**Random Effects**
- Subject

**Check Distributions**

**Qualitative**
- Histogram

**Quantitative**
- Shapiro-Wilk test

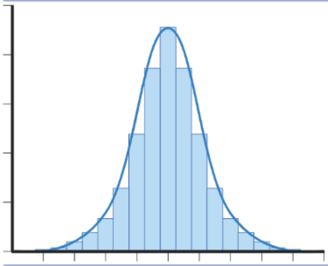

```
        Shapiro-Wilk normality test

data:  Summary_Latency_baseline_EXC$mean_i_latency_choice
W = 0.98886, p-value = 0.5035
```

**Specify Models**

**Normal -** Linear mixed model (lmer)
**Non-Normal -** Generalised linear mixed model (glmer)
- OR - transform **then** lmer

**Dependent Variables:**
- Choice correct (binary), non-normal - **glmer**
- Inverse transformed choice latency (-100/latency), normal - **lmer**

**Generate models of interest**, including all variations of fixed effects

```
baseline_acc_Sex_lme_full <- glmer(choice_correct ~ ang_sep*stim_dist_loc*Sex + (1|Rat_ID),
                        data = baseline_data,
                        family = binomial,
                        control = control_settings,
                        nAGQ = 20)

baseline_acc_Sex_glme_add_int <- glmer(choice_correct ~ ang_sep*Sex + stim_dist_loc*Sex + (1|Rat_ID),
                        data = baseline_data,
                        family = binomial,
                        control = control_settings,
                        nAGQ = 20)

baseline_acc_Sex_glme_nloc <- glmer(choice_correct ~ ang_sep*Sex + (1|Rat_ID),
                        data = baseline_data,
                        family = binomial,
                        control = control_settings,
                        nAGQ = 20)

baseline_acc_Sex_glme_ang_sep <- glmer(choice_correct ~ ang_sep + (1|Rat_ID),
                        data = baseline_data,
                        family = binomial,
                        control = control_settings,
                        nAGQ = 20)
```

Optimise

**Evaluate Models**

**Apply Model Goodness of Fit Tool -** Performance Package in R
Pick **best fitting** model
**Plot Diagnostic plots + simulated residuals** to check generalisation

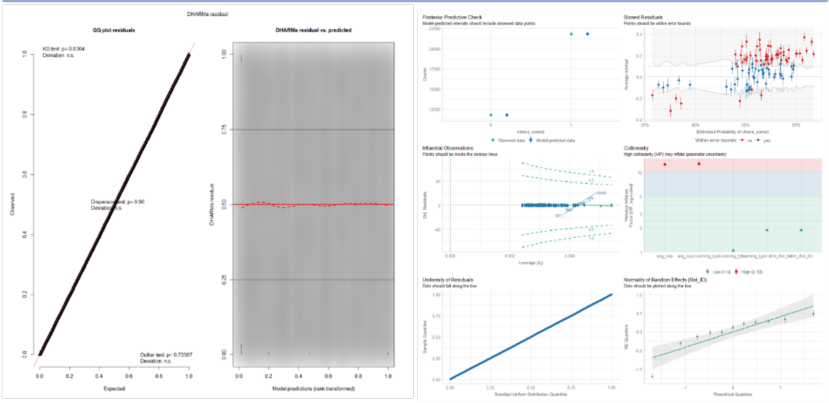


**Apply Best Model**

**Interpret diagnostics** - does it not violate assumptions + generalise well?
**A) Yes -** Continue and apply model to dataset
**B) No -** Alter model, and iterate through evaluation again

### Supplementary Information

**Figure S3: Mixed model specification and evaluation framework** Models derived from classical assumption-based processes. Variables are first mapped for their data types and normality distributions, before specifying mixed models with the equation **model = lmer/glmer(dep_var ~ ind_var1 */+ ind_var2 + (1|Subject)). E.g Glmer** with binomial link function applied to binary response data for choice_correct (trial-by-trial, 0 or 1). Model fit compared between fixed effects structures using performance package (compare_performance()) and anova (likelihood ratio testing (LRT), anova()) between models of increasing complexity. Best LRT value model assessed via full diagnostic plots (performance package, check_model()) and simulated residuals (DHARMa package, residuals_sim()). Poor generalisation results in iteration through simpler models. Process repeated until reasonable generalisation and LRT reached with sufficient fixed effect explanatory structure. Figure created in https://app.biorender.com/

Models used in analysis

**Baseline behaviour**

```
Sex_acc_model <- glmer(choice_correct ~ Sex* ang_sep + Sex*TDD + (1|Rat_ID),
                data = baseline_data,
                family = binomial,
                control = control_settings,
                nAGQ = 20)
LT_acc_model <- glmer(choice_correct ~ LT* ang_sep + LT*TDD + (1|Rat_ID),
                data = baseline_LT_data
                family = binomial,
                control = control_settings,
                nAGQ = 20)

Sex_inverseRT_lme <- lmer(mean_i_latency_choice ~
                 ang_sep*Sex +TDD*Sex + (1|Rat_ID),
                data = Summary_Latency_baseline_Sex)

LT_inverseRT_lme <- lmer(mean_i_latency_choice ~
                 ang_sep*Learning_type +TDD*Learning_type + (1|Rat_ID),
                data = Summary_Latency_baseline_LT)
```

**Behavioural pharmacology**

```
Drug_acc_model <- glmer(choice_correct ~ Learning_type*`Dose_mg/kg`+ang_sep + (1|Rat_ID),
                data = drug_data,
                family = binomial,
                control = control_settings,
                nAGQ = 20)
Drug_lat_model <- lmer(mean_i_latency_choice ~ Learning_type*`Dose_mg_kg`+ang_sep + (1|Rat_ID),
                data = inverseRT_drug_data)
```

**Molecular investigations**

```
RNAScope_lme <- lmer(`GABRA5+_cells` ~ Learning_type*Region + (1|id),
                data = RNAScope_df)
WB_lme <- lmer (GABRA5_Actin ~ Learning_type*Region + (1|id),
                data = WB_df)
control_settings = “bobyqa”
```

### Supplementary Information

#### Figure S4: RNA Scope Controls

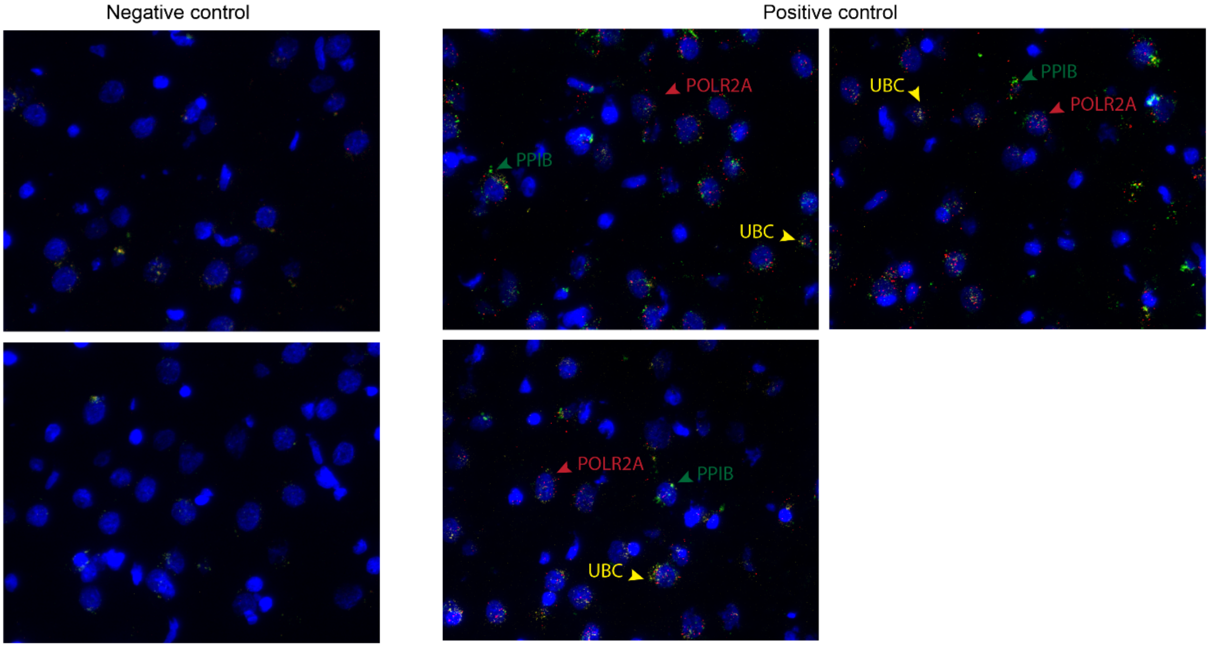


**Figure S4:** Negative and positive control probes were processed in parallel with the target probes to ensure tissue RNA integrity and optimal assay performance. The positive control probe is a mixture of three probes targeting POLR2A in channel 1 (red), PPIB in channel 2 (green), and UBC in channel 3 (yellow). The negative control probe recognizes DapB a bacterial transcript.

#### Figure S5: RNA Scope DAPI Cell Count

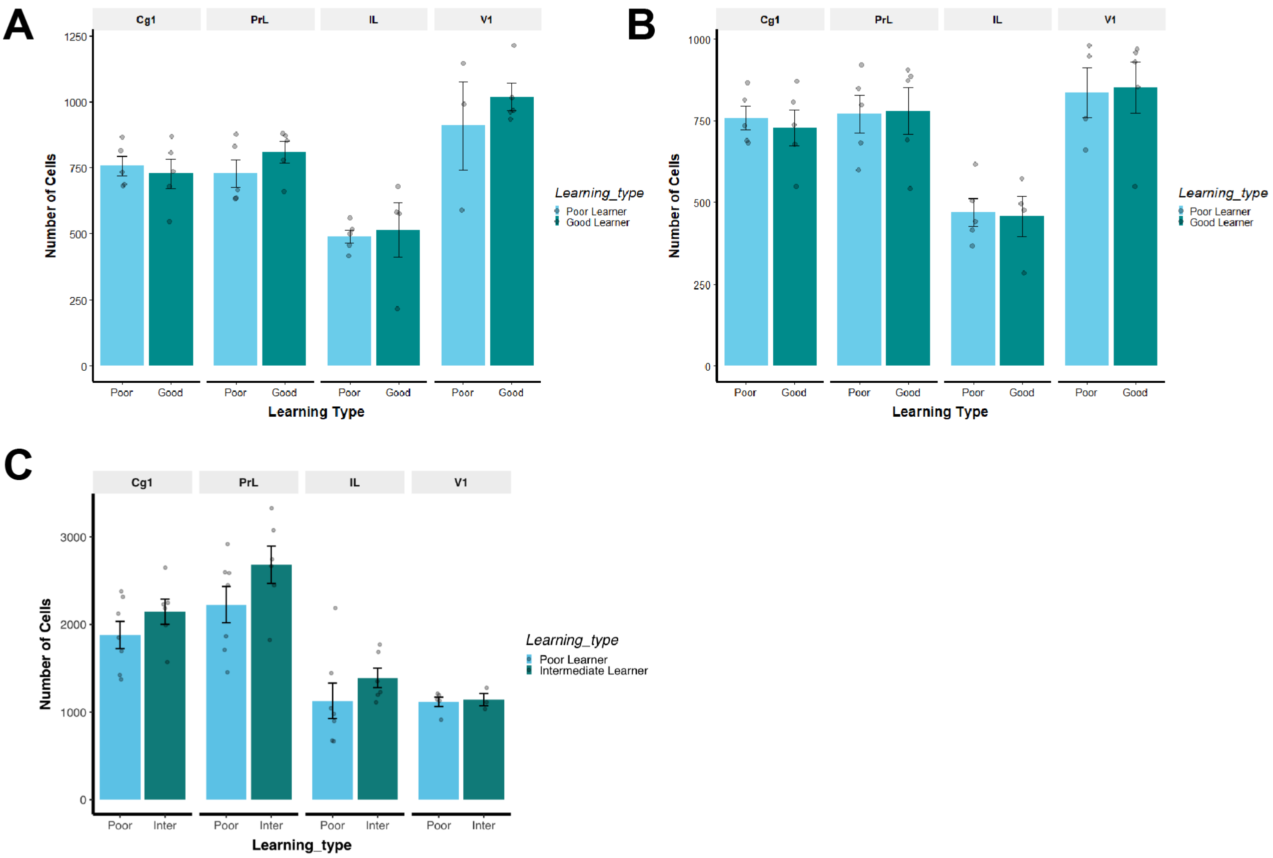


**Figure S5: RNA Scope DAPI Cell Count** A) Males Assay 1, B) Assay 2 And C) Female Assay 1 mean ± SEM DAPI cell counts between groups. Poor-Good (M), Poor-Intermediate (F). NS found within learning type groups of each region. Comparisons performed by Linear mixed effects models between “Learning_Type” to “Poor” reference condition within region.

#### Supplementary Information

### Figure S6: Trial Completion by Compound

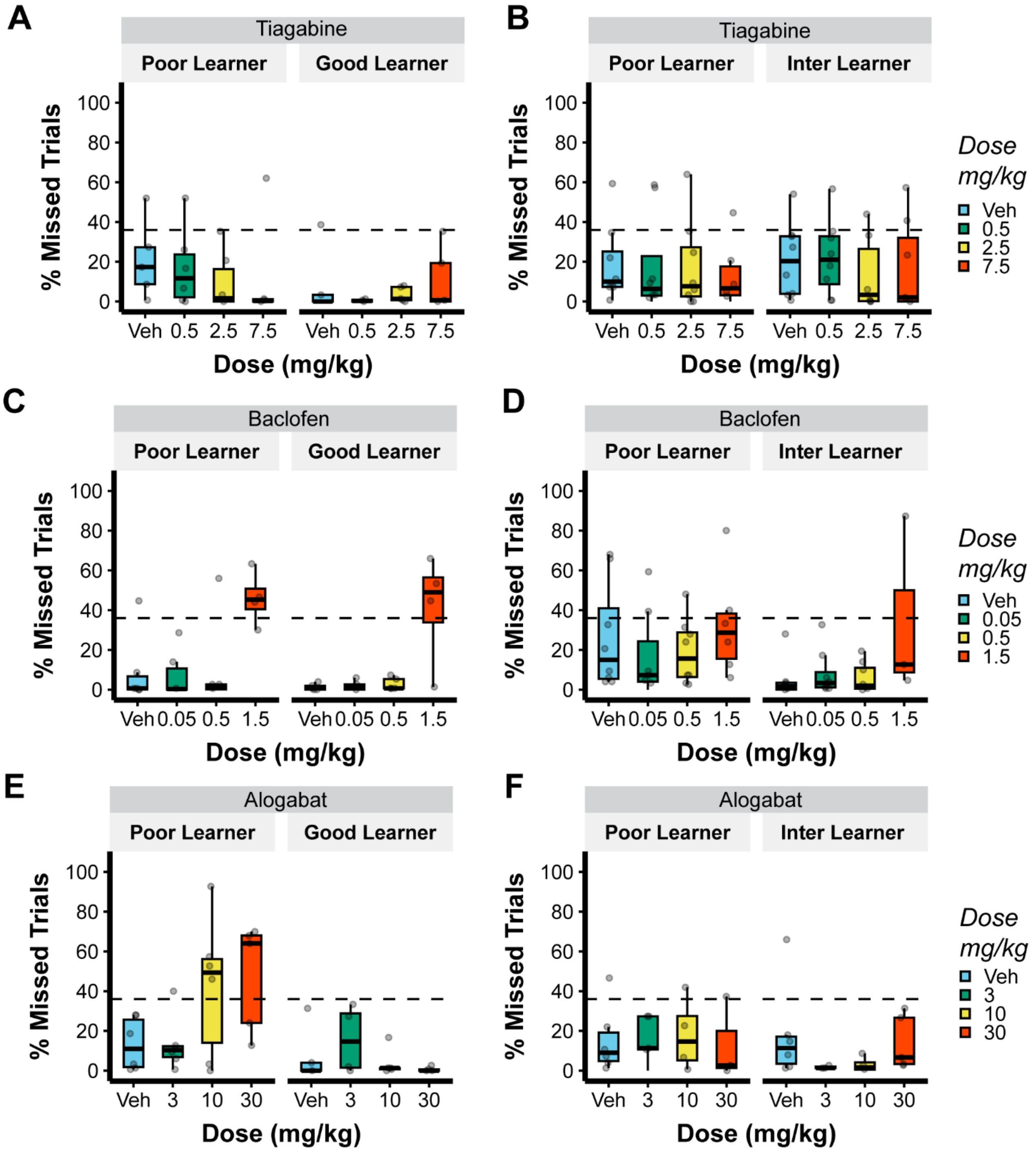


**Figure S6: Trial Completion by Compound** A) Males Tiagabine B) Females Tiagabine C) Males R-Baclofen D) Females R-Baclofen E) Males Alogabat F) Females Alogabat. Dashed line is inclusion cut-off for statistical analysis - >96/150 trials completed. Boxplots denote median, interquartile range with whiskers extending to 1.5x IQR. Outliers denoted by solid points. No statistical comparisons were performed between doses.

### Supplementary Information

#### Figure S7: Female Stratification

**A**

| Learning Type | Criteria | Total Animals |
|---|---|---|
| Poor Learner | Failed complete Pre-training | 8 |
| Intermediate Learner | Reached Discrimination training | 8 |

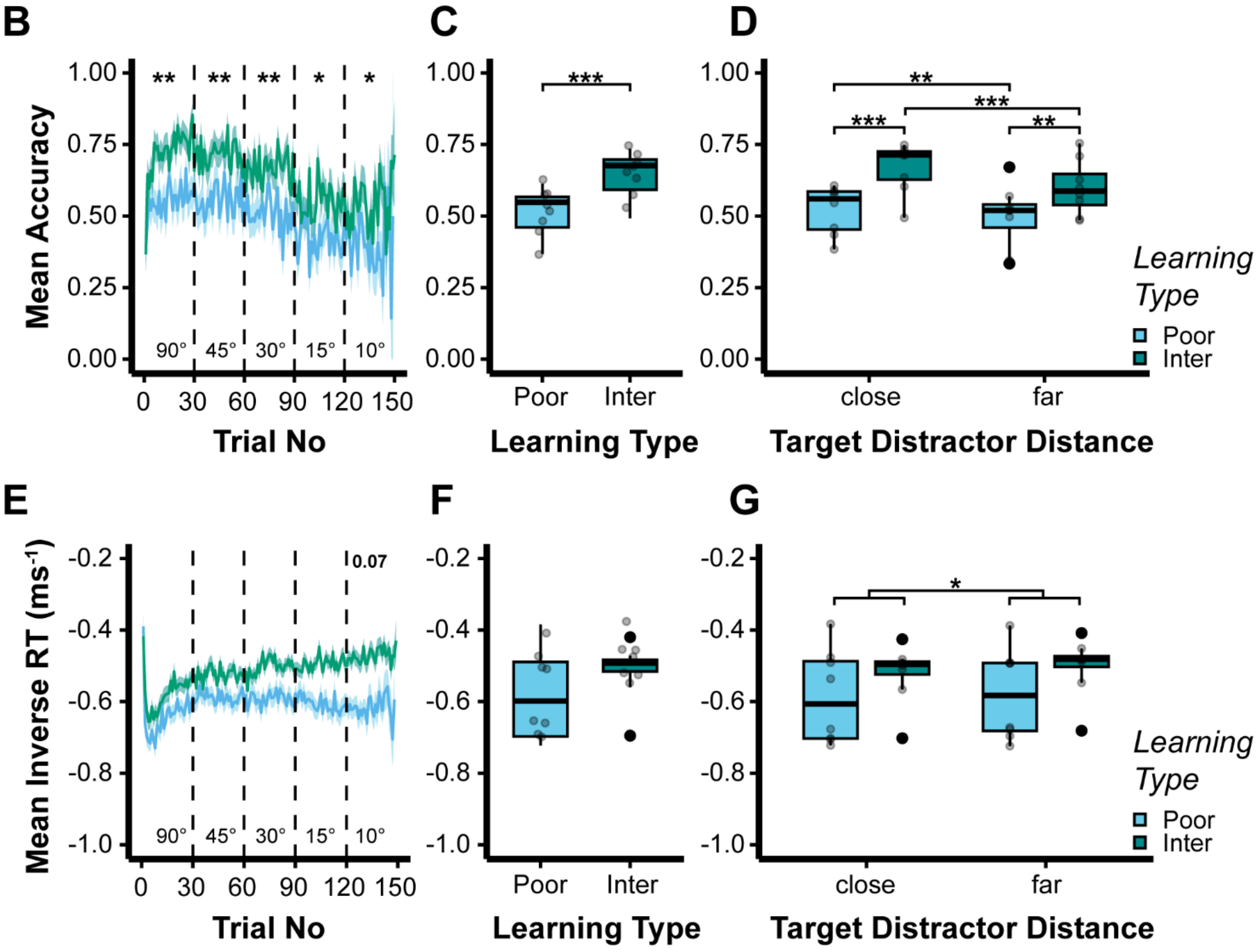


**Figure S7: Female Task acquisition rate predicts perceptual performance**
**A** Summary of stratification based on training stage reached at the end of 53 sessions of pretraining **B** Trial-wise mean accuracy by learning type presented as mean ± SEM. Dashed lines mark distractor frequency blocks. Annotations denote between-learning type comparisons, within distractor frequency block **C** Distribution of all-time full task baseline accuracy by learning type, points denote individual animals **D** Distribution of all-time full task mean session accuracy by target distractor distance. Points denote individual animals per TDD. Comparisons made within TDD-between learning type and within learning type-between TDD.
**E** Trial-wise mean inverseRT by learning type presented as mean ± SEM. Dashed lines mark distractor frequency blocks. Annotations denote between-learning type comparisons, within distractor frequency block **F** Distribution of all-time full task mean session inverseRT by learning type, points denote individual animals
**G** Distribution of all-time full task mean session inverseRT by target distractor distance. Points denote individual animals per TDD. Comparisons made within TDD-between learning type and within learning type-between TDD.
All-time baseline sessions displayed for each learning type with statistical comparisons made by generalized linear mixed-effects model with binomial link for accuracy (**B-D**), Linear mixed-effects model for inverseRT (**E-G**). Boxplots denote median, interquartile range with whiskers extending to 1.5x IQR. Outliers denoted by solid points. Pairwise comparisons made via estimated-marginal-means to “Poor learner” condition in each case, within block **(B,E)** and “close” **(D,G)**. Effect sizes: “*” = p<0.05, “**” = p<0.01**(E-G)** as log-odds ratios **(B-D)**.

### Supplementary Information

#### Figure S8: Female Pharmacology

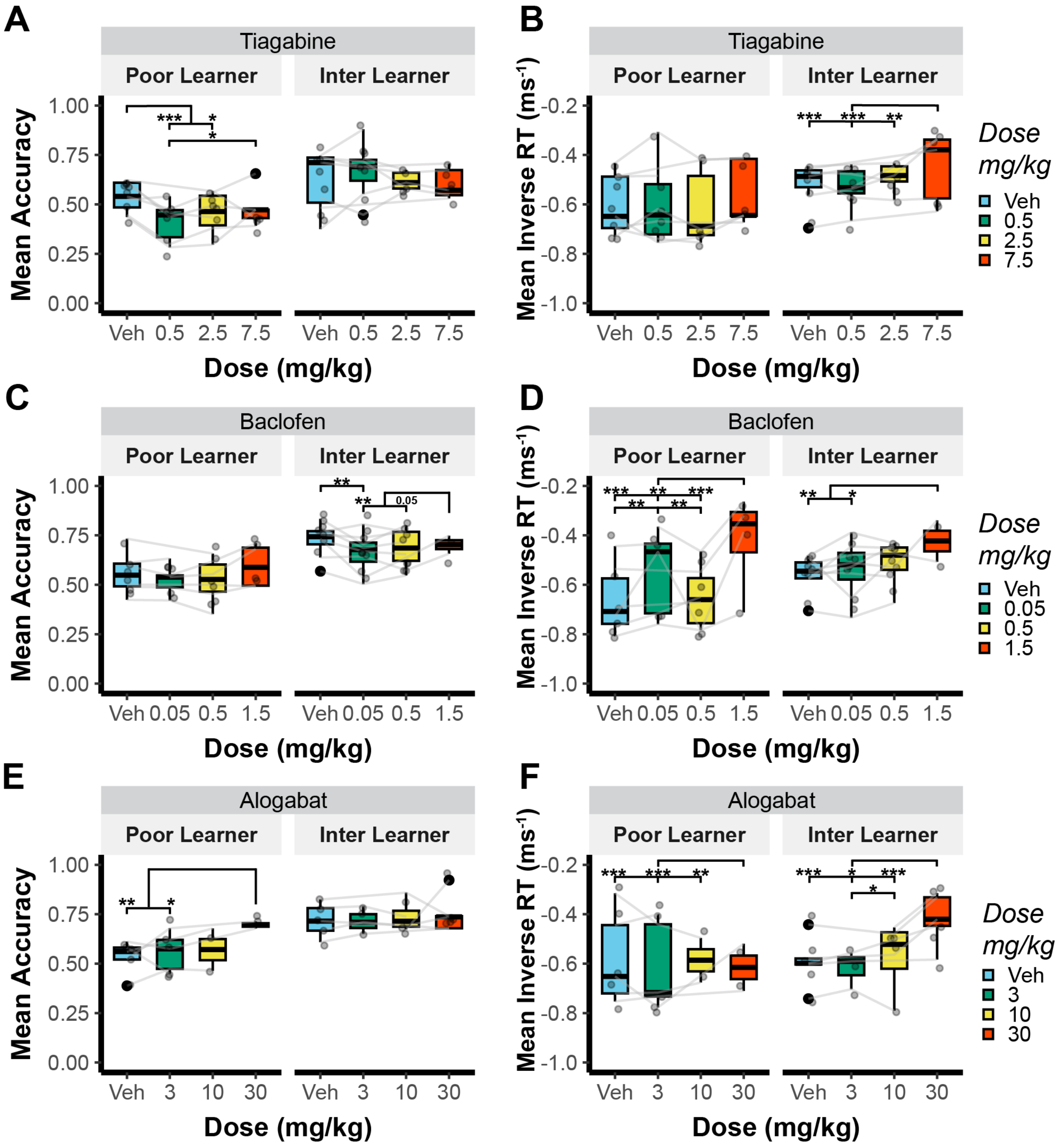


**Figure S8: Task acquisition governs response to GABAergic pharmacology in Females A** Distribution of full task mean accuracy with administration of 3 doses of the GAT-1 inhibitor Tiagabine within learning type **B** Distribution of full task mean inverseRT with administration of 3 doses of the GAT-1 inhibitor Tiagabine within learning type. **C** Distribution of full task mean accuracy with administration of 3 doses of the GABA B agonist R-Baclofen within learning type **D** Distribution of full task mean inverseRT with administration of 3 doses of the GABA B agonist R-Baclofen within learning type
**E** Distribution of full task mean accuracy with administration of 3 doses of the GABRA5 positive allosteric modulator Alogabat within learning type **F** Distribution of full task mean inverseRT with administration of 3 doses of the GABRA5 positive allosteric modulator Alogabat within learning type.
Generalized linear mixed-effects model with binomial link **(A,C,E)**, Linear mixed-effects model **(B,D,F)**. Boxplots denote median, interquartile range with whiskers extending to 1.5x IQR. Outliers denoted by solid points. Points are individual animals connected across doses. Pairwise comparisons made via estimated marginal means to Vehicle condition in each case, within “Learning type”. Effect sizes: “p value” = p<0.1, “*” = p<0.05, “**” = p<0.01, “***” = p<0.001**(B,D,F)** and as log-odds ratios **(A,C,E)**. Subjects with <2 doses/compound were excluded from analysis. n: Poor Learners = 8, Intermediate Learners = 8 but varies by drug/dose, see supplementary figure 6 for dose-wise exclusion thresholds based on <96/150 trials complete.

### Supplementary Information

#### Figure S9: Female GABRA5 Molecular Characterisation

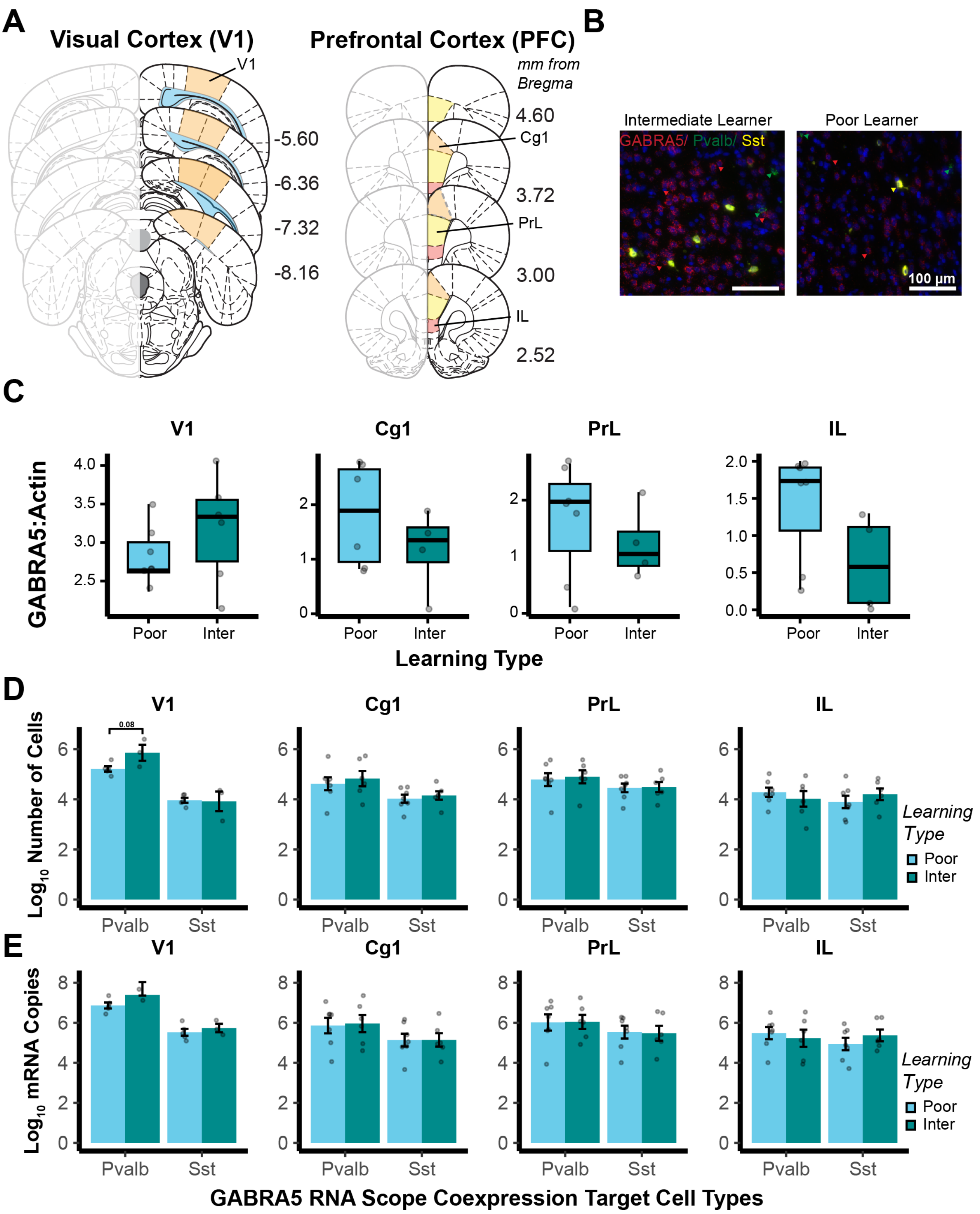

### Supplementary Information

**Figure S9 (overleaf): Female Prefrontal GABA A α5 subunit (GABRA5) mRNA expression levels in inhibitory interneuron cell types correlate with perceptual acuity and learning rate:**
**A** Atlas representations of target regions, left = Primary visual cortex (V1) -5.6mm – -8.16 mm from bregma, right = Prefrontal Cortex (PFC) 4.6 mm – 2.52 mm from bregma. Sub-regions: Anterior Cingulate Cortex (Cg1), Prelimbic Cortex (PrL), Infralimbic Cortex (IL). **B** stratification group sub-region ROI representative images from 2 RNA Scope Fluorescence in-situ-hybridisation assays, scale bar 100 μm. GABRA5 (red) : Pvalb (green) : Sst (yellow). Arrows highlighting cells expressing target probes in each case.
**C** Statistical comparison of Western-immunoblot derived GABRA5 protein expression in cortical ROIs between stratified "Learning types". **D** Cell-wise quantification of GABRA5 co-expression from RNA Scope assays within inhibitory interneuron subtypes: Parvalbumin (Pvalb), Somatostatin (Sst) "Learning type", per target cortical region. **E** Puncta-wise quantification of GABRA5 co-expression from RNA Scope assays within inhibitory interneuron subtypes: Parvalbumin (Pvalb), Somatostatin (Sst) by "Learning type", per target cortical region. mRNA co-expression values displayed for each "Learning type" as mean ± SEM per subject $Log_{10}$ transformed. Comparisons made via Linear mixed-effects model with pairwise comparisons to "Poor Learner" condition each case via estimated-marginal-means, within region and cell type: "p value" = p<0.1, "*" = p<0.05, "**" = p<0.01, "***" = p<0.001.

### Supplementary Information

#### Figure S10: Variation in pharmacological timelines

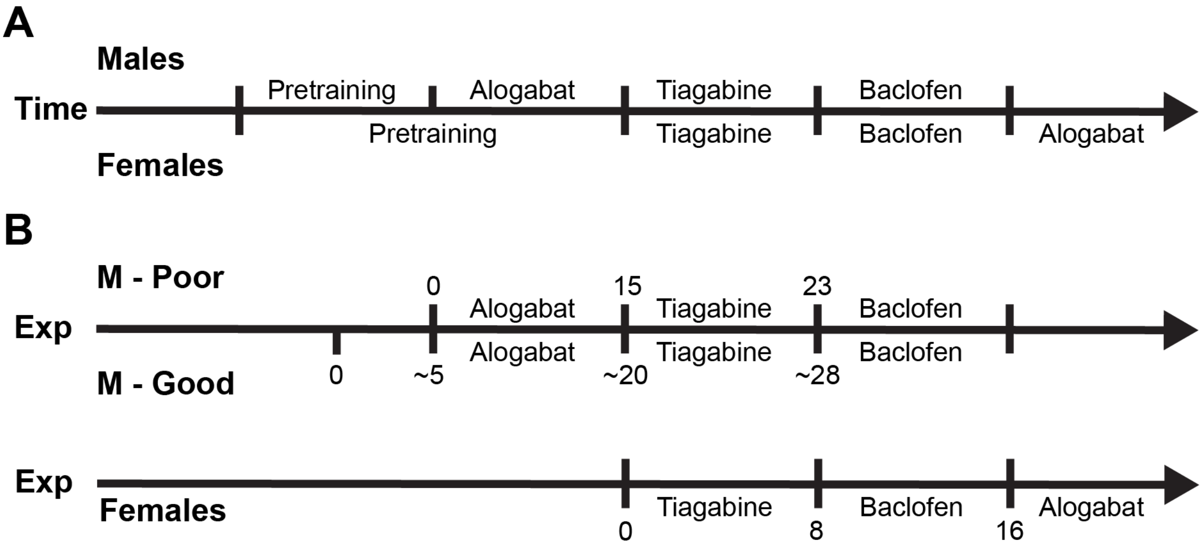


**Figure S10: Female Prefrontal GABA A α5 subunit (GABRA5) mRNA expression levels in inhibitory interneuron cell types correlate with perceptual acuity and learning rate:**
**A** Sex variation in pharmacological schedule by absolute time. **B** Between learning type and within sex variation in pharmacological schedule aligned to absolute time with session numbers of when individual compound latin squares were begun. (~) present in male Good learners as experience was not uniform across this group: 4 IDs had 4 extra baseline sessions, 1 ID had 6 due to rate of task acquisition

### Supplementary Information

#### Figure S11: Baseline performance progression

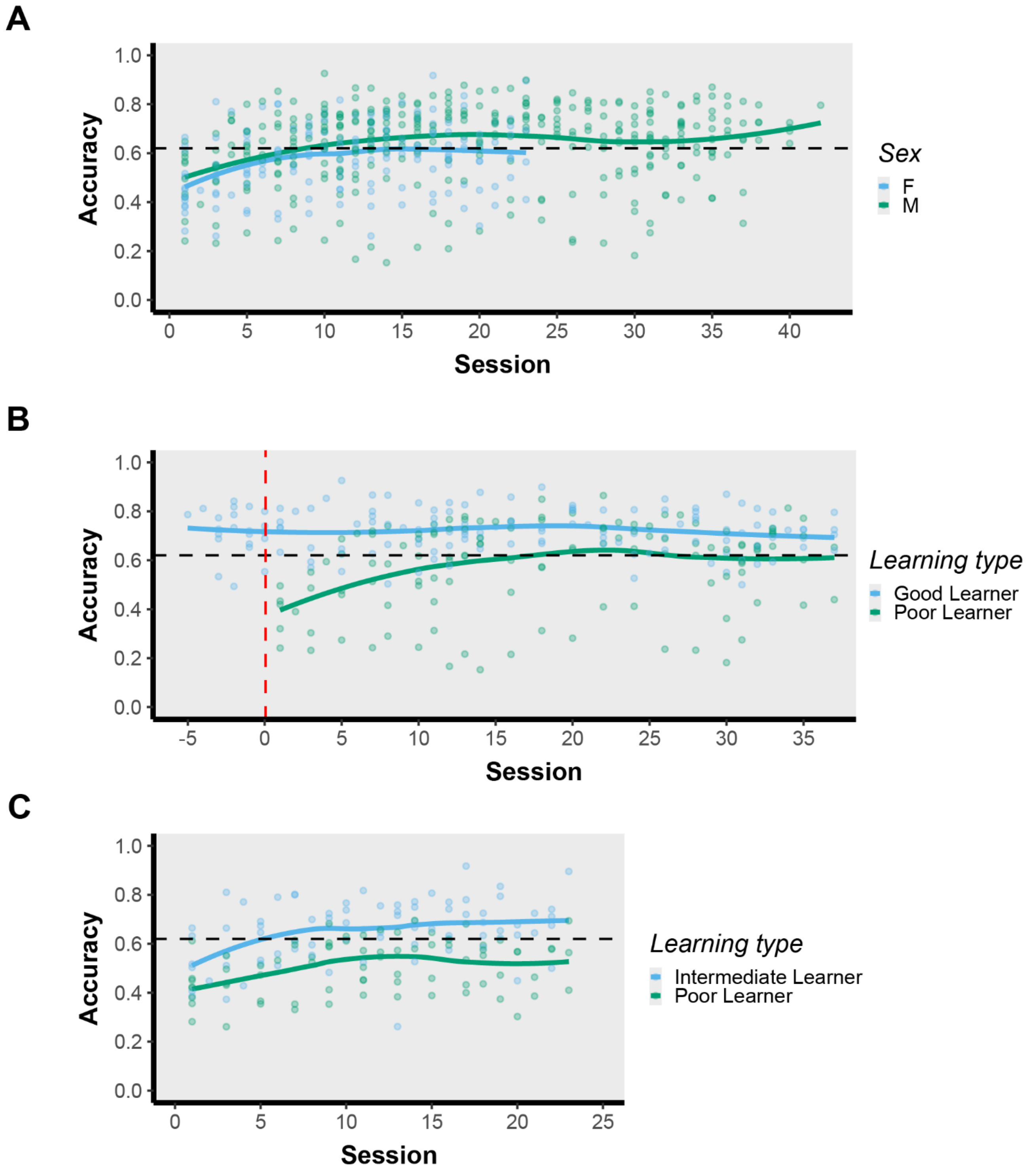


**Figure S11: Session wise baseline accuracy across experiment:**
**A** Sex differences in accuracy aligned to by baseline experience - session. **B** Male learning type stratification differences in accuracy aligned to by baseline experience – session. Red line = session where all animals were moved to baseline task after pretraining. Sessions before this are Good learners only. **C** Female learning type stratification differences in accuracy aligned to by baseline experience – session. No Statistics performed, trend lines present “loess”.

### Supplementary Information

#### Table S1: RNA Scope Probes

| Target | Probe | Source | Identifier |
|---|---|---|---|
| **GABRA5 subunit** | Rn-Gabra5 | Bio-Techne | 411621 |
| **Parvalbumin**-expressing GABAergic interneuron | Rn-Pvalb-C2 | Bio-Techne | 407821-C2 |
| **Somatostatin**-expressing GABAergic interneuron | Rn-Sst | Bio-Techne | 412181-C3 |
| **SLC17A7 (Vesicular glutamate transporter 1)** | Rn-Slc17a7-C2 | Bio-Techne | 317001-C2 |
| **Vasoactive intestinal polypeptide-**expressing GABAergic interneuron | Rn-Vip-C3 | Bio-Techne | 485681-C3 |
| **Positive control probes** | 3-plex Positive Control Probe-Rn | Bio-Techne | 320891 |
| **Negative control probes** | 3-plex Negative Control Probe | Bio-Techne | 320871 |

#### Table S2: Full Statistical Summary
