## Supplementary Table 2 for "Prefrontal α5-GABA-A receptors: putative role in perceptual learning"

| Figure | Outcome | Groups (n) | Obs No. | Test | Reference condition | Random Effect | Fixed Effect Condition | Estimate (SE) | Test Stat: z val, t val or F val | Pr(>\|z\|),Pr(>\|t\|) or Pr(>F) | Panel | post-hoc comparison | Comparison | Estimate (SE) | t/z ratio | Adjusted p value |
|---|---|---|---|---|---|---|---|---|---|---|---|---|---|---|---|---|
| **2** | Accuracy | n=16 - F<br>n=16 - M | 71458 | glmer(logit) | | (1\|id) | (Intercept) | 0.681173 (0.126227) | 5.396 | **6.80E-08** | | | | | | |
| | | | | | ang_sep90 | | ang_sep45 | -0.001682 (0.045105) | -0.037 | 0.970254 | **A** | Sex\|ang_sep | ang_sep90, F-M | -0.143 (0.176) | -0.81 | 0.4181 |
| | | | | | | | ang_sep30 | -0.189659 (0.044631) | -4.249 | **2.14E-05** | | | ang_sep45, F-M | -0.350 (0.176) | -1.985 | **0.0471** |
| | | | | | | | ang_sep15 | -0.598342 (0.045066) | -13.277 | **< 2e-16** | | | ang_sep30, F-M | -0.340 (0.176) | -1.934 | *0.0531* |
| | | | | | | | ang_sep10 | -0.673212 (0.049667) | -13.555 | **< 2e-16** | | | ang_sep15, F-M | -0.388 (0.176) | -2.205 | **0.0275** |
| | | | | | SexF | | SexM | 0.060491 (0.176661) | 0.342 | 0.732042 | | | ang_sep10, F-M | -0.399 (0.177) | -2.246 | **0.0247** |
| | | | | | TDDclose | | TDDfar | -0.239855 (0.030582) | -7.843 | **4.39E-15** | Not marked on figure | ang_sep | ang_sep90 - ang_sep45 | -0.1018 (0.0274) | -3.722 | **0.0018** |
| | | | | | ang_sep90, SexF | | ang_sep45:SexM | 0.207032 (0.054720) | 3.783 | **0.000155** | | | ang_sep90 - ang_sep30 | 0.0909 (0.0270) | 3.364 | **0.0069** |
| | | | | | | | ang_sep30:SexM | 0.197574 (0.054023) | 3.657 | **0.000255** | | | ang_sep90 - ang_sep15 | 0.4757 (0.0271) | 17.561 | **<.0001** |
| | | | | | | | ang_sep15:SexM | 0.245335 (0.054174) | 4.529 | **5.94E-06** | | | ang_sep90 - ang_sep10 | 0.5452 (0.0293) | 18.616 | **<.0001** |
| | | | | | | | ang_sep10:SexM | 0.255954 (0.058578) | 4.369 | **1.25E-05** | | | ang_sep45 - ang_sep30 | 0.1927 (0.0268) | 7.185 | **<.0001** |
| | | | | | TDDclose, SexF | | SexM:TDDfar | 0.164507 (0.036754) | 4.476 | **7.61E-06** | | | ang_sep45 - ang_sep15 | 0.5775 (0.0269) | 21.468 | **<.0001** |
| | | | | | | | | | | | | | ang_sep45 - ang_sep10 | 0.6471 (0.0291) | 22.236 | **<.0001** |
| | | | | | | | | | | | | | ang_sep30 - ang_sep15 | 0.3848 (0.0265) | 14.502 | **<.0001** |
| | | | | | | | | | | | | | ang_sep30 - ang_sep10 | 0.4544 (0.0288) | 15.796 | **<.0001** |
| | | | | | | | | | | | | | ang_sep15 - ang_sep10 | 0.0696 (0.0288) | 2.419 | 0.1103 |
| | | | | | | | | | | | **B** | Sex | Taken from main effects | 0.0604 (0.1766) | 0.342 | 0.732 |
| | | | | | | | | | | | **C** | Sex\|TDD | TDDclose, F-M | -0.242 (0.173) | -1.395 | 0.1629 |
| | | | | | | | | | | | | | TDDfar, F-M | -0.406 (0.174) | -2.328 | **0.0199** |
| | | | | | | | | | | | | TDD\|Sex | F, TDDclose - TDDfar | 0.2399 (0.0306) | 7.843 | **<.0001** |
| | | | | | | | | | | | | | M, TDDclose - TDDfar | 0.0753 (0.0204) | 3.696 | **0.0002** |
| | InverseRT | n=16 - F<br>n=15 - M | 310 | lmer | | (1\|id) | (Intercept) | -0.602045 (0.029008) | -20.755 | **< 2e-16** | | | | | | |
| | | | | | ang_sep90 | | ang_sep45 | 0.052477 (0.007656) | 6.854 | **4.89E-11** | **D** | Sex\|ang_sep | ang_sep90, F-M | -0.00805 (0.0416) | -0.194 | 0.8477 |
| | | | | | | | ang_sep30 | 0.066383 (0.007656) | 8.67 | **4.14E-16** | | | ang_sep45, F-M | -0.03836 (0.0416) | -0.923 | 0.3632 |
| | | | | | | | ang_sep15 | 0.063258 (0.007656) | 8.262 | **6.57E-15** | | | ang_sep30, F-M | -0.05390 (0.0416) | -1.297 | 0.2043 |
| | | | | | | | ang_sep10 | 0.073152 (0.007656) | 9.554 | **< 2e-16** | | | ang_sep15, F-M | -0.07815 (0.0416) | -1.881 | *0.0696* |
| | | | | | SexF | | SexM | 0.006402 (0.041701) | 0.154 | 0.87898 | | | ang_sep10, F-M | -0.07573 (0.0416) | -1.822 | *0.0781* |
| | | | | | TDDclose | | stim_dist_locfar | 0.012270 (0.004842) | 2.534 | **0.01185** | Not marked on figure | ang_sep | ang_sep90 - ang_sep45 | -0.06763 (0.0055) | -12.289 | **<.0001** |
| | | | | | ang_sep90, SexF | | ang_sep45:SexM | 0.030312 (0.011007) | 2.754 | **0.00629** | | | ang_sep90 - ang_sep30 | -0.08931 (0.0055) | -16.228 | **<.0001** |
| | | | | | | | ang_sep30:SexM | 0.045854 (0.011007) | 4.166 | **4.18E-05** | | | ang_sep90 - ang_sep15 | -0.09831 (0.0055) | -17.863 | **<.0001** |
| | | | | | | | ang_sep15:SexM | 0.070098 (0.011007) | 6.369 | **8.21E-10** | | | ang_sep90 - ang_sep10 | -0.10699 (0.0055) | -19.442 | **<.0001** |
| | | | | | | | ang_sep10:SexM | 0.067684 (0.011007) | 6.149 | **2.80E-09** | | | ang_sep45 - ang_sep30 | -0.02168 (0.0055) | -3.939 | **0.001** |
| | | | | | TDDclose, SexF | | SexM:stim_dist_locfar | 0.003292 (0.006961) | 0.473 | 0.63665 | | | ang_sep45 - ang_sep15 | -0.03067 (0.0055) | -5.574 | **<.0001** |
| | | | | anova(lmer) | | | ang_sep | | F(4,269) = 122.4680 | **< 2.2e-16** | | | ang_sep45 - ang_sep10 | -0.03936 (0.0055) | -7.152 | **<.0001** |
| | | | | | | | Sex | | F(1, 29) = 1.5398 | 0.2246 | | | ang_sep30 - ang_sep15 | -0.00900 (0.0055) | -1.635 | 0.4764 |
| | | | | | | | TDD | | F(1, 269) =15.9859 | **8.25E-05** | | | ang_sep30 - ang_sep10 | -0.01768 (0.0055) | -3.213 | **0.0127** |
| | | | | | | | ang_sep:Sex | | F(4, 269) =13.8737 | **2.66E-10** | | | ang_sep15 - ang_sep10 | -0.00869 (0.0055) | -1.578 | 0.5125 |
| | | | | | | | Sex:TDD | | F(1, 269) = 0.2237 | 0.6366 | **E** | Sex | Taken from main effects, anova(lmer) | | | |
| | | | | | | | | | | | **F** | Sex\|TDD | No significant main effect *(anova(lmer))* so pairwise not performed | | | |
| **3** | Accuracy | n=6 - Poor<br>n=5 - Good | 33594 | glmer(logit) | | (1\|id) | (Intercept) | 0.45254 (0.18320) | 2.47 | **0.0135** | | | | | | |
| | | | | | ang_sep90 | | ang_sep45 | 0.24456 (0.05015) | 4.876 | **1.08E-06** | **B** | LT\|ang_sep | ang_sep90, Poor-Good | -0.650 (0.272) | -2.395 | **0.0166** |
| | | | | | | | ang_sep30 | 0.05325 (0.04958) | 1.074 | 0.2828 | | | ang_sep45, Poor-Good | -0.752 (0.272) | -2.765 | **0.0057** |
| | | | | | | | ang_sep15 | -0.32351 (0.05005) | -6.463 | **1.02E-10** | | | ang_sep30, Poor-Good | -0.716 (0.272) | -2.639 | ***0.0083*** |
| | | | | | | | ang_sep10 | -0.24510 (0.05490) | -4.464 | **8.04E-06** | | | ang_sep15, Poor-Good | -0.703 (0.271) | -2.592 | **0.0095** |
| | | | | | LT_Poor | | LT_Good | 0.60797 (0.27194) | 2.236 | **0.0253** | | | ang_sep10, Poor-Good | -0.374 (0.272) | -1.376 | 0.1688 |

| | | | | | | | | | | | | | | | | |
|---|---|---|---|---|---|---|---|---|---|---|---|---|---|---|---|---|
| | | | | | TDDclose | | TDDfar | -0.15374 (0.03437) | -4.474 | **7.69E-06** | Not marked on figure | ang_sep | ang_sep90 - ang_sep45 | -0.2955 (0.0385) | -7.668 | **<.0001** |
| | | | | | ang_sep90,LT_Poor | | ang_sep45:LT_Good | 0.10190 (0.07707) | 1.322 | 0.1861 | | | ang_sep90 - ang_sep30 | -0.0864 (0.0376) | -2.295 | 0.1463 |
| | | | | | | | ang_sep30:LT_Good | 0.06621 (0.07525) | 0.88 | 0.3789 | | | ang_sep90 - ang_sep15 | 0.2971 (0.0370) | 8.027 | **<.0001** |
| | | | | | | | ang_sep15:LT_Good | 0.05284 (0.07403) | 0.714 | 0.4753 | | | ang_sep90 - ang_sep10 | 0.3829 (0.0386) | 9.93 | **<.0001** |
| | | | | | | | ang_sep10:LT_Good | -0.27569 (0.07713) | -3.575 | **3.51E-04** | | | ang_sep45 - ang_sep30 | 0.2092 (0.0385) | 5.429 | **<.0001** |
| | | | | | TDDclose, LT_Poor | | LT_Good :TDDfar | 0.08440 (0.05039) | 1.675 | *0.0939* | | | ang_sep45 - ang_sep15 | 0.5926 (0.0379) | 15.621 | **<.0001** |
| | | | | | | | | | | | | | ang_sep45 - ang_sep10 | 0.6785 (0.0395) | 17.198 | **<.0001** |
| | | | | | | | | | | | | | ang_sep30 - ang_sep15 | 0.3834 (0.0370) | 10.363 | **<.0001** |
| | | | | | | | | | | | | | ang_sep30 - ang_sep10 | 0.4693 (0.0385) | 12.174 | **<.0001** |
| | | | | | | | | | | | | | ang_sep15 - ang_sep10 | 0.0858 (0.0378) | 2.27 | 0.1545 |
| | | | | | | | | | | | **C** | LT | Taken from main effects | 0.60797 (0.27194) | 2.236 | **0.0253** |
| | | | | | | | | | | | **D** | LT\|TDD | TDDclose, Poor-Good | -0.597 (0.268) | -2.229 | **0.0258** |
| | | | | | | | | | | | | | TDDfar, Poor-Good | -0.681 (0.269) | -2.53 | **0.0114** |
| | | | | | | | | | | | | TDD\|LT | Poor, TDDclose - TDDfar | 0.1537 (0.0344) | 4.474 | **<.0001** |
| | | | | | | | | | | | | | Good, TDDclose - TDDfar | 0.0693 (0.0368) | 1.882 | *0.0599* |
| | InverseRT | n=6 - Poor<br>n=5 - Good | 110 | lmer | | (1\|id) | (Intercept) | -0.564611 (0.046894) | -12.04 | **3.73E-07** | | | | | | |
| | | | | | ang_sep90 | | ang_sep45 | 0.095060 (0.012821) | 7.414 | **6.80E-11** | **E** | LT\|ang_sep | ang_sep90, Poor-Good | 0.0825 (0.0693) | 1.191 | 0.2625 |
| | | | | | | | ang_sep30 | 0.135510 (0.012821) | 10.569 | **< 2e-16** | | | ang_sep45, Poor-Good | 0.0948 (0.0693) | 1.368 | 0.2027 |
| | | | | | | | ang_sep15 | 0.154253 (0.012821) | 12.031 | **< 2e-16** | | | ang_sep30, Poor-Good | 0.1029 (0.0693) | 1.484 | 0.1699 |
| | | | | | | | ang_sep10 | 0.164226 (0.012821) | 12.809 | **< 2e-16** | | | ang_sep15, Poor-Good | 0.0944 (0.0693) | 1.362 | 0.2044 |
| | | | | | LT_Poor | | LT_Good | -0.086996 (0.069555) | -1.251 | 0.24 | | | ang_sep10, Poor-Good | 0.0842 (0.0693) | 1.214 | 0.2537 |
| | | | | | TDDclose | | TDDfar | 0.011268 (0.008109) | 1.39 | 0.168 | Not marked on figure | ang_sep | ang_sep90 - ang_sep45 | -0.0889 (0.00951) | -9.352 | **<.0001** |
| | | | | | ang_sep90,LT_Poor | | ang_sep45:LT_Good | -0.012267 (0.019017) | -0.645 | 0.521 | | | ang_sep90 - ang_sep30 | -0.1253 (0.00951) | -13.181 | **<.0001** |
| | | | | | | | ang_sep30:LT_Good | -0.020351 (0.019017) | -1.07 | 0.287 | | | ang_sep90 - ang_sep15 | -0.1483 (0.00951) | -15.597 | **<.0001** |
| | | | | | | | ang_sep15:LT_Good | -0.011886 (0.019017) | -0.625 | 0.534 | | | ang_sep90 - ang_sep10 | -0.1634 (0.00951) | -17.185 | **<.0001** |
| | | | | | | | ang_sep10:LT_Good | -0.001648 (0.019017) | -0.087 | 0.931 | | | ang_sep45 - ang_sep30 | -0.0364 (0.00951) | -3.829 | **0.0022** |
| | | | | | TDDclose, LT_Poor | | LT_Good :TDDfar | 0.008983 (0.012028) | 0.747 | 0.457 | | | ang_sep45 - ang_sep15 | -0.0594 (0.00951) | -6.245 | **<.0001** |
| | | | | anova(lmer) | | | ang_sep | | F(4,89) = 93.9198 | **< 2e-16** | | | ang_sep45 - ang_sep10 | -0.0745 (0.00951) | -7.833 | **<.0001** |
| | | | | | | | LT | | F(1, 9) = 1.8070 | 0.2118 | | | ang_sep30 - ang_sep15 | -0.0230 (0.00951) | -2.416 | 0.1205 |
| | | | | | | | TDD | | F(1, 89) =6.8677 | **0.01032** | | | ang_sep30 - ang_sep10 | -0.0381 (0.00951) | -4.004 | **0.0012** |
| | | | | | | | ang_sep:LT | | F(4, 89) =0.3908 | 0.8148 | | | ang_sep15 - ang_sep10 | -0.0151 (0.00951) | -1.587 | 0.5094 |
| | | | | | | | LT:TDD | | F(1, 89) = 0.5578 | 0.4571 | **F** | LT | Taken from main effects | -0.086996 (0.069555) | -1.251 | 0.24 |
| | | | | | | | | | | | **G** | LT\|TDD | No significant main effect *(anova(lmer))* so pairwise not performed | | | |
| **4** | Accuracy | n=5 - Poor<br>n=5 - Good | 5238 | glmer(logit) | | (1\|id) | (Intercept) | 0.633992 (0.317722) | 1.995 | **0.046** | | | | | | |
| | *Tiagabine* | | | | ang_sep90 | | ang_sep45 | 0.456354 (0.104951) | 4.348 | **1.37E-05** | **A** | Dose | Veh - 0.5 | 0.015257 (0.0982) | 0.155 | 0.9987 |
| | | | | | | | ang_sep30 | 0.166744 (0.101390) | 1.645 | 0.1001 | | | Veh - 2.5 | 0.015992 (0.0929) | 0.172 | 0.9982 |
| | | | | | | | ang_sep15 | -0.227918 (0.098609) | -2.311 | **0.0208** | | | Veh - 7.5 | -0.047110 (0.0936) | -0.503 | 0.9583 |
| | | | | | | | ang_sep10 | -0.630371 (0.101659) | -6.201 | **5.62E-10** | | | 0.5 - 2.5 | 0.000735 (0.0925) | 0.008 | 1 |

| | | | | | | | | | |
|---|---|---|---|---|---|---|---|---|---|
| Dose_Veh | Dose_0.5 | 0.028341 (0.132578) | 0.214 | 0.8307 | | 0.5 - 7.5 | -0.062367 (0.0941) | -0.663 | 0.911 |
| | Dose_2.5 | -0.095466 (0.129351) | -0.738 | 0.4605 | | 2.5 - 7.5 | -0.063101 (0.0887) | -0.711 | 0.8926 |
| | Dose_7.5 | -0.006305 (0.129058) | -0.049 | 0.961 | Dose\|LT | LT_Poor : Veh - 0.5 | -0.0283 (0.133) | -0.214 | 0.9966 |
| LT_Poor | LT_Good | 0.695795 (0.441251) | 1.577 | 0.1148 | | LT_Poor : Veh - 2.5 | 0.0955 (0.129) | 0.738 | 0.8818 |
| Dose_Veh, LT_Poor | Dose_0.5: LT_Good | -0.087195 (0.196310) | -0.444 | 0.6569 | | LT_Poor : Veh - 7.5 | 0.0063 (0.129) | 0.049 | 1 |
| | Dose_2.5: LT_Good | 0.158950 (0.185578) | 0.857 | 0.3917 | | LT_Poor : 0.5 - 2.5 | 0.1238 (0.121) | 1.026 | 0.734 |
| | Dose_7.5: LT_Good | 0.106829 (0.187524) | 0.57 | 0.5689 | | LT_Poor : 0.5 - 7.5 | 0.0346 (0.120) | 0.288 | 0.9917 |
| | | | | | | LT_Poor : 2.5 - 7.5 | -0.0892 (0.117) | -0.763 | 0.8711 |
| | | | | | | LT_Good : Veh - 0.5 | 0.0589 (0.145) | 0.406 | 0.9774 |
| | | | | | | LT_Good : Veh - 2.5 | -0.0635 (0.133) | -0.477 | 0.9642 |
| | | | | | | LT_Good : Veh - 7.5 | -0.1005 (0.136) | -0.74 | 0.8808 |
| | | | | | | LT_Good : 0.5 - 2.5 | -0.1223 (0.140) | -0.872 | 0.8192 |
| | | | | | | LT_Good : 0.5 - 7.5 | -0.1594 (0.145) | -1.1 | 0.6898 |
| | | | | | | LT_Good : 2.5 - 7.5 | -0.0370 (0.134) | -0.277 | 0.9926 |
| | | | | | Dose*LT | Veh Poor Learner - 0.5 Poor Learner | -0.0283 (0.133) | -0.214 | 1 |
| | | | | | | Veh Poor Learner - 2.5 Poor Learner | 0.0955 (0.129) | 0.738 | 0.9958 |
| | | | | | | Veh Poor Learner - 7.5 Poor Learner | 0.0063 (0.129) | 0.049 | 1 |
| | | | | | | Veh Poor Learner - Veh Good Learner | -0.6958 (0.441) | -1.577 | 0.7644 |
| | | | | | | Veh Poor Learner - 0.5 Good Learner | -0.6369 (0.442) | -1.441 | 0.8384 |
| | | | | | | Veh Poor Learner - 2.5 Good Learner | -0.7593 (0.440) | -1.727 | 0.6692 |
| | | | | | | Veh Poor Learner - 7.5 Good Learner | -0.7963 (0.441) | -1.807 | 0.6156 |
| | | | | | | 0.5 Poor Learner - 2.5 Poor Learner | 0.1238 (0.121) | 1.026 | 0.9706 |
| | | | | | | 0.5 Poor Learner - 7.5 Poor Learner | 0.0346 (0.120) | 0.288 | 1 |
| | | | | | | 0.5 Poor Learner - Veh Good Learner | -0.6675 (0.439) | -1.522 | 0.7959 |
| | | | | | | 0.5 Poor Learner - 0.5 Good Learner | -0.6086 (0.439) | -1.385 | 0.8644 |
| | | | | | | 0.5 Poor Learner - 2.5 Good Learner | -0.7309 (0.437) | -1.673 | 0.7047 |
| | | | | | | 0.5 Poor Learner - 7.5 Good Learner | -0.7680 (0.438) | -1.753 | 0.6521 |
| | | | | | | 2.5 Poor Learner - 7.5 Poor Learner | -0.0892 (0.117) | -0.763 | 0.9949 |
| | | | | | | 2.5 Poor Learner - Veh Good Learner | -0.7913 (0.438) | -1.808 | 0.6147 |
| | | | | | | 2.5 Poor Learner - 0.5 Good Learner | -0.7324 (0.439) | -1.67 | 0.7067 |
| | | | | | | 2.5 Poor Learner - 2.5 Good Learner | -0.8547 (0.436) | -1.961 | 0.5087 |
| | | | | | | 2.5 Poor Learner - 7.5 Good Learner | -0.8918 (0.437) | -2.04 | 0.4549 |
| | | | | | | 7.5 Poor Learner - Veh Good Learner | -0.7021 (0.438) | -1.605 | 0.7477 |
| | | | | | | 7.5 Poor Learner - 0.5 Good Learner | -0.6432 (0.438) | -1.467 | 0.8249 |
| | | | | | | 7.5 Poor Learner - 2.5 Good Learner | -0.7656 (0.436) | -1.757 | 0.6495 |
| | | | | | | 7.5 Poor Learner - 7.5 Good Learner | -0.8026 (0.437) | -1.836 | 0.5952 |
| | | | | | | Veh Good Learner - 0.5 Good Learner | 0.0589 (0.145) | 0.406 | 0.9999 |
| | | | | | | Veh Good Learner - 2.5 Good Learner | -0.0635 (0.133) | -0.477 | 0.9998 |
| | | | | | | Veh Good Learner - 7.5 Good Learner | -0.1005 (0.136) | -0.74 | 0.9958 |
| | | | | | | 0.5 Good Learner - 2.5 Good Learner | -0.1223 (0.140) | -0.872 | 0.9885 |

| | | | | | | | | | | | | | | | |
|---|---|---|---|---|---|---|---|---|---|---|---|---|---|---|---|
| | | | | | | | | | | | | 0.5 Good Learner - 7.5 Good Learner | -0.1594 (0.145) | -1.1 | 0.9571 |
| | | | | | | | | | | | | 2.5 Good Learner - 7.5 Good Learner | -0.0370 (0.134) | -0.277 | 1 |
| InverseRT | n=5 - Poor n=5 - Good | 181 | lmer | | (1\|id) | (Intercept) | -0.556715 (0.045773) | -12.162 | **2.83E-07** | | | | | | |
| *Tiagabine* | | | | ang_sep90 | | ang_sep45 | 0.042120 (0.013324) | 3.161 | **1.88E-03** | **B** | Dose | Veh - 0.5 | 0.0334 0.0131 | 2.559 | *0.0549* |
| | | | | | | ang_sep30 | 0.060750 (0.013324) | 4.56 | **1.01E-05** | | | Veh - 2.5 | 0.0440 0.0125 | 3.505 | **0.0033** |
| | | | | | | ang_sep15 | 0.081818 (0.013324) | 6.141 | **6.18E-09** | | | Veh - 7.5 | -0.0324 0.0125 | -2.595 | *0.0501* |
| | | | | | | ang_sep10 | 0.104171 (0.013756) | 7.573 | **2.69E-12** | | | 0.5 - 2.5 | 0.0105 0.0121 | 0.875 | 0.8178 |
| | | | | Dose_Veh | | Dose_0.5 | -0.015345 (0.017949) | -0.855 | 0.39385 | | | 0.5 - 7.5 | -0.0658 0.0121 | -5.457 | **<.0001** |
| | | | | | | Dose_2.5 | -0.010941 (0.017927) | 0.61 | 0.54254 | | | 2.5 - 7.5 | -0.0764 0.0116 | -6.592 | **<.0001** |
| | | | | | | Dose_7.5 | 0.016051 (0.017763) | 0.904 | 0.36755 | | Dose\|LT | LT_Poor : Veh - 0.5 | 0.0153 (0.0180) | 0.855 | 0.828 |
| | | | | LT_Poor | | LT_Good | -0.006855 (0.063584) | -0.108 | 0.91646 | | | LT_Poor : Veh - 2.5 | 0.0109 (0.0179) | 0.61 | 0.9287 |
| | | | | Dose_Veh, LT_Poor | | Dose_0.5: LT_Good | -0.036123 (0.026093) | -1.384 | 0.16814 | | | LT_Poor : Veh - 7.5 | -0.0161 (0.0178) | -0.903 | 0.803 |
| | | | | | | Dose_2.5: LT_Good | -0.066025 (0.025072) | -2.633 | **0.00928** | | | LT_Poor : 0.5 - 2.5 | -0.0044 (0.0166) | -0.266 | 0.9934 |
| | | | | | | Dose_7.5: LT_Good | 0.032761 (0.024999) | 1.311 | 0.19188 | | | LT_Poor : 0.5 - 7.5 | -0.0314 (0.0164) | -1.916 | 0.2255 |
| | | | anova(lmer) | | | LT | | F(1,8) = 0.1558 | 0.703362 | | | LT_Poor : 2.5 - 7.5 | -0.0270 (0.0164) | -1.647 | 0.3555 |
| | | | | | | Dose | | F(3,161) = 17.280 | **8.88E-10** | | | LT_Good : Veh - 0.5 | 0.0515 (0.0190) | 2.716 | **0.0365** |
| | | | | | | ang_sep | | F(4,161) = 17.084 | **1.07E-11** | | | LT_Good : Veh - 2.5 | 0.0770 (0.0175) | 4.39 | **0.0001** |
| | | | | | | Dose:LT | | F(3,161) = 6.680 | **2.81E-04** | | | LT_Good : Veh - 7.5 | -0.0488 (0.0176) | -2.775 | **0.031** |
| | | | | | | | | | | | | LT_Good : 0.5 - 2.5 | 0.0255 (0.0175). | 1.456 | 0.4667 |
| | | | | | | | | | | | | LT_Good : 0.5 - 7.5 | -0.1003 (0.0177) | -5.659 | **<.0001** |
| | | | | | | | | | | | | LT_Good : 2.5 - 7.5 | -0.1258 (0.0164) | -7.673 | **<.0001** |
| | | | | | | | | | | | Dose*LT | Veh Poor Learner - 0.5 Poor Learner | 0.01535 (0.0180) | 0.855 | 0.9895 |
| | | | | | | | | | | | | Veh Poor Learner - 2.5 Poor Learner | 0.01094 (0.0179) | 0.61 | 0.9987 |
| | | | | | | | | | | | | Veh Poor Learner - 7.5 Poor Learner | -0.01605 (0.0178) | -0.903 | 0.9854 |
| | | | | | | | | | | | | Veh Poor Learner - Veh Good Learner | 0.00685 (0.0636) | 0.108 | 1 |
| | | | | | | | | | | | | Veh Poor Learner - 0.5 Good Learner | 0.05832 (0.0636) | 0.917 | 0.9764 |
| | | | | | | | | | | | | Veh Poor Learner - 2.5 Good Learner | 0.08382 (0.0632) | 1.326 | 0.8685 |
| | | | | | | | | | | | | Veh Poor Learner - 7.5 Good Learner | -0.04196 (0.0633) | -0.663 | 0.9962 |
| | | | | | | | | | | | | 0.5 Poor Learner - 2.5 Poor Learner | -0.00440 (0.0166) | -0.266 | 1 |
| | | | | | | | | | | | | 0.5 Poor Learner - 7.5 Poor Learner | -0.03140 (0.0164) | -1.916 | 0.5423 |
| | | | | | | | | | | | | 0.5 Poor Learner - Veh Good Learner | -0.00849 (0.0632) | -0.134 | 1 |
| | | | | | | | | | | | | 0.5 Poor Learner - 0.5 Good Learner | 0.04298 (0.0632) | 0.68 | 0.9956 |
| | | | | | | | | | | | | 0.5 Poor Learner - 2.5 Good Learner | 0.06848 (0.0629) | 1.089 | 0.9438 |
| | | | | | | | | | | | | 0.5 Poor Learner - 7.5 Good Learner | -0.05730 (0.0629) | -0.911 | 0.977 |
| | | | | | | | | | | | | 2.5 Poor Learner - 7.5 Poor Learner | -0.02699 (0.0164) | -1.647 | 0.7211 |
| | | | | | | | | | | | | 2.5 Poor Learner - Veh Good Learner | -0.00409 (0.0632) | -0.065 | 1 |
| | | | | | | | | | | | | 2.5 Poor Learner - 0.5 Good Learner | 0.04738 (0.0632) | 0.75 | 0.9922 |
| | | | | | | | | | | | | 2.5 Poor Learner - 2.5 Good Learner | 0.07288 (0.0629) | 1.159 | 0.9251 |
| | | | | | | | | | | | | 2.5 Poor Learner - 7.5 Good Learner | -0.05290 (0.0629) | -0.841 | 0.985 |
| | | | | | | | | | | | | 7.5 Poor Learner - Veh Good Learner | 0.02291 (0.0632) | 0.363 | 0.9999 |

| | | | | | | | | | | | | | | | |
|---|---|---|---|---|---|---|---|---|---|---|---|---|---|---|---|
| | | | | | | | | | | | | 7.5 Poor Learner - 0.5 Good Learner | 0.07437 (0.0632) | 1.177 | 0.9201 |
| | | | | | | | | | | | | 7.5 Poor Learner - 2.5 Good Learner | 0.09987 (0.0628) | 1.59 | 0.7465 |
| | | | | | | | | | | | | 7.5 Poor Learner - 7.5 Good Learner | -0.02591 (0.0629) | -0.412 | 0.9998 |
| | | | | | | | | | | | | Veh Good Learner - 0.5 Good Learner | 0.05147 (0.0190) | 2.716 | 0.1253 |
| | | | | | | | | | | | | Veh Good Learner - 2.5 Good Learner | 0.07697 (0.0175) | 4.39 | **0.0005** |
| | | | | | | | | | | | | Veh Good Learner - 7.5 Good Learner | -0.04881 (0.0176) | -2.775 | 0.1086 |
| | | | | | | | | | | | | 0.5 Good Learner - 2.5 Good Learner | 0.02550 (0.0175) | 1.456 | 0.8294 |
| | | | | | | | | | | | | 0.5 Good Learner - 7.5 Good Learner | -0.10028 (0.0177) | -5.659 | **<.0001** |
| | | | | | | | | | | | | 2.5 Good Learner - 7.5 Good Learner | -0.12578 (0.0164) | -7.673 | **<.0001** |
| Accuracy | n=5 - Poor n=5 - Good | 4660 | glmer(logit) | | (1\|id) | (Intercept) | 0.45198 (0.23550) | 1.919 | *0.054956* | | | | | | |
| Baclofen | | | | ang_sep90 | | ang_sep45 | 0.42851 (0.11184) | 3.832 | **1.27E-04** | **C** | Dose | Veh - 0.05 | -0.137 0.0845 | -1.623 | 0.3655 |
| | | | | | | ang_sep30 | 0.33062 (0.11032) | 2.997 | **0.002727** | | | Veh - 0.5 | -0.409 0.0863 | -4.742 | **<.0001** |
| | | | | | | ang_sep15 | -0.12539 (0.10522) | -1.192 | 0.233415 | | | Veh - 1.5 | -0.634 0.2060 | *-3.076* | **0.0113** |
| | | | | | | ang_sep10 | -0.40979 (0.10506) | -3.901 | **9.60E-05** | | | 0.05 - 0.5 | -0.272 0.0881 | -3.087 | **0.0109** |
| | | | | Dose_Veh | | Dose_0.05 | -0.01975 (0.11370) | -0.174 | 0.86206 | | | 0.05 - 1.5 | -0.496 0.2060 | -2.405 | *0.0761* |
| | | | | | | Dose_0.5 | 0.49194 (0.11763) | 4.182 | **2.89E-05** | | | 0.5 - 1.5 | -0.224 0.2070 | -1.083 | 0.7002 |
| | | | | | | Dose_1.5 | 1.07488 (0.34479) | 3.118 | **0.001824** | | Dose\|LT | LT_Poor : Veh - 0.05 | 0.0198 (0.114) | 0.174 | 0.9981 |
| | | | | LT_Poor | | LT_Good | 0.58159 (0.32048) | 1.815 | *0.069564* | | | LT_Poor : Veh - 0.5 | -0.4919 (0.118) | -4.182 | **0.0002** |
| | | | | Dose_Veh, LT_Poor | | Dose_0.05: LT_Good | 0.31389 (0.16906) | 1.857 | *0.063366* | | | LT_Poor : Veh - 1.5 | -1.0749 (0.345) | -3.118 | **0.0099** |
| | | | | | | Dose_0.5: LT_Good | -0.16582 (0.17249) | -0.961 | 0.336385 | | | LT_Poor : 0.05 - 0.5 | -0.5117 (0.118) | -4.339 | **0.0001** |
| | | | | | | Dose_1.5: LT_Good | -0.88274 (0.41232) | -2.141 | **0.032279** | | | LT_Poor : 0.05 - 1.5 | -1.0946 (0.344) | -3.182 | **0.008** |
| | | | | | | | | | | | | LT_Poor : 0.5 - 1.5 | -0.5829 (0.346) | -1.683 | 0.3327 |
| | | | | | | | | | | | | LT_Good : Veh - 0.05 | -0.2941 (0.125) | -2.351 | *0.0868* |
| | | | | | | | | | | | | LT_Good : Veh - 0.5 | -0.3261 (0.126) | -2.585 | **0.048** |
| | | | | | | | | | | | | LT_Good : Veh - 1.5 | -0.1921 (0.226) | -0.851 | 0.83 |
| | | | | | | | | | | | | LT_Good : 0.05 - 0.5 | -0.0320 (0.131) | -0.244 | 0.9949 |
| | | | | | | | | | | | | LT_Good : 0.05 - 1.5 | 0.1020 (0.228) | 0.447 | 0.9702 |
| | | | | | | | | | | | | LT_Good : 0.5 - 1.5 | 0.1340 (0.228) | 0.587 | 0.9362 |
| | | | | | | | | | | | Dose*LT | Veh Poor Learner - 0.05 Poor Learner | 0.0198 (0.114) | 0.174 | 1 |
| | | | | | | | | | | | | Veh Poor Learner - 0.5 Poor Learner | -0.4919 (0.118) | -4.182 | **0.0008** |
| | | | | | | | | | | | | Veh Poor Learner - 1.5 Poor Learner | -1.0749 (0.345) | -3.118 | **0.0386** |
| | | | | | | | | | | | | Veh Poor Learner - Veh Good Learner | -0.5816 (0.320) | -1.815 | 0.61 |
| | | | | | | | | | | | | Veh Poor Learner - 0.05 Good Learner | -0.8757 (0.322) | -2.716 | 0.1176 |
| | | | | | | | | | | | | Veh Poor Learner - 0.5 Good Learner | -0.9077 (0.323) | -2.812 | 0.0919 |
| | | | | | | | | | | | | Veh Poor Learner - 1.5 Good Learner | -0.7737 (0.374) | -2.07 | 0.4347 |
| | | | | | | | | | | | | 0.05 Poor Learner - 0.5 Poor Learner | -0.5117 (0.118) | -4.339 | **0.0004** |
| | | | | | | | | | | | | 0.05 Poor Learner - 1.5 Poor Learner | -1.0946 (0.344) | -3.182 | **0.0316** |
| | | | | | | | | | | | | 0.05 Poor Learner - Veh Good Learner | -0.6013 (0.321) | -1.876 | 0.5676 |
| | | | | | | | | | | | | 0.05 Poor Learner - 0.05 Good Learner | -0.8955 (0.322) | -2.777 | 0.1008 |
| | | | | | | | | | | | | 0.05 Poor Learner - 0.5 Good Learner | -0.9275 (0.323) | -2.872 | 0.0782 |

| | | | | | | | | | | | | | | | |
|---|---|---|---|---|---|---|---|---|---|---|---|---|---|---|---|
| | | | | | | | | | | | | 0.05 Poor Learner - 1.5 Good Learner | -0.7935 (0.374) | -2.122 | 0.4004 |
| | | | | | | | | | | | | 0.5 Poor Learner - 1.5 Poor Learner | -0.5829 (0.346) | -1.683 | 0.6986 |
| | | | | | | | | | | | | 0.5 Poor Learner - Veh Good Learner | -0.0896 (0.322) | -0.278 | 1 |
| | | | | | | | | | | | | 0.5 Poor Learner - 0.05 Good Learner | -0.3838 (0.324) | -1.185 | 0.9364 |
| | | | | | | | | | | | | 0.5 Poor Learner - 0.5 Good Learner | -0.4158 (0.324) | -1.282 | 0.9057 |
| | | | | | | | | | | | | 0.5 Poor Learner - 1.5 Good Learner | -0.2818 (0.375) | -0.751 | 0.9954 |
| | | | | | | | | | | | | 1.5 Poor Learner - Veh Good Learner | 0.4933 (0.456) | 1.081 | 0.9609 |
| | | | | | | | | | | | | 1.5 Poor Learner - 0.05 Good Learner | 0.1992 (0.458) | 0.435 | 0.9999 |
| | | | | | | | | | | | | 1.5 Poor Learner - 0.5 Good Learner | 0.1672 (0.458) | 0.365 | 1 |
| | | | | | | | | | | | | 1.5 Poor Learner - 1.5 Good Learner | 0.3012 (0.495) | 0.608 | 0.9988 |
| | | | | | | | | | | | | Veh Good Learner - 0.05 Good Learner | -0.2941 (0.125) | -2.351 | 0.266 |
| | | | | | | | | | | | | Veh Good Learner - 0.5 Good Learner | -0.3261 (0.126) | -2.585 | 0.1614 |
| | | | | | | | | | | | | Veh Good Learner - 1.5 Good Learner | -0.1921 (0.226) | -0.851 | 0.9901 |
| | | | | | | | | | | | | 0.05 Good Learner - 0.5 Good Learner | -0.0320 (0.131) | -0.244 | 1 |
| | | | | | | | | | | | | 0.05 Good Learner - 1.5 Good Learner | 0.1020 (0.228) | 0.447 | 0.9998 |
| | | | | | | | | | | | | 0.5 Good Learner - 1.5 Good Learner | 0.1340 (0.228) | 0.587 | 0.999 |
| InverseRT | n=5 - Poor n=5 - Good | 159 | lmer | | (1\|id) | (Intercept) | -0.54368 (0.05033) | -10.803 | **1.18E-06** | | | | | | |
| *Baclofen* | | | | ang_sep90 | | ang_sep45 | 0.07102 (0.01597) | 4.447 | **1.77E-05** | **D** | Dose | Veh - 0.05 | 0.00917 (0.0128) | 0.717 | 0.89 |
| | | | | | | ang_sep30 | 0.09913 (0.01597) | 6.207 | **5.84E-09** | | | Veh - 0.5 | 0.01318 (0.0128) | 1.032 | 0.7313 |
| | | | | | | ang_sep15 | 0.11518 (0.01597) | 7.212 | **3.23E-11** | | | Veh - 1.5 | -0.09314 (0.0255) | -3.653 | **0.002** |
| | | | | | | ang_sep10 | 0.14000 (0.01613) | 8.679 | **9.58E-15** | | | 0.05 - 0.5 | 0.00401 (0.0128) | 0.314 | 0.9892 |
| | | | | Dose_Veh | | Dose_0.05 | -0.03762 (0.01807) | -2.082 | **0.039154** | | | 0.05 - 1.5 | -0.10231 (0.0255) | -4.013 | **0.0006** |
| | | | | | | Dose_0.5 | -0.06474 (0.01807) | -3.583 | **0.000469** | | | 0.5 - 1.5 | -0.10632 (0.0255) | -4.17 | **0.0003** |
| | | | | | | Dose_1.5 | 0.14096 (0.03745) | 3.764 | **0.000245** | | Dose\|LT | LT_Poor \| Veh - 0.05 | 0.03762 (0.0181) | 2.082 | 0.164 |
| | | | | LT_Poor | | LT_Good | -0.11393 (0.06972) | -1.634 | 0.137572 | | | LT_Poor \| Veh - 0.5 | 0.06474 (0.0181) | 3.583 | **0.0026** |
| | | | | Dose_Veh, LT_Poor | | Dose_0.05: LT_Good | 0.05691 (0.02555) | 2.227 | **0.027538** | | | LT_Poor \| Veh - 1.5 | -0.14096 (0.0375) | -3.76 | **0.0014** |
| | | | | | | Dose_0.5: LT_Good | 0.10312 (0.02555) | 4.035 | **8.96E-05** | | | LT_Poor \| 0.05 - 0.5 | 0.02711 (0.0181) | 1.501 | 0.4399 |
| | | | | | | Dose_1.5: LT_Good | -0.09563 (0.05094) | -1.878 | *0.062526* | | | LT_Poor \| 0.05 - 1.5 | -0.17858 (0.0375) | -4.764 | **<.0001** |
| | | | anova(lmer) | | | LT | | F(1,8.3) = 2.020 | 0.191569 | | | LT_Poor \| 0.5 - 1.5 | -0.20570 (0.0375) | -5.487 | **<.0001** |
| | | | | | | Dose | | F(3,139) = 6.049 | **6.68E-04** | | | LT_Good \| Veh - 0.05 | -0.01929 (0.0181) | -1.068 | 0.7098 |
| | | | | | | ang_sep | | F(4,139) = 22.422 | **2.65E-14** | | | LT_Good \| Veh - 0.5 | -0.03838 (0.0181) | -2.124 | 0.1506 |
| | | | | | | Dose:LT | | F(3,139) = 8.560 | **2.95E-05** | | | LT_Good \| Veh - 1.5 | -0.04533 (0.0346) | -1.311 | 0.5573 |
| | | | | | | | | | | | | LT_Good \| 0.05 - 0.5 | -0.01909 (0.0181) | -1.056 | 0.7165 |
| | | | | | | | | | | | | LT_Good \| 0.05 - 1.5 | -0.02604 (0.0346) | -0.753 | 0.8751 |
| | | | | | | | | | | | | LT_Good \| 0.5 - 1.5 | -0.00695 (0.0346) | -0.201 | 0.9971 |
| | | | | | | | | | | | Dose*LT | Veh Poor Learner - 0.05 Poor Learner | 0.03762 (0.0181) | 2.082 | 0.4315 |
| | | | | | | | | | | | | Veh Poor Learner - 0.5 Poor Learner | 0.06474 (0.0181) | 3.583 | **0.0108** |
| | | | | | | | | | | | | Veh Poor Learner - 1.5 Poor Learner | -0.14096 (0.0375) | -3.76 | **0.0059** |
| | | | | | | | | | | | | Veh Poor Learner - Veh Good Learner | 0.11393 (0.0697) | 1.634 | 0.7237 |
| | | | | | | | | | | | | Veh Poor Learner - 0.05 Good Learner | 0.09464 (0.0697) | 1.357 | 0.8551 |

| | | | | | | | | | | | | | | | |
|---|---|---|---|---|---|---|---|---|---|---|---|---|---|---|---|
| | | | | | | | | | | | | Veh Poor Learner - 0.5 Good Learner | 0.07555 (0.0697) | 1.084 | 0.945 |
| | | | | | | | | | | | | Veh Poor Learner - 1.5 Good Learner | 0.06861 (0.0757) | 0.906 | 0.9796 |
| | | | | | | | | | | | | 0.05 Poor Learner - 0.5 Poor Learner | 0.02711 (0.0181) | 1.501 | 0.8059 |
| | | | | | | | | | | | | 0.05 Poor Learner - 1.5 Poor Learner | -0.17858 (0.0375) | -4.764 | **0.0001** |
| | | | | | | | | | | | | 0.05 Poor Learner - Veh Good Learner | 0.07631 (0.0697) | 1.094 | 0.9424 |
| | | | | | | | | | | | | 0.05 Poor Learner - 0.05 Good Learner | 0.05702 (0.0697) | 0.818 | 0.9871 |
| | | | | | | | | | | | | 0.05 Poor Learner - 0.5 Good Learner | 0.03793 (0.0697) | 0.544 | 0.9989 |
| | | | | | | | | | | | | 0.05 Poor Learner - 1.5 Good Learner | 0.03098 (0.0757) | 0.409 | 0.9998 |
| | | | | | | | | | | | | 0.5 Poor Learner - 1.5 Poor Learner | -0.20570 (0.0375) | -5.487 | **<.0001** |
| | | | | | | | | | | | | 0.5 Poor Learner - Veh Good Learner | 0.04919 (0.0697) | 0.706 | 0.9944 |
| | | | | | | | | | | | | 0.5 Poor Learner - 0.05 Good Learner | 0.02990 (0.0697) | 0.429 | 0.9998 |
| | | | | | | | | | | | | 0.5 Poor Learner - 0.5 Good Learner | 0.01082 (0.0697) | 0.155 | 1 |
| | | | | | | | | | | | | 0.5 Poor Learner - 1.5 Good Learner | 0.00387 (0.0757) | 0.051 | 1 |
| | | | | | | | | | | | | 1.5 Poor Learner - Veh Good Learner | 0.25489 (0.0771) | 3.307 | *0.078* |
| | | | | | | | | | | | | 1.5 Poor Learner - 0.05 Good Learner | 0.23560 (0.0771) | 3.057 | 0.1177 |
| | | | | | | | | | | | | 1.5 Poor Learner - 0.5 Good Learner | 0.21651 (0.0771) | 2.809 | 0.174 |
| | | | | | | | | | | | | 1.5 Poor Learner - 1.5 Good Learner | 0.20957 (0.0825) | 2.54 | 0.2439 |
| | | | | | | | | | | | | Veh Good Learner - 0.05 Good Learner | -0.01929 (0.0181) | -1.068 | 0.9624 |
| | | | | | | | | | | | | Veh Good Learner - 0.5 Good Learner | -0.03838 (0.0181) | -2.124 | 0.4049 |
| | | | | | | | | | | | | Veh Good Learner - 1.5 Good Learner | -0.04533 (0.0346) | -1.311 | 0.8932 |
| | | | | | | | | | | | | 0.05 Good Learner - 0.5 Good Learner | -0.01909 (0.0181) | -1.056 | 0.9645 |
| | | | | | | | | | | | | 0.05 Good Learner - 1.5 Good Learner | -0.02604 (0.0346) | -0.753 | 0.9951 |
| | | | | | | | | | | | | 0.5 Good Learner - 1.5 Good Learner | -0.00695 (0.0346) | -0.201 | 1 |
| Accuracy | n=5 - Poor n=5 - Good | 4556 | glmer(logit) | | (1\|id) | (Intercept) | 0.31417 (0.19613) | 1.602 | 0.10919 | | | | | | |
| *Alogabat* | | | | ang_sep90 | | ang_sep45 | 0.08074 (0.11164) | 0.723 | 0.46951 | **E** | Dose | Veh - 3 | -0.3264 (0.0916) | -3.564 | **0.0021** |
| | | | | | | ang_sep30 | -0.29015 (0.10764) | -2.695 | **0.00703** | | | Veh - 10 | -0.5892 (0.1050) | -5.597 | **<.0001** |
| | | | | | | ang_sep15 | -0.66945 (0.10725) | -6.242 | **4.33E-10** | | | Veh - 30 | -0.2476 (0.1070) | *-2.312* | *0.0953* |
| | | | | | | ang_sep10 | -0.89572 (0.11508) | -7.784 | **7.05E-15** | | | 3 - 10 | -0.2628 (0.1110) | -2.376 | *0.0818* |
| | | | | Dose_Veh | | Dose_3 | 0.49763 (0.11415) | 4.359 | **1.30E-05** | | | 3 - 30 | 0.0789 (0.1120) | 0.702 | 0.8962 |
| | | | | | | Dose_10 | 1.13432 (0.16660) | 6.808 | **9.87E-12** | | | 10 - 30 | 0.3416 (0.1230) | 2.782 | **0.0277** |
| | | | | | | Dose_30 | -0.11599 (0.16392) | -0.708 | 0.47919 | | Dose\|LT | LT_Poor : Veh - 3 | -0.498 (0.114) | -4.359 | **0.0001** |
| | | | | LT_Poor | | LT_Good | 1.23780 (0.26429) | 4.683 | **2.82E-06** | | | LT_Poor : Veh - 10 | -1.134 (0.167) | -6.808 | **<.0001** |
| | | | | Dose_Veh, LT_Poor | | Dose_3: LT_Good | -0.34242 (0.18348) | -1.866 | *0.062* | | | LT_Poor : Veh - 30 | 0.116 (0.164) | 0.708 | 0.8941 |
| | | | | | | Dose_10: LT_Good | -1.09027 (0.21047) | -5.18 | **2.22E-07** | | | LT_Poor : 3 - 10 | -0.637 (0.167) | -3.808 | **0.0008** |
| | | | | | | Dose_30: LT_Good | 0.72712 (0.21477) | 3.386 | **0.00071** | | | LT_Poor : 3 - 30 | 0.614 (0.165) | 3.709 | **0.0012** |
| | | | | | | | | | | | | LT_Poor : 10 - 30 | 1.250 (0.204) | 6.141 | **<.0001** |
| | | | | | | | | | | | | LT_Good : Veh - 3 | -0.155 (0.143) | -1.082 | 0.7006 |
| | | | | | | | | | | | | LT_Good : Veh - 10 | -0.044 (0.129) | -0.342 | 0.9862 |

| | | | | | | | | | | | | | | | |
|---|---|---|---|---|---|---|---|---|---|---|---|---|---|---|---|
| | | | | | | | | | | | | LT_Good : Veh - 30 | -0.611 (0.138) | -4.418 | **0.0001** |
| | | | | | | | | | | | | LT_Good : 3 - 10 | 0.111 (0.145) | 0.767 | 0.8695 |
| | | | | | | | | | | | | LT_Good : 3 - 30 | -0.456 (0.153) | -2.977 | **0.0154** |
| | | | | | | | | | | | | LT_Good : 10 - 30 | -0.567 (0.138) | -4.112 | **0.0002** |
| | | | | | | | | | | | Dose*LT | Veh Poor Learner - 3 Poor Learner | -0.498 (0.114) | -4.359 | **0.0003** |
| | | | | | | | | | | | | Veh Poor Learner - 10 Poor Learner | -1.134 (0.167) | -6.808 | **<.0001** |
| | | | | | | | | | | | | Veh Poor Learner - 30 Poor Learner | 0.116 (0.164) | 0.708 | 0.9968 |
| | | | | | | | | | | | | Veh Poor Learner - Veh Good Learner | -1.238 (0.264) | -4.683 | **0.0001** |
| | | | | | | | | | | | | Veh Poor Learner - 3 Good Learner | -1.393 (0.273) | -5.107 | **<.0001** |
| | | | | | | | | | | | | Veh Poor Learner - 10 Good Learner | -1.282 (0.264) | -4.857 | **<.0001** |
| | | | | | | | | | | | | Veh Poor Learner - 30 Good Learner | -1.849 (0.269) | -6.869 | **<.0001** |
| | | | | | | | | | | | | 3 Poor Learner - 10 Poor Learner | -0.637 (0.167) | -3.808 | **0.0035** |
| | | | | | | | | | | | | 3 Poor Learner - 30 Poor Learner | 0.614 (0.165) | 3.709 | **0.0051** |
| | | | | | | | | | | | | 3 Poor Learner - Veh Good Learner | -0.740 (0.264) | -2.803 | *0.0941* |
| | | | | | | | | | | | | 3 Poor Learner - 3 Good Learner | -0.895 (0.273) | -3.284 | **0.0228** |
| | | | | | | | | | | | | 3 Poor Learner - 10 Good Learner | -0.784 (0.264) | -2.975 | *0.0588* |
| | | | | | | | | | | | | 3 Poor Learner - 30 Good Learner | -1.351 (0.269) | -5.027 | **<.0001** |
| | | | | | | | | | | | | 10 Poor Learner - 30 Poor Learner | 1.250 (0.204) | 6.141 | **<.0001** |
| | | | | | | | | | | | | 10 Poor Learner - Veh Good Learner | -0.103 (0.292) | -0.355 | 1 |
| | | | | | | | | | | | | 10 Poor Learner - 3 Good Learner | -0.259 (0.299) | -0.864 | 0.9891 |
| | | | | | | | | | | | | 10 Poor Learner - 10 Good Learner | -0.148 (0.291) | -0.507 | 0.9996 |
| | | | | | | | | | | | | 10 Poor Learner - 30 Good Learner | -0.715 (0.296) | -2.415 | 0.2337 |
| | | | | | | | | | | | | 30 Poor Learner - Veh Good Learner | -1.354 (0.291) | -4.656 | **0.0001** |
| | | | | | | | | | | | | 30 Poor Learner - 3 Good Learner | -1.509 (0.298) | -5.06 | **<.0001** |
| | | | | | | | | | | | | 30 Poor Learner - 10 Good Learner | -1.398 (0.290) | -4.812 | **<.0001** |
| | | | | | | | | | | | | 30 Poor Learner - 30 Good Learner | -1.965 (0.295) | -6.651 | **<.0001** |
| | | | | | | | | | | | | Veh Good Learner - 3 Good Learner | -0.155 (0.143) | -1.082 | 0.9607 |
| | | | | | | | | | | | | Veh Good Learner - 10 Good Learner | -0.044 (0.129) | -0.342 | 1 |
| | | | | | | | | | | | | Veh Good Learner - 30 Good Learner | -0.611 (0.138) | -4.418 | **0.0003** |
| | | | | | | | | | | | | 3 Good Learner - 10 Good Learner | 0.111 (0.145) | 0.767 | 0.9947 |
| | | | | | | | | | | | | 3 Good Learner - 30 Good Learner | -0.456 (0.153) | -2.977 | *0.0585* |
| | | | | | | | | | | | | 10 Good Learner - 30 Good Learner | -0.567 (0.138) | -4.112 | **0.001** |
| | InverseRT | n=5 - Poor n=5 - Good | 159 | lmer | | (1\|id) | (Intercept) | -0.4718 (0.0434) | -10.872 | **3.85E-07** | | | | | |
| | *Alogabat* | | | | ang_sep90 | | ang_sep45 | 0.07865 (0.01753) | 4.487 | **1.50E-05** | **F** | Dose | No significant main effect *(anova(lmer))* so pairwise not performed | | |
| | | | | | | | ang_sep30 | 0.0849 (0.01753) | 4.844 | **3.35E-06** | | Dose\|LT | LT_Poor : Veh - 3 | 0.000518 (0.0206) | 0.025 | 1 |
| | | | | | | | ang_sep15 | 0.1024 (0.01753) | 5.843 | **3.48E-08** | | | LT_Poor : Veh - 10 | -0.003863 (0.0290) | -0.133 | 0.9992 |

| | | | | | | | | | | |
|---|---|---|---|---|---|---|---|---|---|---|
| | | ang_sep10 | 0.1146 (0.01859) | 6.165 | **7.20E-09** | | LT_Poor : Veh - 30 | 0.001238 (0.0300) | 0.041 | 1 |
| | Dose_Veh | Dose_3 | -5.18e-04 (0.02063) | -0.025 | 0.98 | | LT_Poor : 3 - 10 | -0.004381 (0.0290) | -0.151 | 0.9988 |
| | | Dose_10 | 3.863e-03 (0.02898) | 0.133 | 0.8942 | | LT_Poor : 3 - 30 | 0.000720 (0.0300) | 0.024 | 1 |
| | | Dose_30 | -1.24e-03 (0.0299) | -0.041 | 0.967 | | LT_Poor : 10 - 30 | 0.005101 (0.0354) | 0.144 | 0.9989 |
| | LT_Poor | LT_Good | -0.06262 (0.0593) | -1.056 | 0.3175 | | LT_Good : Veh - 3 | -0.008759 (0.0225) | -0.389 | 0.9799 |
| | Dose_Veh, LT_Poor | Dose_3: LT_Good | 9.277e-03 (0.03057) | 0.303 | 0.762 | | LT_Good : Veh - 10 | 0.057542 (0.0204) | 2.826 | **0.0275** |
| | | Dose_10: LT_Good | -0.0614 (0.03541) | -1.734 | *0.0851* | | LT_Good : Veh - 30 | -0.040181 (0.0204) | -1.973 | 0.2032 |
| | | Dose_30: LT_Good | 0.04142 (0.03619) | 1.145 | 0.2543 | | LT_Good : 3 - 10 | 0.066301 (0.0225) | 2.95 | **0.0193** |
| anova(lmer) | | LT | | F(1,8.1) = 1.308 | 0.2856 | | LT_Good : 3 - 30 | -0.031422 (0.0225) | -1.398 | 0.5027 |
| | | Dose | | F(3,140.2) = 1.834 | 0.1438 | | LT_Good : 10 - 30 | -0.097723 (0.0201) | -4.853 | **<.0001** |
| | | ang_sep | | F(4,139) =2.4567 | **8.97E-09** | Dose*LT | Veh Poor Learner - 3 Poor Learner | 0.000518 (0.0206) | 0.025 | 1 |
| | | Dose:LT | | F(3,140.1) = 2.291 | *0.0809* | | Veh Poor Learner - 10 Poor Learner | -0.003863 (0.0290) | -0.133 | 1 |
| | | | | | | | Veh Poor Learner - 30 Poor Learner | 0.001238 (0.0300) | 0.041 | 1 |
| | | | | | | | Veh Poor Learner - Veh Good Learner | 0.062615 (0.0593) | 1.056 | 0.9524 |
| | | | | | | | Veh Poor Learner - 3 Good Learner | 0.053856 (0.0601) | 0.897 | 0.9796 |
| | | | | | | | Veh Poor Learner - 10 Good Learner | 0.120157 (0.0593) | 2.028 | 0.5117 |
| | | | | | | | Veh Poor Learner - 30 Good Learner | 0.022435 (0.0593) | 0.379 | 0.9999 |
| | | | | | | | 3 Poor Learner - 10 Poor Learner | -0.004381 (0.0290) | -0.151 | 1 |
| | | | | | | | 3 Poor Learner - 30 Poor Learner | 0.000720 (0.0300) | 0.024 | 1 |
| | | | | | | | 3 Poor Learner - Veh Good Learner | 0.062098 (0.0592) | 1.05 | 0.9536 |
| | | | | | | | 3 Poor Learner - 3 Good Learner | 0.053339 (0.0599) | 0.89 | 0.9803 |
| | | | | | | | 3 Poor Learner - 10 Good Learner | 0.119640 (0.0591) | 2.025 | 0.5136 |
| | | | | | | | 3 Poor Learner - 30 Good Learner | 0.021917 (0.0591) | 0.371 | 0.9999 |
| | | | | | | | 10 Poor Learner - 30 Poor Learner | 0.005101 (0.0354) | 0.144 | 1 |
| | | | | | | | 10 Poor Learner - Veh Good Learner | 0.066478 (0.0627) | 1.059 | 0.9539 |
| | | | | | | | 10 Poor Learner - 3 Good Learner | 0.057719 (0.0634) | 0.91 | 0.9793 |
| | | | | | | | 10 Poor Learner - 10 Good Learner | 0.124020 (0.0627) | 1.979 | 0.531 |
| | | | | | | | 10 Poor Learner - 30 Good Learner | 0.026297 (0.0627) | 0.42 | 0.9998 |
| | | | | | | | 30 Poor Learner - Veh Good Learner | 0.061377 (0.0632) | 0.971 | 0.9705 |
| | | | | | | | 30 Poor Learner - 3 Good Learner | 0.052618 (0.0639) | 0.824 | 0.9881 |
| | | | | | | | 30 Poor Learner - 10 Good Learner | 0.118919 (0.0631) | 1.884 | 0.5837 |

| | | | | | | | | | | | | | |
|---|---|---|---|---|---|---|---|---|---|---|---|---|---|
| | | | | | | | | | | 30 Poor Learner - 30 Good Learner | 0.021197 (0.0631) | 0.336 | 1 |
| | | | | | | | | | | Veh Good Learner - 3 Good Learner | -0.008759 (0.0225) | -0.389 | 0.9999 |
| | | | | | | | | | | Veh Good Learner - 10 Good Learner | 0.057542 (0.0204) | 2.826 | *0.0971* |
| | | | | | | | | | | Veh Good Learner - 30 Good Learner | -0.040181 (0.0204) | -1.973 | 0.5038 |
| | | | | | | | | | | 3 Good Learner - 10 Good Learner | 0.066301 (0.0225) | 2.95 | *0.0706* |
| | | | | | | | | | | 3 Good Learner - 30 Good Learner | -0.031422 (0.0225) | -1.398 | 0.8568 |
| | | | | | | | | | | 10 Good Learner - 30 Good Learner | -0.097723 (0.0201) | -4.853 | **0.0001** |
| **5** | Total DAPI Assay 1 Pvalb, Sst | n=6 - Poor n=5 - Good | anova(lmer) | (1\|id) | LT | F(1,6.01) = 0.0487 | 0.832698 | Figure S5 | Region significance means no comparisons were made between regions with specific marker co-expression | | | | |
| | | | | | Region | F(3,16.8) = 6.982 | **0.002963** | | | | | | |
| | | | | | LT:Region | F(3,16.8) =0.202 | 0.8935 | | | | | | |
| | Total DAPI Assay 2 slc17A7, VIP | n=6 - Poor n=5 - Good | anova(lmer) | (1\|id) | LT | F(1,7.9) = 0.0089 | 0.9274 | | | | | | |
| | | | | | Region | F(3,22.38) = 17.81 | **3.91E-06** | | | | | | |
| | | | | | LT:Region | F(3,22.38) = 0.1024 | 0.9578 | | | | | | |
| | sqrt GABRA5:Actin | n=6 - Poor n=5 - Good | anova(lmer) | | LT | F(1,7.9) = 0.015 | 0.9057 | **E** | LT | No significant main effect *(anova(lmer))* so pairwise not performed | | | |
| | | | | | Region | F(3,20.97) = 15.51 | 1.53E-05 | | Region | Taken from main effects, anova(lmer) | | | |
| | | | | | LT:Region | F(3,20.97) = 2.28 | *0.1084* | | LT\|Region | Cg1, Poor-Good | 0.0427 (0.176) | 0.243 | 0.81 |
| | | | | | | | | | | PrL, Poor-Good | -0.3813 (0.186) | -2.046 | *0.0512* |
| | | | | | | | | | | IL, Poor-Good | 0.0914 (0.186) | 0.49 | 0.6282 |
| | | | | | | | | | | V1, Poor-Good | 0.1897 (0.196) | 0.968 | 0.3417 |
| | Number of Pvalb - GABRA5+ cells | n=6 - Poor n=5 - Good | anova(lmer) | (1\|id) | LT | F(1,6.1) = 0.0103 | 0.9223 | **F** | LT | No significant main effect *(anova(lmer))* so pairwise not performed | | | |
| | | | | | Region | F(3,16.7) = 1.45 | 0.2633 | | Region | No significant main effect *(anova(lmer))* so pairwise not performed | | | |
| | | | | | LT:Region | F(3,16.7) = 2.12 | 0.1128 | | LT\|Region | No significant main effect *(anova(lmer))* so pairwise not performed | | | |
| | Number of Sst - GABRA5+ cells | n=6 - Poor n=5 - Good | anova(lmer) | (1\|id) | LT | F(1,22) = 11.45 | **0.002672** | | Region | Explained by total DAPI variation, no pairwise comparisons performed | | | |
| | | | | | Region | F(3,22) = 6.11 | **0.003481** | | LT\|Region | Cg1, Poor-Good | -18.40 (12.5) | -1.47 | 0.1558 |
| | | | | | LT:Region | F(3, 22) = 2.44 | *0.0912* | | | PrL, Poor-Good | -49.93 (12.5) | -3.989 | **0.0006** |
| | | | | | | | | | | IL, Poor-Good | -4.33 (12.5) | -0.346 | 0.7325 |
| | | | | | | | | | | V1, Poor-Good | -14.67 (14.3) | -1.028 | 0.3153 |
| | Number of VIP - GABRA5+ cells | n=6 - Poor n=5 - Good | anova(lmer) | (1\|id) | LT | F(1,29) = 0.7821 | 0.3838 | | Region | Explained by total DAPI variation, no pairwise comparisons performed | | | |
| | | | | | Region | F(3,29) = 9.737 | **0.000132** | | LT\|Region | No significant main effect *(anova(lmer))* so pairwise not performed | | | |
| | | | | | LT:Region | F(3, 29) = 1.077 | 0.3741 | | | | | | |
| | Number of slc17A7 - GABRA5+ cells | n=6 - Poor n=5 - Good | anova(lmer) | (1\|id) | LT | F(1,8.3) = 0.0257 | 0.8765 | | Region | Explained by total DAPI variation, no pairwise comparisons performed | | | |
| | | | | | Region | F(3,22.9) = 4.481 | **0.0129** | | LT\|Region | No significant main effect *(anova(lmer))* so pairwise not performed | | | |
| | | | | | LT:Region | F(3, 22.9) = 0.234 | 0.8717 | | | | | | |
| | Pvalb + GABRA5 mRNA copies | n=6 - Poor n=5 - Good | anova(lmer) | (1\|id) | LT | F(1,6.1) = 0.8375 | 0.3950 | **G** | Region | Explained by total DAPI variation, no pairwise comparisons performed | | | |
| | | | | | Region | F(3,16.3) = 6.510 | **0.0042** | | LT\|Region | Cg1, Poor-Good | -571 (825) | -0.693 | 0.5001 |
| | | | | | LT:Region | F(3, 16.3) = 7.925 | **0.0018** | | | PrL, Poor-Good | -2945 (825) | -3.571 | **0.0031** |
| | | | | | | | | | | IL, Poor-Good | -226 (825) | -0.274 | 0.7881 |
| | | | | | | | | | | V1, Poor-Good | 1380 (894) | 1.544 | 0.1419 |
| | Sst + GABRA5 mRNA copies | n=6 - Poor n=5 - Good | anova(lmer) | (1\|id) | LT | F(1,5.5) = 5.967 | *0.0536* | | LT | Poor - Good | Taken from main effects | | |
| | | | | | Region | F(3,15.9) = 10.46 | **0.0005** | | Region | Explained by total DAPI variation, no pairwise comparisons performed | | | |
| | | | | | LT:Region | F(3, 15.9) = 3.399 | **0.0438** | | LT\|Region | Cg1, Poor-Good | -437 (168) | -2.596 | **0.0187** |
| | | | | | | | | | | PrL, Poor-Good | -563 (168) | -3.343 | **0.0038** |
| | | | | | | | | | | IL, Poor-Good | 11 (168) | 0.065 | 0.9486 |
| | | | | | | | | | | V1, Poor-Good | -180 (186) | -0.967 | 0.3457 |
| | VIP + GABRA5 mRNA copies | n=6 - Poor n=5 - Good | anova(lmer) | (1\|id) | LT | F(1,29) = 3.261 | *0.08133* | | LT | Poor - Good | 68.9 (38.4) | 1.796 | 0.1117 |
| | | | | | Region | F(3,29) = 15.77 | **2.86E-06** | | Region | Explained by total DAPI variation, no pairwise comparisons performed | | | |
| | | | | | LT:Region | F(3, 29) = 2.005 | 0.1353 | | LT\|Region | No significant main effect *(anova(lmer))* so pairwise not performed | | | |

| | | | | | | | | | | |
|---|---|---|---|---|---|---|---|---|---|---|
| | slc17A7 + GABRA5 mRNA copies | n=6 - Poor<br>n=5 - Good | anova(lmer) | (1\|id) | LT | F(1,8.4) = 0.1321 | 0.7253 | | Region | Explained by total DAPI variation, no pairwise comparisons performed |
| | | | | | Region | F(3,23) = 9.269 | **0.0003** | | LT\|Region | No significant main effect *(anova(lmer))* so pairwise not performed |
| | | | | | LT:Region | F(3, 23) = 0.3225 | 0.8090 | | | |

| Figure | Outcome | Groups (n) | Obs No. | Test | Reference condition | Random Effect | Fixed Effect Condition | Estimate (SE) | Test Stat: z val, t val or F val | Pr(>\|z\|),Pr(>\|t\|) or Pr(>F) | Panel | post-hoc comparison | Comparison | Estimate (SE) | t/z ratio | Adjusted p value |
|---|---|---|---|---|---|---|---|---|---|---|---|---|---|---|---|---|
| **S2** | Sessions to Criteria | n=16 M | 33 | lmer | PT2 | (1\|id) | (Intercept) | 2.5625 (0.9499) | 2.698 | **0.00917** | | | | | | |
| | | | | | | | PT3 | 1.2409 (1.2870) | 0.964 | 0.3402 | | | | | | |
| | | | | | | | PT4 | 9.7480 (1.4136) | 6.896 | **<0.0001** | | | | | | |
| | | | | | VD1 | (1\|id) | (Intercept) | 7.801 (1.334) | 5.847 | 2.01E-07 | | | | | | |
| | | | | | | | VD2 | -4.090 (1.942) | -2.106 | **0.041** | | | | | | |
| | | | | | | | VD3 | -4.904 (2.057) | -2.384 | **0.021** | | | | | | |
| | | | | | | | VD4 | -4.504 (2.057) | -2.189 | **0.034** | | | | | | |
| | | | | | | | VD5 | -4.704 (2.057) | -1.8 | *0.078* | | | | | | |
| | Sessions to Criteria | n=16 F | 51 | lmer | PT2 | (1\|id) | (Intercept) | 11.562 (1.452) | 7.961 | **1.59E-09** | | | | | | |
| | | | | | | | | 1.125 (2.017) | 0.558 | 0.5834 | | | | | | |
| | | | | | | | | 7.268 (2.752) | 2.641 | **0.013** | | | | | | |
| | | | | | VD1 | (1\|id) | (Intercept) | 10.321 (4.102) | 2.516 | 0.016 | | | | | | |
| | | | | | | | VD2 | -4.000 (5.704) | 0.701 | 0.492 | | | | | | |
| | | | | | | | VD3 | -7.921 (5.963) | -1.128 | 0.2703 | | | | | | |
| | | | | | | | VD4 | | Not reached | | | | | | | |
| | | | | | | | VD5 | | Not reached | | | | | | | |
| **S7** | Accuracy | n=8 - Poor n=8 - Inter | 20990 | glmer(logit) | | (1\|id) | (Intercept) | 0.29676 (0.12486) | 2.377 | **0.017466** | | | | | | |
| | | | | | ang_sep90 | | ang_sep45 | 0.01149 (0.0632) | 0.182 | 0.855688 | **B** | LT\|ang_sep | ang_sep90, Poor-Inter | -0.68 (0.176) | -3.872 | **0.0001** |
| | | | | | | | ang_sep30 | -0.14343 (0.06295) | -2.278 | **0.022706** | | | ang_sep45, Poor-Inter | -0.656 (0.175) | -3.752 | **0.0002** |
| | | | | | | | ang_sep15 | -0.44843 (0.06442) | -6.961 | **3.38E-12** | | | ang_sep30, Poor-Inter | -0.588 (0.174) | -3.37 | **0.0008** |
| | | | | | | | ang_sep10 | -0.54129 (0.07161) | -7.558 | **4.08E-14** | | | ang_sep15, Poor-Inter | -0.389 (0.175) | -2.224 | **0.0261** |
| | | | | | LT_Poor | | LT_Inter | 0.76906 (0.1767) | 4.352 | **1.35E-05** | | | ang_sep10, Poor-Inter | -0.424 (0.18) | -2.356 | **0.0185** |
| | | | | | TDDclose | | TDDfar | -0.14879 (0.04365) | -3.408 | **0.000653** | | ang_sep | ang_sep90 - ang_sep45 | 0.000329 (0.0453) | 0.007 | 1 |
| | | | | | ang_sep90, LT_Poor | | ang_sep45:LT_Inter | -0.02365 (0.09062) | -0.261 | 0.794129 | | | ang_sep90 - ang_sep30 | 0.189527 (0.0448) | 4.234 | **0.0002** |
| | | | | | | | ang_sep30:LT_Inter | -0.09219 (0.08954) | -1.03 | 0.303163 | | | ang_sep90 - ang_sep15 | 0.593979 (0.0451) | 13.169 | **<.0001** |
| | | | | | | | ang_sep15:LT_Inter | -0.29109 (0.09021) | -3.227 | **0.001252** | | | ang_sep90 - ang_sep10 | 0.669362 (0.0497) | 13.462 | **<.0001** |
| | | | | | | | ang_sep10:LT_Inter | -0.25614 (0.09945) | -2.576 | **0.010007** | | | ang_sep45 - ang_sep30 | 0.189198 (0.0441) | 4.293 | **0.0002** |
| | | | | | TDDclose, LT_Poor | | LT_Inter:TDDfar | -0.1782 (0.0611) | -2.917 | **0.00354** | | | ang_sep45 - ang_sep15 | 0.593650 0.0444 | 13.371 | **<.0001** |
| | | | | | | | | | | | | | ang_sep45 - ang_sep10 | 0.669033 0.0490 | 13.643 | **<.0001** |
| | | | | | | | | | | | | | ang_sep30 - ang_sep15 | 0.404452 0.0438 | 9.226 | **<.0001** |
| | | | | | | | | | | | | | ang_sep30 - ang_sep10 | 0.479834 0.0485 | 9.885 | **<.0001** |
| | | | | | | | | | | | | | ang_sep15 - ang_sep10 | 0.075383 0.0488 | 1.546 | 0.5325 |
| | | | | | | | | | | | **C** | LT | Taken from main effects | | | |
| | | | | | | | | | | | **D** | LT\|TDD | TDDclose, Poor-Inter | -0.636 (0.167) | -3.813 | **0.0001** |
| | | | | | | | | | | | | | TDDfar, Poor-Inter | -0.458 (0.170) | -2.69 | **0.0072** |
| | | | | | | | | | | | | TDD\|LT | Poor, TDDclose - TDDfar | 0.149 (0.0437) | 3.408 | **0.0007** |
| | | | | | | | | | | | | | Inter, TDDclose - TDDfar | 0.327 (0.0427) | 7.649 | *<.0001* |
| | InverseRT | n=8 - Poor n=8 - Inter | 160 | lmer | | (1\|id) | (Intercept) | -0.62187 (0.039516) | -15.737 | **7.33E-11** | | | | | | |
| | | | | | ang_sep90 | | ang_sep45 | 0.049622 (0.011654) | 4.258 | **3.85E-05** | **E** | LT\|ang_sep | ang_sep90, Poor-Inter | -0.0458 (0.0556) | -0.824 | 0.4228 |
| | | | | | | | ang_sep30 | 0.05592 (0.011654) | 4.798 | **4.20E-06** | | | ang_sep45, Poor-Inter | -0.0516 (0.0556) | -0.927 | 0.3688 |
| | | | | | | | ang_sep15 | 0.042285 (0.011654) | 3.629 | **0.0004** | | | ang_sep30, Poor-Inter | -0.0668 (0.0556) | -1.2 | 0.2487 |
| | | | | | | | ang_sep10 | 0.042118 (0.011654) | 3.614 | **0.0004** | | | ang_sep15, Poor-Inter | -0.0878 (0.0556) | -1.578 | 0.1354 |
| | | | | | LT_Poor | | LT_Inter | 0.039649 (0.055884) | 0.709 | 0.489 | | | ang_sep10, Poor-Inter | -0.1079 (0.0556) | -1.94 | 0.0714 |
| | | | | | TDDclose | | TDDfar | 0.00607 (0.00737) | 0.824 | 0.412 | Not marked on figure | ang_sep | ang_sep90 - ang_sep45 | -0.05248 (0.00824) | -6.368 | <.0001 |
| | | | | | ang_sep90, LT_Poor | | ang_sep45:LT_Inter | 0.005709 (0.016481) | 0.346 | 0.730 | | | ang_sep90 - ang_sep30 | -0.06638 (0.00824) | -8.056 | <.0001 |
| | | | | | | | ang_sep30:LT_Inter | 0.020926 (0.016481) | 1.27 | 0.206 | | | ang_sep90 - ang_sep15 | -0.06326 (0.00824) | -7.677 | <.0001 |
| | | | | | | | ang_sep15:LT_Inter | 0.041946 (0.016481) | 2.545 | **0.012** | | | ang_sep90 - ang_sep10 | -0.07315 (0.00824) | -8.877 | <.0001 |
| | | | | | | | ang_sep10:LT_Inter | 0.062067 (0.016481) | 3.766 | **0.000** | | | ang_sep45 - ang_sep30 | -0.01391 (0.00824) | -1.688 | 0.4451 |
| | | | | | TDDclose, LT_Poor | | LT_Inter:TDDfar | 0.012401 (0.010423) | 1.19 | 0.236 | | | ang_sep45 - ang_sep15 | -0.01078 (0.00824) | -1.308 | 0.6865 |
| | | | | anova(lmer) | | | ang_sep | | F(4,134) = 25.6304 | **8.80E-16** | | | ang_sep45 - ang_sep10 | -0.02068 (0.00824) | -2.509 | 0.0947 |

| | | | | | | | | | | | | | | | | |
|---|---|---|---|---|---|---|---|---|---|---|---|---|---|---|---|---|
| | | | | | | | LT | | F(1,14) = 1.7344 | 0.209 | | | ang_sep30 - ang_sep15 | 0.00312 (0.00824) | 0.379 | 0.9956 |
| | | | | | | | TDD | | F(1, 134) = 5.5432 | **0.020** | | | ang_sep30 - ang_sep10 | -0.00677 (0.00824) | -0.821 | 0.9236 |
| | | | | | | | ang_sep:LT | | F(4, 134) = 4.9123 | 0.0010 | | | ang_sep15 - ang_sep10 | -0.00989 (0.00824) | -1.201 | 0.751 |
| | | | | | | | LT:TDD | | F(1, 134) = 1.4156 | 0.236 | **F** | LT | Taken from main effects | | | |
| | | | | | | | | | | | **G** | LT\|TDD | No significant main effect *(anova(lmer))* so pairwise not performed | | | |
| | | | | | | | | | | | | | No significant main effect *(anova(lmer))* so pairwise not performed | | | |
| **S8** | Accuracy | n=7 - Poor n=8 - Inter | 6445 | glmer(logit) | | (1\|id) | (Intercept) | 0.49393 (0.16355) | 3.02 | **0.00253** | | | | | | |
| | *Tiagabine* | | | | ang_sep90 | | ang_sep45 | 0.32515 (0.21607) | 0.21607 | 0.13236 | **A** | Dose | Veh - 0.5 | 0.1409 (0.073) | 1.931 | 0.215 |
| | | | | | | | ang_sep30 | -0.47112 (0.09871) | -4.773 | **1.82E-06** | | | Veh - 2.5 | 0.0656 (0.0771) | 0.851 | 0.8301 |
| | | | | | | | ang_sep15 | -0.295 (0.10014) | -2.946 | **0.00322** | | | Veh - 7.5 | 0.037 (0.0765) | 0.484 | 0.9626 |
| | | | | | | | ang_sep10 | -0.13725 (0.10705) | -1.282 | 0.1998 | | | 0.5 - 2.5 | -0.0754 (0.0771) | -0.978 | 0.762 |
| | | | | | Dose_Veh | | Dose_0.5 | -0.13482 (0.07993) | -1.687 | 0.09167 | | | 0.5 - 7.5 | -0.1039 (0.0801) | -1.297 | 0.5651 |
| | | | | | | | Dose_2.5 | -0.35917 (0.07929) | -4.53 | **5.91E-06** | | | 2.5 - 7.5 | -0.0285 (0.0816) | -0.35 | 0.9853 |
| | | | | | | | Dose_7.5 | -0.6641 (0.08155) | -8.143 | **3.85E-16** | | Dose\|LT | LT_Poor : Veh - 0.5 | 0.4711 (0.0987) | 4.773 | **<.0001** |
| | | | | | LT_Poor | | LT_Inter | -0.71609 (0.08929) | -8.02 | **1.06E-15** | | | LT_Poor : Veh - 2.5 | 0.295 (0.1) | 2.946 | **0.017** |
| | | | | | Dose_Veh, LT_Poor | | Dose_0.5: LT_Inter | 0.66037 (0.14599) | 0.14599 | **6.08E-06** | | | LT_Poor : Veh - 7.5 | 0.1373 (0.107) | 1.282 | 0.5743 |
| | | | | | | | Dose_2.5: LT_Inter | 0.45888 (0.15394) | 0.15394 | **0.00287** | | | LT_Poor : 0.5 - 2.5 | -0.1761 (0.104) | -1.699 | 0.3243 |
| | | | | | | | Dose_7.5: LT_Inter | 0.20047 (0.15267) | 0.15267 | 0.18917 | | | LT_Poor : 0.5 - 7.5 | -0.3339 (0.111) | -3.003 | **0.0142** |
| | | | | | | | | | | | | | LT_Poor : 2.5 - 7.5 | -0.1577 (0.107) | -1.478 | 0.4506 |
| | | | | | | | | | | | | | LT_Inter : Veh - 0.5 | -0.1893 (0.108) | -1.76 | 0.293 |
| | | | | | | | | | | | | | LT_Inter : Veh - 2.5 | -0.1639 (0.117) | -1.4 | 0.4992 |
| | | | | | | | | | | | | | LT_Inter : Veh - 7.5 | -0.0632 (0.109) | -0.58 | 0.9382 |
| | | | | | | | | | | | | | LT_Inter : 0.5 - 2.5 | 0.0254 (0.114) | 0.223 | 0.9961 |
| | | | | | | | | | | | | | LT_Inter : 0.5 - 7.5 | 0.126 (0.115) | 1.094 | 0.6932 |
| | | | | | | | | | | | | | LT_Inter : 2.5 - 7.5 | 0.1007 (0.123) | 0.815 | 0.8472 |
| | | | | | | | | | | | | Dose*LT | Veh Poor Learner - 0.5 Poor Learner | 0.4711 (0.0987) | 4.773 | **<.0001** |
| | | | | | | | | | | | | | Veh Poor Learner - 2.5 Poor Learner | 0.295 (0.1) | 2.946 | *0.0638* |
| | | | | | | | | | | | | | Veh Poor Learner - 7.5 Poor Learner | 0.1373 (0.107) | 1.282 | 0.9057 |
| | | | | | | | | | | | | | Veh Poor Learner - Veh Inter Learner | -0.3251 (0.216) | -1.505 | 0.8052 |
| | | | | | | | | | | | | | Veh Poor Learner - 0.5 Inter Learner | -0.5144 (0.217) | -2.372 | 0.2552 |
| | | | | | | | | | | | | | Veh Poor Learner - 2.5 Inter Learner | -0.489 (0.221) | -2.212 | 0.3443 |
| | | | | | | | | | | | | | Veh Poor Learner - 7.5 Inter Learner | -0.3884 (0.22) | -1.764 | 0.6448 |
| | | | | | | | | | | | | | 0.5 Poor Learner - 2.5 Poor Learner | -0.1761 (0.104) | -1.699 | 0.6883 |
| | | | | | | | | | | | | | 0.5 Poor Learner - 7.5 Poor Learner | -0.3339 (0.111) | -3.003 | *0.0543* |
| | | | | | | | | | | | | | 0.5 Poor Learner - Veh Inter Learner | -0.7963 (0.218) | -3.659 | **0.0062** |
| | | | | | | | | | | | | | 0.5 Poor Learner - 0.5 Inter Learner | -0.9855 (0.218) | -4.511 | **0.0002** |
| | | | | | | | | | | | | | 0.5 Poor Learner - 2.5 Inter Learner | -0.9601 (0.223) | -4.313 | **0.0004** |
| | | | | | | | | | | | | | 0.5 Poor Learner - 7.5 Inter Learner | -0.8595 (0.222) | -3.876 | **0.0027** |
| | | | | | | | | | | | | | 2.5 Poor Learner - 7.5 Poor Learner | -0.1577 (0.107) | -1.478 | 0.8192 |
| | | | | | | | | | | | | | 2.5 Poor Learner - Veh Inter Learner | -0.6202 (0.218) | -2.848 | *0.0836* |
| | | | | | | | | | | | | | 2.5 Poor Learner - 0.5 Inter Learner | -0.8094 (0.219) | -3.702 | **0.0053** |
| | | | | | | | | | | | | | 2.5 Poor Learner - 2.5 Inter Learner | -0.784 (0.223) | -3.521 | **0.0102** |
| | | | | | | | | | | | | | 2.5 Poor Learner - 7.5 Inter Learner | -0.6834 (0.222) | -3.081 | **0.0431** |

| | | | | | | | | | | | | | | | |
|---|---|---|---|---|---|---|---|---|---|---|---|---|---|---|---|
| | | | | | | | | | | | | 7.5 Poor Learner - Veh Inter Learner | -0.4624 (0.221) | -2.094 | 0.4186 |
| | | | | | | | | | | | | 7.5 Poor Learner - 0.5 Inter Learner | -0.6517 (0.222) | -2.94 | *0.0649* |
| | | | | | | | | | | | | 7.5 Poor Learner - 2.5 Inter Learner | -0.6263 (0.226) | -2.776 | 0.101 |
| | | | | | | | | | | | | 7.5 Poor Learner - 7.5 Inter Learner | -0.5256 (0.225) | -2.339 | 0.2725 |
| | | | | | | | | | | | | Veh Inter Learner - 0.5 Inter Learner | -0.1893 (0.108) | -1.76 | 0.6475 |
| | | | | | | | | | | | | Veh Inter Learner - 2.5 Inter Learner | -0.1639 (0.117) | -1.4 | 0.8576 |
| | | | | | | | | | | | | Veh Inter Learner - 7.5 Inter Learner | -0.0632 (0.109) | -0.58 | 0.9991 |
| | | | | | | | | | | | | 0.5 Inter Learner - 2.5 Inter Learner | 0.0254 (0.114) | 0.223 | 1 |
| | | | | | | | | | | | | 0.5 Inter Learner - 7.5 Inter Learner | 0.126 (0.115) | 1.094 | 0.9583 |
| | | | | | | | | | | | | 2.5 Inter Learner - 7.5 Inter Learner | 0.1007 (0.123) | 0.815 | 0.9923 |
| InverseRT | n=7 - Poor n=8 - Inter | 229 | lmer | | (1\|id) | (Intercept) | -0.637989 (0.044818) | -14.235 | **2.49E-10** | | | | | | |
| *Tiagabine* | | | | ang_sep90 | | ang_sep45 | 0.082821 (0.060471) | 1.37 | 0.191432 | **B** | Dose | Veh - 0.5 | -0.00959 (0.0124) | -0.773 | 0.8664 |
| | | | | | | ang_sep30 | 0.024689 (0.01734) | 1.424 | 0.156008 | | | Veh - 2.5 | 0.0000458 (0.0134) | 0.003 | 1 |
| | | | | | | ang_sep15 | -0.007223 (0.017665) | -0.409 | 0.683068 | | | Veh - 7.5 | -0.0497 (0.0134) | -3.721 | **0.0015** |
| | | | | | | ang_sep10 | 0.02198 (0.018733) | 1.173 | 0.242022 | | | 0.5 - 2.5 | 0.00964 (0.0132) | 0.727 | 0.8861 |
| | | | | Dose_Veh | | Dose_0.5 | 0.070744 (0.01388) | 5.097 | **7.85E-07** | | | 0.5 - 7.5 | -0.0401 (0.0139) | -2.885 | **0.0223** |
| | | | | | | Dose_2.5 | 0.062788 (0.01388) | 4.524 | **1.03E-05** | | | 2.5 - 7.5 | -0.0497 (0.0143) | -3.469 | **0.0035** |
| | | | | | | Dose_7.5 | 0.046621 (0.01388) | 3.359 | **0.000934** | | Dose\|LT | LT_Poor : Veh - 0.5 | -0.02469 (0.0173) | -1.424 | 0.486 |
| | | | | LT_Poor | | LT_Inter | 0.034271 (0.014952) | 2.292 | **0.022922** | | | LT_Poor : Veh - 2.5 | 0.00722 (0.0177) | 0.409 | 0.9769 |
| | | | | Dose_Veh, LT_Poor | | Dose_0.5: LT_Inter | -0.030197 (0.024797) | -1.218 | 0.224725 | | | LT_Poor : Veh - 7.5 | -0.02198 (0.0187) | -1.173 | 0.6446 |
| | | | | | | Dose_2.5: LT_Inter | 0.014353 (0.026751) | 0.537 | 0.59215 | | | LT_Poor : 0.5 - 2.5 | 0.03191 (0.0183) | 1.743 | 0.304 |
| | | | | | | Dose_7.5: LT_Inter | 0.055429 (0.026659) | 2.079 | **0.038842** | | | LT_Poor : 0.5 - 7.5 | 0.00271 (0.0195) | 0.139 | 0.999 |
| | | | anova(lmer) | | | LT | | F(1,13) = 2.4911 | 0.138401 | | | LT_Poor : 2.5 - 7.5 | -0.0292 (0.0189) | -1.547 | 0.4115 |
| | | | | | | Dose | | F(3,205) = 5.666 | **9.52E-04** | | | LT_Inter : Veh - 0.5 | 0.00551 (0.0177) | 0.31 | 0.9896 |
| | | | | | | ang_sep | | F(4,204) = 8.00 | **5.25E-06** | | | LT_Inter : Veh - 2.5 | -0.00713 (0.0201) | -0.354 | 0.9847 |
| | | | | | | Dose:LT | | F(3,205) = 3.286 | **2.18E-02** | | | LT_Inter : Veh - 7.5 | -0.07741 (0.019) | -4.074 | **0.0004** |
| | | | | | | | | | | | | LT_Inter : 0.5 - 2.5 | -0.01264 (0.0192) | -0.659 | 0.9121 |
| | | | | | | | | | | | | LT_Inter : 0.5 - 7.5 | -0.08292 (0.0196) | -4.186 | **0.0002** |
| | | | | | | | | | | | | LT_Inter : 2.5 - 7.5 | -0.07028 (0.0216) | -3.255 | **0.0072** |
| | | | | | | | | | | | Dose*LT | Veh Poor Learner - 0.5 Poor Learner | -0.02469 (0.0173) | -1.424 | 0.8454 |
| | | | | | | | | | | | | Veh Poor Learner - 2.5 Poor Learner | 0.00722 (0.0177) | 0.409 | 0.9999 |
| | | | | | | | | | | | | Veh Poor Learner - 7.5 Poor Learner | -0.02198 (0.0187) | -1.173 | 0.9388 |
| | | | | | | | | | | | | Veh Poor Learner - Veh Inter Learner | -0.08282 (0.0605) | -1.37 | 0.8569 |
| | | | | | | | | | | | | Veh Poor Learner - 0.5 Inter Learner | -0.07731 (0.0605) | -1.279 | 0.893 |
| | | | | | | | | | | | | Veh Poor Learner - 2.5 Inter Learner | -0.08995 (0.0611) | -1.473 | 0.8099 |
| | | | | | | | | | | | | Veh Poor Learner - 7.5 Inter Learner | -0.16023 (0.061) | -2.625 | 0.2199 |
| | | | | | | | | | | | | 0.5 Poor Learner - 2.5 Poor Learner | 0.03191 (0.0183) | 1.743 | 0.6588 |
| | | | | | | | | | | | | 0.5 Poor Learner - 7.5 Poor Learner | 0.00271 (0.0195) | 0.139 | 1 |
| | | | | | | | | | | | | 0.5 Poor Learner - Veh Inter Learner | -0.05813 (0.0607) | -0.958 | 0.9738 |
| | | | | | | | | | | | | 0.5 Poor Learner - 0.5 Inter Learner | -0.05262 (0.0607) | -0.868 | 0.9848 |
| | | | | | | | | | | | | 0.5 Poor Learner - 2.5 Inter Learner | -0.06526 (0.0613) | -1.065 | 0.955 |
| | | | | | | | | | | | | 0.5 Poor Learner - 7.5 Inter Learner | -0.13554 (0.0612) | -2.214 | 0.3951 |
| | | | | | | | | | | | | 2.5 Poor Learner - 7.5 Poor Learner | -0.0292 (0.0189) | -1.547 | 0.7809 |

| | | | | | | | | | | | | | | |
|---|---|---|---|---|---|---|---|---|---|---|---|---|---|---|
| | | | | | | | | | | | | 2.5 Poor Learner - Veh Inter Learner | -0.09004 (0.0607) | -1.483 | 0.8048 |
| | | | | | | | | | | | | 2.5 Poor Learner - 0.5 Inter Learner | -0.08454 (0.0607) | -1.392 | 0.8473 |
| | | | | | | | | | | | | 2.5 Poor Learner - 2.5 Inter Learner | -0.09717 (0.0613) | -1.585 | 0.7522 |
| | | | | | | | | | | | | 2.5 Poor Learner - 7.5 Inter Learner | -0.16745 (0.0613) | -2.732 | 0.1847 |
| | | | | | | | | | | | | 7.5 Poor Learner - Veh Inter Learner | -0.06084 (0.061) | -0.997 | 0.9679 |
| | | | | | | | | | | | | 7.5 Poor Learner - 0.5 Inter Learner | -0.05533 (0.061) | -0.907 | 0.9807 |
| | | | | | | | | | | | | 7.5 Poor Learner - 2.5 Inter Learner | -0.06797 (0.0616) | -1.103 | 0.9467 |
| | | | | | | | | | | | | 7.5 Poor Learner - 7.5 Inter Learner | -0.13825 (0.0616) | -2.245 | 0.3784 |
| | | | | | | | | | | | | Veh Inter Learner - 0.5 Inter Learner | 0.00551 (0.0177) | 0.31 | 1 |
| | | | | | | | | | | | | Veh Inter Learner - 2.5 Inter Learner | -0.00713 (0.0201) | -0.354 | 1 |
| | | | | | | | | | | | | Veh Inter Learner - 7.5 Inter Learner | -0.07741 (0.019) | -4.074 | **0.0017** |
| | | | | | | | | | | | | 0.5 Inter Learner - 2.5 Inter Learner | -0.01264 (0.0192) | -0.659 | 0.9979 |
| | | | | | | | | | | | | 0.5 Inter Learner - 7.5 Inter Learner | -0.08292 (0.0198) | -4.186 | **0.0011** |
| | | | | | | | | | | | | 2.5 Inter Learner - 7.5 Inter Learner | -0.07028 (0.0216) | -3.255 | **0.0284** |
| Accuracy | n=7 - Poor n=8 - Inter | 6410 | glmer(logit) | | (1\|id) | (Intercept) | 0.642085 (0.175076) | 3.667 | **0.000245** | | | | | | |
| Baclofen | | | | ang_sep90 | | ang_sep45 | 0.803184 (0.226622) | 3.544 | **0.000394** | **C** | Dose | Veh - 0.05 | 0.212 (0.074) | 2.866 | **0.0217** |
| | | | | | | ang_sep30 | -0.07505 (0.113264) | -0.663 | 0.507579 | | | Veh - 0.5 | 0.1739 (0.0725) | 2.399 | 0.0772 |
| | | | | | | ang_sep15 | -0.129297 (0.109113) | -1.185 | 0.236027 | | | Veh - 1.5 | -0.16 (0.104) | -1.537 | 0.4153 |
| | | | | | | ang_sep10 | 0.121429 (0.128092) | 0.948 | 0.34314 | | | 0.05 - 0.5 | -0.0381 (0.0738) | -0.516 | 0.9553 |
| | | | | Dose_Veh | | Dose_0.05 | 0.007741 (0.086965) | 0.089 | 0.929067 | | | 0.05 - 1.5 | -0.372 (0.104) | -3.565 | **0.0021** |
| | | | | | | Dose_0.5 | -0.324132 (0.084618) | -3.831 | **0.000128** | | | 0.5 - 1.5 | -0.3339 (0.103) | -3.244 | **0.0065** |
| | | | | | | Dose_1.5 | -0.819509 (0.084999) | -9.641 | **<.0001** | | Dose\|LT | LT_Poor : Veh - 0.05 | 0.0751 (0.113) | 0.663 | 0.9111 |
| | | | | LT_Poor | | LT_Inter | -1.035106 (0.091423) | -11.322 | **<.0001** | | | LT_Poor : Veh - 0.5 | 0.1293 (0.109) | 1.185 | 0.6364 |
| | | | | Dose_Veh, LT_Poor | | Dose_0.05: LT_Inter | -0.273911 (0.148037) | -1.85 | *0.06427* | | | LT_Poor : Veh - 1.5 | -0.1214 (0.128) | -0.948 | 0.7789 |
| | | | | | | Dose_0.5: LT_Inter | -0.089238 (0.144946) | -0.616 | 0.538115 | | | LT_Poor : 0.05 - 0.5 | 0.0542 (0.114) | 0.475 | 0.9647 |
| | | | | | | Dose_1.5: LT_Inter | 0.077172 (0.208036) | 0.371 | 0.710672 | | | LT_Poor : 0.05 - 1.5 | -0.1965 (0.13) | -1.515 | 0.4287 |
| | | | | | | | | | | | | LT_Poor : 0.5 - 1.5 | -0.2507 (0.126) | -1.997 | 0.189 |
| | | | | | | | | | | | | LT_Inter : Veh - 0.05 | 0.349 (0.0953) | 3.663 | **0.0014** |
| | | | | | | | | | | | | LT_Inter : Veh - 0.5 | 0.2185 (0.0954) | 2.29 | 0.1003 |
| | | | | | | | | | | | | LT_Inter : Veh - 1.5 | -0.1986 (0.164) | -1.211 | 0.62 |
| | | | | | | | | | | | | 0.05 - 0.5 | -0.1304 (0.0935) | -1.395 | 0.5026 |
| | | | | | | | | | | | | LT_Inter : 0.05 - 1.5 | -0.5476 (0.164) | -3.348 | **0.0045** |
| | | | | | | | | | | | | LT_Inter : 0.5 - 1.5 | -0.4171 (0.163) | -2.558 | *0.0515* |
| | | | | | | | | | | | Dose*LT | Veh Poor Learner - 0.05 Poor Learner | .0751 (0.11 | 0.663 | 0.9979 |
| | | | | | | | | | | | | Veh Poor Learner - 0.5 Poor Learner | .1293 (0.10 | 1.185 | 0.9364 |
| | | | | | | | | | | | | Veh Poor Learner - 1.5 Poor Learner | .1214 (0.12 | -0.948 | 0.9813 |
| | | | | | | | | | | | | Veh Poor Learner - Veh Inter Learner | .8032 (0.22 | -3.544 | **0.0094** |
| | | | | | | | | | | | | Veh Poor Learner - 0.05 Inter Learner | .4542 (0.22 | -2.012 | 0.4733 |
| | | | | | | | | | | | | Veh Poor Learner - 0.5 Inter Learner | .5846 (0.22 | -2.588 | 0.1601 |
| | | | | | | | | | | | | Veh Poor Learner - 1.5 Inter Learner | .0018 (0.26 | -3.814 | 0.0034 |
| | | | | | | | | | | | | 0.05 Poor Learner - 0.5 Poor Learner | .0542 (0.11 | 0.475 | 0.9998 |

| | | | | | | | | | | | | | | | |
|---|---|---|---|---|---|---|---|---|---|---|---|---|---|---|---|
| | | | | | | | | | | | | 0.05 Poor Learner - 1.5 Poor Learner | .1965 (0.13 | -1.515 | 0.7999 |
| | | | | | | | | | | | | 0.05 Poor Learner - Veh Inter Learner | .8782 (0.22 | -3.842 | 0.0031 |
| | | | | | | | | | | | | 0.05 Poor Learner - 0.05 Inter Learner | .5293 (0.22 | -2.324 | 0.28 |
| | | | | | | | | | | | | 0.05 Poor Learner - 0.5 Inter Learner | .6597 (0.22 | -2.895 | 0.0735 |
| | | | | | | | | | | | | 0.05 Poor Learner - 1.5 Inter Learner | .0768 (0.26 | -4.074 | 0.0012 |
| | | | | | | | | | | | | 0.5 Poor Learner - 1.5 Poor Learner | .2507 (0.12 | -1.997 | 0.4837 |
| | | | | | | | | | | | | 0.5 Poor Learner - Veh Inter Learner | .9325 (0.22 | -4.115 | 0.001 |
| | | | | | | | | | | | | 0.5 Poor Learner - 0.05 Inter Learner | .5835 (0.22 | -2.586 | 0.161 |
| | | | | | | | | | | | | 0.5 Poor Learner - 0.5 Inter Learner | .7139 (0.22 | -3.161 | **0.0338** |
| | | | | | | | | | | | | 0.5 Poor Learner - 1.5 Inter Learner | .1311 (0.26 | -4.306 | 0.0004 |
| | | | | | | | | | | | | 1.5 Poor Learner - Veh Inter Learner | .6818 (0.23 | -2.884 | 0.0758 |
| | | | | | | | | | | | | 1.5 Poor Learner - 0.05 Inter Learner | .3328 (0.23 | -1.413 | 0.8515 |
| | | | | | | | | | | | | 1.5 Poor Learner - 0.5 Inter Learner | .4632 (0.23 | -1.966 | 0.5053 |
| | | | | | | | | | | | | 1.5 Poor Learner - 1.5 Inter Learner | .8804 (0.27 | -3.247 | **0.0257** |
| | | | | | | | | | | | | Veh Inter Learner - 0.05 Inter Learner | .349 (0.095 | 3.663 | **0.0061** |
| | | | | | | | | | | | | Veh Inter Learner - 0.5 Inter Learner | 2185 (0.095 | 2.29 | 0.2989 |
| | | | | | | | | | | | | Veh Inter Learner - 1.5 Inter Learner | .1986 (0.16 | -1.211 | 0.9291 |
| | | | | | | | | | | | | 0.05 Inter Learner - 0.5 Inter Learner | 1304 (0.09 | -1.395 | 0.8601 |
| | | | | | | | | | | | | 0.05 Inter Learner - 1.5 Inter Learner | .5476 (0.16 | -3.348 | **0.0185** |
| | | | | | | | | | | | | 0.5 Inter Learner - 1.5 Inter Learner | .4171 (0.16 | -2.558 | 0.1715 |
| InverseRT | n=7 - Poor n=8 - Inter | 228 | lmer | | (1\|id) | (Intercept) | 196.783 (24.259) | 24.506 | **2.11E-08** | | | | | | |
| *Baclofen* | | | | ang_sep90 | | ang_sep45 | 5.61 (30.999) | 0.181 | 0.85832 | **D** | Dose | Veh - 0.05 | -0.0551 (0.0158) | -3.48 | **0.0034** |
| | | | | | | ang_sep30 | 46.698 (7.675) | 207.596 | **0.00887** | | | Veh - 0.5 | -0.0278 (0.0153) | -1.813 | 0.27 |
| | | | | | | ang_sep15 | -7.652 (16.869) | 206.264 | 0.65059 | | | Veh - 1.5 | -0.161 (0.0219) | -7.357 | **<.0001** |
| | | | | | | ang_sep10 | 173.137 (19.745) | 208.677 | **6.41E-16** | | | 0.05 - 0.5 | 0.0273 (0.016) | 1.708 | 0.3222 |
| | | | | Dose_Veh | | Dose_0.05 | -14.719 (12.72) | 203.009 | 0.24858 | | | 0.05 - 1.5 | -0.1059 (0.0221) | -4.798 | **<.0001** |
| | | | | | | Dose_0.5 | 7.473 (12.72) | 203.009 | 0.55756 | | | 0.5 - 1.5 | -0.1332 (0.0216) | -6.159 | **<.0001** |
| | | | | | | Dose_1.5 | 7.309 (12.72) | 203.009 | 0.56621 | | Dose\|LT | LT_Poor \| Veh - 0.05 | -0.0945 (0.0249) | -3.797 | **0.0011** |
| | | | | LT_Poor | | LT_Inter | 4.34 (13.325) | 203.25 | 0.74497 | | | LT_Poor \| Veh - 0.5 | -0.0124 (0.0237) | -0.521 | 0.954 |
| | | | | Dose_Veh, LT_Poor | | Dose_0.05: LT_Inter | -28.428 (22.531) | -1.262 | 0.20848 | | | LT_Poor \| Veh - 1.5 | -0.2138 (0.0278) | -7.677 | **<.0001** |
| | | | | | | Dose_0.5: LT_Inter | 36.239 (21.85) | 1.659 | *0.09873* | | | LT_Poor \| 0.05 - 0.5 | 0.0822 (0.0254) | 3.238 | **0.0076** |
| | | | | | | Dose_1.5: LT_Inter | -100.383 (31.06) | -3.232 | **0.00143** | | | LT_Poor \| 0.05 - 1.5 | -0.1192 (0.0284) | -4.195 | **0.0002** |
| | | | anova(lmer) | | | LT | | F(1,13.5) = 0.3766 | 0.5496 | | | LT_Poor \| 0.5 - 1.5 | -0.2014 (0.0273) | -7.381 | **<.0001** |
| | | | | | | Dose | | F(3,207) = 22.75 | **9.08E-13** | | | LT_Inter \| Veh - 0.05 | -0.0158 (0.0196) | -0.804 | 0.8525 |
| | | | | | | ang_sep | | F(4,203) = 1.05 | **0.3840** | | | LT_Inter \| Veh - 0.5 | -0.0433 (0.0195) | -2.223 | 0.1204 |
| | | | | | | Dose:LT | | F(3,207) = 7.34 | **0.0001** | | | LT_Inter \| Veh - 1.5 | -0.1083 (0.0338) | -3.204 | **0.0085** |
| | | | | | | | | | | | | LT_Inter \| 0.05 - 0.5 | -0.0275 (0.0195) | -1.414 | 0.492 |
| | | | | | | | | | | | | LT_Inter \| 0.05 - 1.5 | -0.0925 (0.0338) | -2.738 | **0.0337** |
| | | | | | | | | | | | | LT_Inter \| 0.5 - 1.5 | -0.065 (0.0336) | -1.936 | 0.2162 |
| | | | | | | | | | | | Dose*LT | Veh Poor Learner - 0.05 Poor Learner | -0.09453 (0.0249) | -3.797 | **0.0047** |
| | | | | | | | | | | | | Veh Poor Learner - 0.5 Poor Learner | -0.01236 (0.0237) | -0.521 | 0.9995 |

| | | | | | | | | | | | | | | | |
|---|---|---|---|---|---|---|---|---|---|---|---|---|---|---|---|
| | | | | | | | | | | | | Veh Poor Learner - 1.5 Poor Learner | -0.21377 (0.0278) | -7.677 | **<.0001** |
| | | | | | | | | | | | | Veh Poor Learner - Veh Inter Learner | -0.08269 (0.0587) | -1.41 | 0.8403 |
| | | | | | | | | | | | | Veh Poor Learner - 0.05 Inter Learner | -0.09845 (0.0587) | -1.678 | 0.7001 |
| | | | | | | | | | | | | Veh Poor Learner - 0.5 Inter Learner | -0.12599 (0.0586) | -2.149 | 0.4277 |
| | | | | | | | | | | | | Veh Poor Learner - 1.5 Inter Learner | -0.19099 (0.0647) | -2.951 | 0.1075 |
| | | | | | | | | | | | | 0.05 Poor Learner - 0.5 Poor Learner | 0.08217 (0.0254) | 3.238 | **0.0299** |
| | | | | | | | | | | | | 0.05 Poor Learner - 1.5 Poor Learner | -0.11923 (0.0284) | -4.195 | **0.001** |
| | | | | | | | | | | | | 0.05 Poor Learner - Veh Inter Learner | 0.01185 (0.0592) | 0.2 | 1 |
| | | | | | | | | | | | | 0.05 Poor Learner - 0.05 Inter Learner | -0.00391 (0.0592) | -0.066 | 1 |
| | | | | | | | | | | | | 0.05 Poor Learner - 0.5 Inter Learner | -0.03146 (0.0592) | -0.531 | 0.9993 |
| | | | | | | | | | | | | 0.05 Poor Learner - 1.5 Inter Learner | -0.09645 (0.0652) | -1.478 | 0.8109 |
| | | | | | | | | | | | | 0.5 Poor Learner - 1.5 Poor Learner | -0.2014 (0.0273) | -7.381 | **<.0001** |
| | | | | | | | | | | | | 0.5 Poor Learner - Veh Inter Learner | -0.07032 (0.0588) | -1.196 | 0.9216 |
| | | | | | | | | | | | | 0.5 Poor Learner - 0.05 Inter Learner | -0.08609 (0.0588) | -1.465 | 0.8145 |
| | | | | | | | | | | | | 0.5 Poor Learner - 0.5 Inter Learner | -0.11363 (0.0587) | -1.934 | 0.5491 |
| | | | | | | | | | | | | 0.5 Poor Learner - 1.5 Inter Learner | -0.17863 (0.0648) | -2.755 | 0.1557 |
| | | | | | | | | | | | | 1.5 Poor Learner - Veh Inter Learner | 0.13108 (0.0605) | 2.168 | 0.4132 |
| | | | | | | | | | | | | 1.5 Poor Learner - 0.05 Inter Learner | 0.11532 (0.0605) | 1.908 | 0.5632 |
| | | | | | | | | | | | | 1.5 Poor Learner - 0.5 Inter Learner | 0.08778 (0.0604) | 1.453 | 0.8212 |
| | | | | | | | | | | | | 1.5 Poor Learner - 1.5 Inter Learner | 0.02278 (0.0664) | 0.343 | 1 |
| | | | | | | | | | | | | Veh Inter Learner - 0.05 Inter Learner | -0.01576 (0.0196) | -0.804 | 0.9928 |
| | | | | | | | | | | | | Veh Inter Learner - 0.5 Inter Learner | -0.0433 (0.0195) | -2.223 | 0.3423 |
| | | | | | | | | | | | | Veh Inter Learner - 1.5 Inter Learner | -0.1083 (0.0338) | -3.204 | **0.0331** |
| | | | | | | | | | | | | 0.05 Inter Learner - 0.5 Inter Learner | -0.02754 (0.0195) | -1.414 | 0.8499 |
| | | | | | | | | | | | | 0.05 Inter Learner - 1.5 Inter Learner | -0.09254 (0.0338) | -2.738 | 0.1173 |
| | | | | | | | | | | | | 0.5 Inter Learner - 1.5 Inter Learner | -0.065 (0.0336) | -1.936 | 0.5278 |
| Accuracy | n=7 - Poor n=8 - Inter | 4254 | glmer(logit) | | (1\|id) | (Intercept) | 0.46684 (0.16794) | 2.78 | **0.00544** | | | | | | |
| *Alogabat* | | | | ang_sep90 | | ang_sep45 | 0.88065 (0.21375) | 4.12 | **3.79E-05** | **E** | Dose | Veh - 3 | -0.1186 (0.0921) | -1.288 | 0.5704 |
| | | | | | | ang_sep30 | 0.08592 (0.11399) | 0.754 | 0.451017 | | | Veh - 10 | -0.1332 (0.107) | -1.245 | 0.5981 |
| | | | | | | ang_sep15 | 0.30547 (0.15918) | 1.919 | *0.05499* | | | Veh - 30 | -0.3317 (0.105) | -3.171 | **0.0083** |
| | | | | | | ang_sep10 | 0.57691 (0.1641) | 3.516 | **0.000439** | | | 3 - 10 | -0.0145 (0.116) | -0.125 | 0.9993 |
| | | | | Dose_Veh | | Dose_3 | 0.01136 (0.11026) | 0.103 | 0.917941 | | | 3 - 30 | -0.213 (0.113) | -1.88 | 0.2367 |
| | | | | | | Dose_10 | -0.13505 (0.10858) | -1.244 | 0.213567 | | | 10 - 30 | -0.1985 (0.128) | -1.546 | 0.4098 |
| | | | | | | Dose_30 | -0.77224 (0.10641) | -7.257 | **3.96E-13** | | Dose\|LT | LT_Poor : Veh - 3 | -0.0859 (0.114) | -0.754 | 0.8751 |
| | | | | LT_Poor | | LT_Inter | -0.93778 (0.1126) | -8.328 | **<0.0001** | | | LT_Poor : Veh - 10 | -0.3055 (0.159) | -1.919 | 0.22 |
| | | | | Dose_Veh, LT_Poor | | Dose_3: LT_Inter | 0.06545 (0.18425) | 0.355 | 0.722398 | | | LT_Poor : Veh - 30 | -0.5769 (0.164) | -3.516 | **0.0025** |
| | | | | | | Dose_10: LT_Inter | -0.34462 (0.21514) | -1.602 | 0.109183 | | | LT_Poor : 3 - 10 | -0.2195 (0.159) | -1.377 | 0.5139 |
| | | | | | | Dose_30: LT_Inter | -0.49051 (0.20926) | -2.344 | **0.019076** | | | LT_Poor : 3 - 30 | -0.491 (0.167) | -2.931 | **0.0178** |

| | | | | |
|---|---|---|---|---|
| | LT_Poor : 10 - 30 | -0.2714 (0.213) | -1.272 | 0.581 |
| | LT_Inter : Veh - 3 | -0.1514 (0.145) | -1.046 | 0.7223 |
| | LT_Inter : Veh - 10 | 0.0392 (0.144) | 0.272 | 0.9929 |
| | LT_Inter : Veh - 30 | -0.0864 (0.13) | -0.666 | 0.91 |
| | LT_Inter : 3 - 10 | 0.1905 (0.17) | 1.119 | 0.6775 |
| | LT_Inter : 3 - 30 | 0.065 (0.152) | 0.427 | 0.9738 |
| | LT_Inter : 10 - 30 | -0.1256 (0.146) | -0.861 | 0.8249 |
| Dose*LT | Veh Poor Learner - 3 Poor Learner | -0.0859 (0.114) | -0.754 | 0.9953 |
| | Veh Poor Learner - 10 Poor Learner | -0.3055 (0.159) | -1.919 | 0.5377 |
| | Veh Poor Learner - 30 Poor Learner | -0.5769 (0.164) | -3.516 | **0.0104** |
| | Veh Poor Learner - Veh Inter Learner | -0.8806 (0.214) | -4.12 | **0.001** |
| | Veh Poor Learner - 3 Inter Learner | -1.032 (0.228) | -4.524 | **0.0002** |
| | Veh Poor Learner - 10 Inter Learner | -0.8415 (0.219) | -3.837 | **0.0031** |
| | Veh Poor Learner - 30 Inter Learner | -0.967 (0.215) | -4.489 | **0.0002** |
| | 3 Poor Learner - 10 Poor Learner | -0.2195 (0.159) | -1.377 | 0.868 |
| | 3 Poor Learner - 30 Poor Learner | -0.491 (0.167) | -2.931 | *0.0665* |
| | 3 Poor Learner - Veh Inter Learner | -0.7947 (0.215) | -3.695 | **0.0054** |
| | 3 Poor Learner - 3 Inter Learner | -0.9461(0.229) | -4.125 | **0.001** |
| | 3 Poor Learner - 10 Inter Learner | -0.7556 (0.221) | -3.423 | **0.0144** |
| | 3 Poor Learner - 30 Inter Learner | -0.8811 (0.217) | -4.067 | **0.0012** |
| | 10 Poor Learner - 30 Poor Learner | -0.2714 (0.213) | -1.272 | 0.9094 |
| | 10 Poor Learner - Veh Inter Learner | -0.5752 (0.242) | -2.377 | 0.2528 |
| | 10 Poor Learner - 3 Inter Learner | -0.7266 (0.254) | -2.855 | *0.082* |
| | 10 Poor Learner - 10 Inter Learner | -0.536 (0.247) | -2.166 | 0.3725 |
| | 10 Poor Learner - 30 Inter Learner | -0.6616 (0.244) | -2.717 | 0.1175 |
| | 30 Poor Learner - Veh Inter Learner | -0.3037 (0.246) | -1.234 | 0.922 |
| | 30 Poor Learner - 3 Inter Learner | -0.4551 (0.259) | -1.756 | 0.65 |
| | 30 Poor Learner - 10 Inter Learner | -0.2646 (0.25) | -1.059 | 0.9649 |
| | 30 Poor Learner - 30 Inter Learner | -0.3901 (0.248) | -1.575 | 0.7652 |
| | Veh Inter Learner - 3 Inter Learner | -0.1514 (0.145) | -1.046 | 0.9673 |
| | Veh Inter Learner - 10 Inter Learner | 0.0392 (0.144) | 0.272 | 1 |
| | Veh Inter Learner - 30 Inter Learner | -0.0864 (0.13) | -0.666 | 0.9978 |
| | 3 Inter Learner - 10 Inter Learner | 0.1905 (0.17) | 1.119 | 0.9528 |
| | 3 Inter Learner - 30 Inter Learner | 0.065 (0.152) | 0.427 | 0.9999 |
| | 10 Inter Learner - 30 Inter Learner | -0.1256 (0.146) | -0.861 | 0.9893 |

| | | | | | | | | | | | | | | | |
|---|---|---|---|---|---|---|---|---|---|---|---|---|---|---|---|
| InverseRT | n=7 - Poor n=8 - Inter | 151 | lmer | | (1\|id) | (Intercept) | -0.63529 (0.06782) | -9.368 | **3.22E-06** | | | | | | |
| *Alogabat* | | | | ang_sep90 | | ang_sep45 | 0.01281 (0.09096) | 0.141 | 0.890925 | **F** | Dose | Veh - 3 | 0.000935 (0.0146) | 0.064 | 0.9999 |
| | | | | | | ang_sep30 | -0.04044 (0.01883) | -2.148 | **0.033602** | | | Veh - 10 | 0.04702 (0.0172) | 2.738 | **0.0351** |
| | | | | | | ang_sep15 | -0.05829 (0.02692) | -2.165 | **0.032178** | | | Veh - 30 | -0.101748 (0.0167) | -6.105 | **<.0001** |
| | | | | | | ang_sep10 | 0.09372 (0.02692) | 3.481 | **0.000678** | | | 3 - 10 | 0.046085 (0.0187) | 2.461 | *0.0709* |
| | | | | Dose_Veh | | Dose_3 | 0.05612 (0.01657) | 3.387 | **0.000936** | | | 3 - 30 | -0.102683 (0.018) | -5.713 | **<.0001** |
| | | | | | | Dose_10 | 0.08404 (0.01657) | 5.072 | **1.33E-06** | | | 10 - 30 | -0.148768 (0.0208) | -7.151 | **<.0001** |
| | | | | | | Dose_30 | 0.07798 (0.01657) | 4.706 | **6.38E-06** | | Dose\|LT | LT_Poor : Veh - 3 | 0.0404 (0.0188) | 2.148 | 0.1437 |
| | | | | LT_Poor | | LT_Inter | 0.0821 (0.01729) | 4.749 | **5.32E-06** | | | LT_Poor : Veh - 10 | 0.0583 (0.0269) | 2.164 | 0.139 |
| | | | | Dose_Veh, LT_Poor | | Dose_3: LT_Inter | 0.07902 (0.02926) | 2.701 | **0.007843** | | | LT_Poor : Veh - 30 | -0.0937 (0.0269) | -3.479 | **0.0038** |
| | | | | | | Dose_10: LT_Inter | 0.02253 (0.03432) | 0.657 | 0.512639 | | | LT_Poor : 3 - 10 | 0.0178 (0.0269) | 0.662 | 0.911 |
| | | | | | | Dose_30: LT_Inter | 0.01606 (0.03331) | 0.482 | 0.630456 | | | LT_Poor : 3 - 30 | -0.1342 (0.0269) | -4.98 | **<.0001** |
| | | | anova(lmer) | | | LT | | F(1,9) = 0.2215 | 0.6492 | | | LT_Poor : 10 - 30 | -0.152 (0.0357) | -4.261 | **0.0002** |
| | | | | | | Dose | | F(3,131) = 19.34 | **1.94E-10** | | | LT_Inter : Veh - 3 | -0.0386 (0.0224) | -1.722 | 0.3166 |
| | | | | | | ang_sep | | F(4,130) =8.98 | **1.97E-06** | | | LT_Inter : Veh - 10 | 0.0358 (0.0213) | 1.678 | 0.3397 |
| | | | | | | Dose:LT | | F(3,131) = 2.52 | *0.0612* | | | LT_Inter : Veh - 30 | -0.1098 (0.0196) | -5.591 | **<.0001** |
| | | | | | | | | | | | | LT_Inter : 3 - 10 | 0.0743 (0.026) | 2.858 | **0.0253** |
| | | | | | | | | | | | | LT_Inter : 3 - 30 | -0.0712 (0.0238) | -2.992 | **0.0172** |
| | | | | | | | | | | | | LT_Inter : 10 - 30 | -0.1455 (0.0214) | -6.796 | **<.0001** |
| | | | | | | | | | | | Dose*LT | Veh Poor Learner - 3 Poor Learner | 0.0404 (0.0188) | 2.148 | 0.3905 |
| | | | | | | | | | | | | Veh Poor Learner - 10 Poor Learner | 0.0583 (0.0269) | 2.164 | 0.3806 |
| | | | | | | | | | | | | Veh Poor Learner - 30 Poor Learner | -0.0937 (0.0269) | -3.479 | **0.0153** |
| | | | | | | | | | | | | Veh Poor Learner - Veh Inter Learner | -0.0128 (0.091) | -0.141 | 1 |
| | | | | | | | | | | | | Veh Poor Learner - 3 Inter Learner | -0.0514 (0.0919) | -0.559 | 0.9987 |
| | | | | | | | | | | | | Veh Poor Learner - 10 Inter Learner | 0.0229 (0.0913) | 0.251 | 1 |
| | | | | | | | | | | | | Veh Poor Learner - 30 Inter Learner | -0.1226 (0.091) | -1.347 | 0.8606 |
| | | | | | | | | | | | | 3 Poor Learner - 10 Poor Learner | 0.0178 (0.0269) | 0.662 | 0.9978 |
| | | | | | | | | | | | | 3 Poor Learner - 30 Poor Learner | -0.1342 (0.0269) | -4.98 | **0.0001** |
| | | | | | | | | | | | | 3 Poor Learner - Veh Inter Learner | -0.0533 (0.091) | -0.586 | 0.9983 |
| | | | | | | | | | | | | 3 Poor Learner - 3 Inter Learner | -0.0918 (0.0919) | -1 | 0.9641 |
| | | | | | | | | | | | | 3 Poor Learner - 10 Inter Learner | -0.0175 (0.0913) | -0.192 | 1 |
| | | | | | | | | | | | | 3 Poor Learner - 30 Inter Learner | -0.163 (0.091) | -1.791 | 0.6388 |
| | | | | | | | | | | | | 10 Poor Learner - 30 Poor Learner | -0.152 (0.0357) | -4.261 | **0.001** |
| | | | | | | | | | | | | 10 Poor Learner - Veh Inter Learner | -0.0711 (0.093) | -0.764 | 0.9917 |
| | | | | | | | | | | | | 10 Poor Learner - 3 Inter Learner | -0.1097 (0.0939) | -1.168 | 0.9255 |
| | | | | | | | | | | | | 10 Poor Learner - 10 Inter Learner | -0.0353 (0.0933) | -0.379 | 0.9999 |
| | | | | | | | | | | | | 10 Poor Learner - 30 Inter Learner | -0.1809 (0.0931) | -1.944 | 0.5535 |
| | | | | | | | | | | | | 30 Poor Learner - Veh Inter Learner | 0.0809 (0.093) | 0.87 | 0.9829 |
| | | | | | | | | | | | | 30 Poor Learner - 3 Inter Learner | 0.0423 (0.0939) | 0.451 | 0.9997 |
| | | | | | | | | | | | | 30 Poor Learner - 10 Inter Learner | 0.1167 (0.0933) | 1.25 | 0.8989 |
| | | | | | | | | | | | | 30 Poor Learner - 30 Inter Learner | -0.0289 (0.0931) | -0.31 | 1 |
| | | | | | | | | | | | | Veh Inter Learner - 3 Inter Learner | -0.0386 (0.0224) | -1.722 | 0.6731 |

| | | | | | | | | | | | | | |
|---|---|---|---|---|---|---|---|---|---|---|---|---|---|
| | | | | | | | | | | Veh Inter Learner - 10 Inter Learner | 0.0358 (0.0213) | 1.678 | 0.7016 |
| | | | | | | | | | | Veh Inter Learner - 30 Inter Learner | -0.1098 (0.0196) | -5.591 | **<.0001** |
| | | | | | | | | | | 3 Inter Learner - 10 Inter Learner | 0.0743 (0.026) | 2.858 | *0.09* |
| | | | | | | | | | | 3 Inter Learner - 30 Inter Learner | -0.0712 (0.0238) | -2.992 | *0.0636* |
| | | | | | | | | | | 10 Inter Learner - 30 Inter Learner | -0.1455 (0.0214) | -6.796 | **<.0001** |
| **S9** | Total DAPI Assay 1 Pvalb, Sst PFC only | n=7 - Poor n=6 - Inter | anova(lmer) | (1\|id) | LT | F(1,11.8) = 2.7657 | 0.1222 | **Figure S5** | Region significance means no comparisons were made between regions with specific marker co-expression | | | | |
| | | | | | Region | F(3,29.49) = 31.44 | **2.55E-09** | | | | | | |
| | | | | | LT:Region | F(3,29.49) = 0.42 | 0.7496 | | | | | | |
| | GABRA5:Actin | n=7 - Poor n=6 - Inter | anova(lmer) | | LT | F(1,10.84) = 0.0103 | 0.3632 | **C** | LT | No significant main effect *(anova(lmer))* so pairwise not performed | | | |
| | | | | | Region | F(3,27.623) = 22.0728 | 1.69E-07 | | Region | Cg1 - PrL | 0.086 (0.269) | 0.319 | 0.9885 |
| | | | | | LT:Region | F(3,27.623) = 1.7418 | 0.1816 | | | Cg1 - IL | 0.498 (0.269) | 0.2743 | 0.2743 |
| | | | | | | | | | | Cg1 - V1 | -1.422 (0.262) | -5.425 | **<.0001** |
| | | | | | | | | | | PrL - IL | 0.412 (0.264) | 1.558 0 | 0.419 |
| | | | | | | | | | | PrL - V1 | -1.508 (0.257) | -5.869 | **<.0001** |
| | | | | | | | | | | IL - V1 | -1.920 (0.257) | -7.471 | **<.0001** |
| | | | | | | | | | LT\|Region | No significant main effect *(anova(lmer))* so pairwise not performed | | | |
| | log10 Number of Pvalb - GABRA5+ cells | n=7 - Poor n=6 - Inter | anova(lmer) | (1\|id) | LT | F(1,11.11) = 0.4533 | 0.51455 | **F** | LT | No significant main effect *(anova(lmer))* so pairwise not performed | | | |
| | | | | | Region | F(3,28.25) = 16.9745 | 1.69E-06 | | Region | No significant main effect *(anova(lmer))* so pairwise not performed | | | |
| | | | | | LT:Region | F(3,28.25) = 2.3512 | *0.0935* | | LT\|Region | Cg1, Poor-Good | -0.204 (0.354) | -0.576 | 0.5708 |
| | | | | | | | | | | PrL, Poor-Good | -0.113 (0.354) | -0.32 | 0.7524 |
| | | | | | | | | | | IL, Poor-Good | 0.262 (0.354) | 0.741 | 0.4674 |
| | | | | | | | | | | V1, Poor-Good | -0.753 (0.417) | -1.806 | *0.081* |
| | log10 Number of Sst - GABRA5+ cells | n=7 - Poor n=6 - Inter | anova(lmer) | (1\|id) | LT | F(1,11.114) = 0.4305 | 0.52508 | | LT | No significant main effect *(anova(lmer))* so pairwise not performed | | | |
| | | | | | Region | F(3,28.52) = 3.3726 | **0.03199** | | Region | Explained by total DAPI variation, no pairwise comparisons performed | | | |
| | | | | | LT:Region | F(3, 28.52) = 0..2738 | 0.8438 | | LT\|Region | No significant main effect *(anova(lmer))* so pairwise not performed | | | |
| | log10 Pvalb + GABRA5 mRNA copies | n=7 - Poor n=6 - Inter | anova(lmer) | (1\|id) | LT | F(1,10.99) = 0.1810 | 0.6787 | **G** | LT | No significant main effect *(anova(lmer))* so pairwise not performed | | | |
| | | | | | Region | F(3,28.1) = 15.8262 | **0.0000** | | Region | Explained by total DAPI variation, no pairwise comparisons performed | | | |
| | | | | | LT:Region | F(3, 28.1) = 1.6862 | 0.1925 | | LT\|Region | No significant main effect *(anova(lmer))* so pairwise not performed | | | |
| | log10 Sst + GABRA5 mRNA copies | n=7 - Poor n=6 - Inter | anova(lmer) | (1\|id) | LT | F(1,11) = 0.2737 | 0.6112 | | LT | No significant main effect *(anova(lmer))* so pairwise not performed | | | |
| | | | | | Region | F(3,28.2) = 2.3873 | *0.0900* | | Region | Explained by total DAPI variation, no pairwise comparisons performed | | | |
| | | | | | LT:Region | F(3, 28.2) = 0.6914 | 0.5649 | | LT\|Region | No significant main effect *(anova(lmer))* so pairwise not performed | | | |
